# Aspartyl protease in the secretome of honey bee trypanosomatid parasite is essential for the efficient infection of host

**DOI:** 10.1101/2023.01.23.525124

**Authors:** Xuye Yuan, Jianying Sun, Tatsuhiko Kadowaki

**Author notes:** Corresponding author: Tatsuhiko Kadowaki, Department of Biological Sciences, Xi’an Jiaotong-Liverpool University, 111 Ren’ai Road, Suzhou Dushu Lake Higher Education Town, Jiangsu Province 215123, China.

## Abstract

Exoproteome represents the proteome consisting of all secreted proteins and proteins derived from the cell surface and lysed cell. The exoproteome of the trypanosomatid parasite should interact with the host cells and the associated microbiota; however, the roles of infecting insect hosts are not yet understood. To uncover the functions of exoproteome, we identified the exoproteome of honey bee trypanosomatid parasite, *Lotmaria passim*, and found that approximately 28 % are shared with that common between *Leishmania* spp. It demonstrates a core exoproteome with conserved functions exists in the Leishmaniinae lineage. The bioinformatic characterization suggests that *L. passim* exoproteome may interact with the host and its microbiota as well as their metabolites. Deletion of genes encoding two secretome proteins revealed that an aspartyl protease but not chitinase affects the development of *L. passim* under the culture condition and is necessary for the efficient infection in the honey bee gut. Our results demonstrate that the exoproteome represents a resource to uncover the mechanisms of trypanosomatid parasites to infect the insect host by interacting with the gut environment.

## Introduction

The protein content outside a cell in either a conditioned medium or the extracellular matrix is known as the exoproteome (Esteves *et al*., 2022). The exoproteome contains actively secreted proteins and non-secreted proteins derived from cell surface proteins and lysed cells. The part of the exoproteome is a secretome composed of proteins that are conventionally and non-conventionally secreted (Hathout, 2007). In the non-conventionally secreted proteins, the proteins associated with extracellular vesicles (EVs) or exosomes have been extensively studied with various organisms to date (Denzer *et al*., 2000; Hessvik and Llorente, 2018). Exosomes carry proteins, nucleic acids, and lipids and mediate intercellular communication in both short and long distances. The contents of exosomes could vary based on the cell type and physiological conditions and have, for example, anti-inflammatory and anti-microbial functions (Keshtkar *et al*., 2021).

Exoproteomes have also been characterized with trypanosomatid parasites, including *Leishmania* spp. to enable the identification of proteins involved in parasite survival, pathogenesis, and other biologically relevant processes. Pissarra *et al*., identified exoproteomes of seven *Leishmania* species and found the shared core set of proteins as well as the species-specific proteins (Pissarra *et al*., 2022). These results suggest the exoproteome has both conserved functions to adapt to the host and specific roles related to species-specific pathogenesis. *Leishmania major* was shown to secrete exosomes under both culture condition and sandfly midgut and exacerbate the disease pathology through over induction of inflammatory cytokines in a mammalian host (Atayde *et al*., 2015). However, the roles of exosomes for infection in the insect vector have never been addressed. One of the conventionally secreted proteins, chitinase, was tested with *Leishmania mexicana* for the infection in a sand fly. The ectopic expression increased survival and growth in sand flies by facilitating escape from the peritrophic membrane (PM) and colonization of the stomodeal valve (Rogers *et al*., 2008). PM was shown to protect a gut parasite like *Leishmania* spp. from digestive enzymes in the midgut (Pimenta *et al*., 1997) but also act as a physical barrier for further development (Sacks and Kamhawi, 2001).

*Lotmaria passim* is the most prevalent trypanosomatid parasite infecting honey bees across the globe (Regan *et al*., 2018; Stevanovic *et al*., 2016; Arismendi *et al*., 2016; Ravoet *et al*., 2015). It specifically colonizes in honey bee hindgut, affects the host physiology, and may associate with winter colony loss (Ravoet *et al*., 2013; Liu *et al*., 2020). The related species, *Crithidia bombi* (Schwarz *et al*., 2015) infects bumble bee gut and imposes various negative impacts on the host performance (Brown, Schmid - Hempel and Schmid Hempel, 2003; Gegear, Otterstatter and Thomson, 2006). Both species are monoxenous parasites infecting only bees as the host; however, the mechanisms to establish and maintain the infection are not fully understood. To identify *L. passim* factors interacting with the honey bee hindgut environment, we first characterized the exoproteome of promastigotes under culture condition. We then studied two conventionally secreted proteins, aspartyl protease (LpAsp) and chitinase (LpCht) as the models to uncover the impact of secretome on the infection of *L. passim* in the honey bees. We expected these enzymes are not essential for the growth of *L. passim* under culture condition so that parasites with the loss-of-functions should be viable. However, the gene mutation could have a significant impact on the infection in the honey bees. Our study will give insight into how a gut parasite interacts with the insect host by the secretome.

## Materials and Methods

### Characterization of *L. passim* exoproteome

We cultured *L. passim* promastigotes in 30 mL of Insectagro DS2 serum-free/protein-free medium (Corning). Parasites were grown at 28.5 °C in a culture flask until the early stationary phase (2 × 10^7^/mL). The viability of parasites was more than 94 % based on propidium iodide (SIGMA) staining followed by the flow cytometry analysis. We collected the conditioned culture medium by centrifugation at 1000 × g for 10 min and filtration using a 0.2 μm pore-size filter (Millipore). The filtered medium was then concentrated by ultrafiltration using Vivaspin 20 (3 kDa cut-off, Sartorius). Protein concentration was determined using a BCA protein assay kit (Beyotime) and diluted to 1 mg/mL with 100 mM ammonium bicarbonate. Proteins (50 μg) were first reduced with 10 mM dithiothreitol at 95 °C for 5 min followed by alkylation with 55 mM iodoacetamide at room temperature for 20 min under the dark. Trypsin (1 μg) was added for digestion at 37 °C overnight, and then the protein sample was desalted using a spin column (MonoSpin C18) and dried up. The proteins in 0.1 % formic acid (2 μg at 0.1 μg/μL) were applied to Easy-nLC1000 coupled to LTQ Orbitrap Elite mass spectrometer (Thermo Fisher). It was equipped with a nanoelectrospray source and operated in data-dependent acquisition (DDA) mode with the following settings: spray voltage 2,000 V, s-lens RF level 60 %, capillary temperature 275 °C, scans 300-2,000 m/z. Peptides were separated by a 15 cm analytical RSLC column (AcclaimTM PepMapTM 100 C18 2 μm pore size, 150 mm length, 50 μm i.d.) with the mobile phase consisting of 0.1 % formic acid in water (A) and of 0.1 % formic acid in acetonitrile (B) by the following gradient elution: 0-2 min (95 % A-5 % B)→152 min (70 % A-30 % B)→162 min (10 % A-90 % B)→172 min (10 % A-90 % B)→182 min (95 % A-5 % B)→187 min (95 % A-5 % B) at a flow rate of 300 nL/min. The ten most intense ions from the full scan were selected for tandem mass spectrometry. The normalized collision energy was 35 V and the default charge state was 2 in HCD mode. Scans were collected in a positive polarity mode. The sample was analyzed using MaxQuant (Version: 2.2.0.0) and the parameters were set at Fragment tolerance: 20 ppm (monoisotopic), Fixed modifications: +57 on C (carbamidomethyl), Variable modifications: +16 on M (oxidation), +42 on peptide N-terminal (acetyl), Database: *L. passim* annotated proteins (Liu *et al*., 2020), Digestion enzyme: trypsin, and Max missed cleavages: 2.

We used eggNOG-mapper v2 (Cantalapiedra *et al*., 2021) for functional annotation of *L. passim* proteins. We next mapped the annotated *L. passim* proteins to Kyoto Encyclopedia of Genes and Genomes (KEGG) (Kanehisa *et al*., 2017) Orthology database to build the annotation databases of Gene Ontology (GO) and KEGG pathway using the R package AnnotationForge (Carlson and Pagès, 2022). Finally, clusterProfiler (Yu *et al*., 2012) of the R software package was used to perform GO and KEGG pathway enrichment analyses. We compared 193 *L. passim* exoproteome proteins with 306 exoproteome proteins common between seven *Leishmania* spp (Pissarra *et al*., 2022) using reciprocal BLASTP and 67 protein pairs showed the e-value lower than 6E-03. Among them, 55 were considered homologs with an e-value lower than 1E-50. We used the proteome annotation pipeline from the STRING database (https://string-db.org/) (Szklarczyk *et al*., 2023) to predict the protein-protein interaction (PPI) for *L. passim* annotated proteins. We then constructed the PPI network for 193 *L. passim* exoproteome proteins based on the prediction and visualized the PPI network using Cytoscape software (https://cytoscape.org/) (Shannon *et al*., 2003).

### Identification of *L. passim* proteins potentially secreted by conventional pathway

We first identified 301 proteins with N-terminal signal peptides (SPs) by screening 9339 annotated *L. passim* proteins by SignalP v5.0 (Almagro Armenteros *et al*., 2019). Next, we used TMHMM v2.0 (Krogh *et al*., 2001) to filter the proteins with transmembrane domains and finally identified 157 conventionally secreted protein candidates after the manual inspection.

### Testing secretion of LpAsp-GFP and LpCht-GFP fusion proteins

To construct plasmid DNA expressing LpAsp-GFP or LpCht-GFP fusion protein, we first amplified *LpAsp* (Lp_000442900.1) and *LpCht* (Lp_160010600.1) DNAs by PCR using KOD-FX DNA polymerase (TOYOBO), *L. passim* genomic DNA, and the following primer pairs; LpAsp-5 and LpAsp-3 as well as LpCht-5 and LpCht-3. The PCR products were digested with XbaI, gel purified, and then cloned in the XbaI site of pTrex-n-eGFP (Peng *et al*., 2014). We collected actively growing *L. passim* (4 × 10^7^), washed twice with 5 mL PBS, and then resuspended in 0.4 mL of Cytomix buffer without EDTA (20 mM KCl, 0.15 mM CaCl_2_, 10 mM K_2_HPO_4_, 25 mM HEPES and 5 mM MgCl_2_, pH 7.6) (Van den Hoff, Moorman and Lamers, 1992; Ngo *et al*., 1998). The parasites were electroporated twice (1 min interval) with 10 μg of plasmid DNA expressing LpAsp-GFP or LpCht-GFP as well as pTrex-n-eGFP using a Gene Pulser X cell electroporator (Bio-Rad) and cuvette (2 mm gap). We set the voltage, capacitance, and resistance at 1.5 kV, 25 μF, and infinity, respectively. The electroporated parasites were cultured in 4 mL of modified FP-FB medium (Salathé *et al*., 2012), and then G418 (200 μg/mL, SIGMA) was added after 24 h to select the G418-resistant clones.

To visualize GFP, we washed live *L. passim* expressing GFP, LpAsp-GFP, or LpCht-GFP three times with PBS, and then mounted them on poly-L-lysine coated slide glass. We started culturing the above *L. passim* in 4 mL culture medium with G418 at 10^4^/mL and then collected the parasites as well as a conditioned medium after 4 days. GFP and the fusion proteins were immunoprecipitated by adding GFP-Trap Agarose (ChromoTek) for 5 h at 4 □, and then the beads were washed three times with 1 mL washing buffer (10 mM Tris-HCl, pH7.5, 150 mM NaCl, 0.05 % NP-40, 0.5 mM EDTA). We finally suspended the beads and the parasites collected above (4 × 10^7^) with 40 and 200 μL of SDS-PAGE sample buffer (2 % SDS, 10 % glycerol, 10 % β-mercaptoethanol, 0.25 % bromophenol blue, 50 mM Tris-HCl, pH 6.8), respectively. After heating the samples at 95 □ for 5 min and centrifugation, 15 μL supernatants were applied to 12 % SDS-PAGE and the proteins were transferred to a nitrocellulose membrane (Pall Life Sciences). The membrane was then blocked with PBST (PBS with 0.1 % Tween-20) containing 5 % BSA at room temperature for 30 min followed by incubating with 1000-fold diluted anti-GFP antibody (proteintech) at 4 □ overnight. The membrane was washed five times with PBST (5 min each) and then incubated with 10,000-fold diluted IRDye^®^ 680RD donkey anti-rabbit IgG (H+L) (LI-COR Biosciences) in PBST containing 5 % skim milk at room temperature for 2 h. The membrane was washed as above and then visualized using Odyssey Imaging System (LI-COR Biosciences).

### Deletion of *LpAsp* and *LpCht* genes by CRISPR

To delete *LpAsp* and *LpCht* genes, we first designed the gRNA sequences using a custom gRNA design tool (http://grna.ctegd.uga.edu) (Peng and Tarleton, 2015). Two complementary oligonucleotides (0.1 nmole each) corresponding to these sgRNA sequences (LpAsp_For and LpAsp_Rev as well as LpCht_For and LpCht_Rev) were phosphorylated by T4 polynucleotide kinase (TAKARA) followed by annealing and cloning into BbsI digested pSPneogRNAH vector (Zhang and Matlashewski, 2015). We electroporated *L. passim* expressing Cas9 (Liu, Lei and Kadowaki, 2019) with 10 μg of plasmid DNA constructed above and selected the transformants by blasticidin (50 μg/mL, Macklin) and G418 to establish the parasite expressing both Cas9 and *LpAsp* gRNA or *LpCht* gRNA.

We constructed the donor DNA for *LpAsp* gene by fusion PCR of three DNA fragments: 5’UTR (515 bp, LpAsp5’UTR-F and LpAsp5’UTR-R), the open reading frame (ORF) of *Hygromycin B phosphotransferase* (*Hph*) gene derived from pCsV1300 (Park *et al*., 2013) (1026 bp, LpAspHph-F and LpAspHph-R), and 3’ half of *LpAsp* ORF (460 bp, LpAsp3’ORF-F and LpAsp3’ORF-R). Similarly, the donor DNA for *LpCht* was prepared as above with the 5’UTR (574 bp, LpCht5’UTR-F and LpCht5’UTR-R), *Hph* ORF (1026 bp, LpChtHph-F and LpChtHph-R), and the 3’UTR (589 bp, LpCht3’ORF-F and LpCht3’ORF-R). The fusion PCR products were cloned into the EcoRV site of pBluescript II SK(+) and the linearized plasmid DNA (10 μg) by HindIII was used for electroporation of *L. passim* expressing both Cas9 and *LpAsp* gRNA or *LpCht* gRNA as described above.

After electroporation, *L. passim* resistant to blasticidin, G418, and hygromycin (150 μg/mL, SIGMA) were selected and the single parasite was cloned by serial dilutions in a 96-well plate. We initially determined the genotype of each clone through the detection of 5’ wild-type (WT) and knock-out (KO) alleles for *LpAsp* and *LpCht* by PCR. After identifying the heterozygous (+/-) and homozygous (-/-) KO clones, their 5’WT (LpAsp5’UTR-Outer-F and LpAsp-72R as well as LpCht5’UTR-Outer-F and LpCht-108R), 5’KO (LpAsp5’UTR-Outer-F and Hyg-159R as well as LpCht5’UTR-Outer-F and Hyg-159R), 3’WT (LpAsp-555F and LpAsp3’UTR-Outer-R as well as LpCht-756F and LpCht3’UTR-Outer-R), and 3’KO (Hyg-846F and LpAsp3’UTR-Outer-R as well as Hyg-846F and LpCht3’UTR-Outer-R) alleles were confirmed by PCR using the specific primer sets.

### Detection of *LpAsp* and *LpCht* mRNAs by RT-PCR

Total RNA was extracted from wild-type, *LpAsp*, and *LpCht* heterozygous and homozygous mutant parasites using TRIzol reagent (SIGMA) and treated with 1 unit of RNase-free DNase (Promega) at 37 °C for 30 min. Total RNA (0.2 μg) was reverse transcribed using ReverTra Ace (TOYOBO) and random primer followed by PCR with KOD-FX DNA polymerase. For Figure 3C, a primer corresponding to *L. passim* splice leader sequence (LpSL-F) was used as the forward primer. As the reverse primers, LpAsp-260R and LpGAPDH-R were used. For Figure 3D, LpCht-756F and LpCht-1248R as well as LpGAPDH-F and LpGAPDH-R were used.

**Figure 1.**
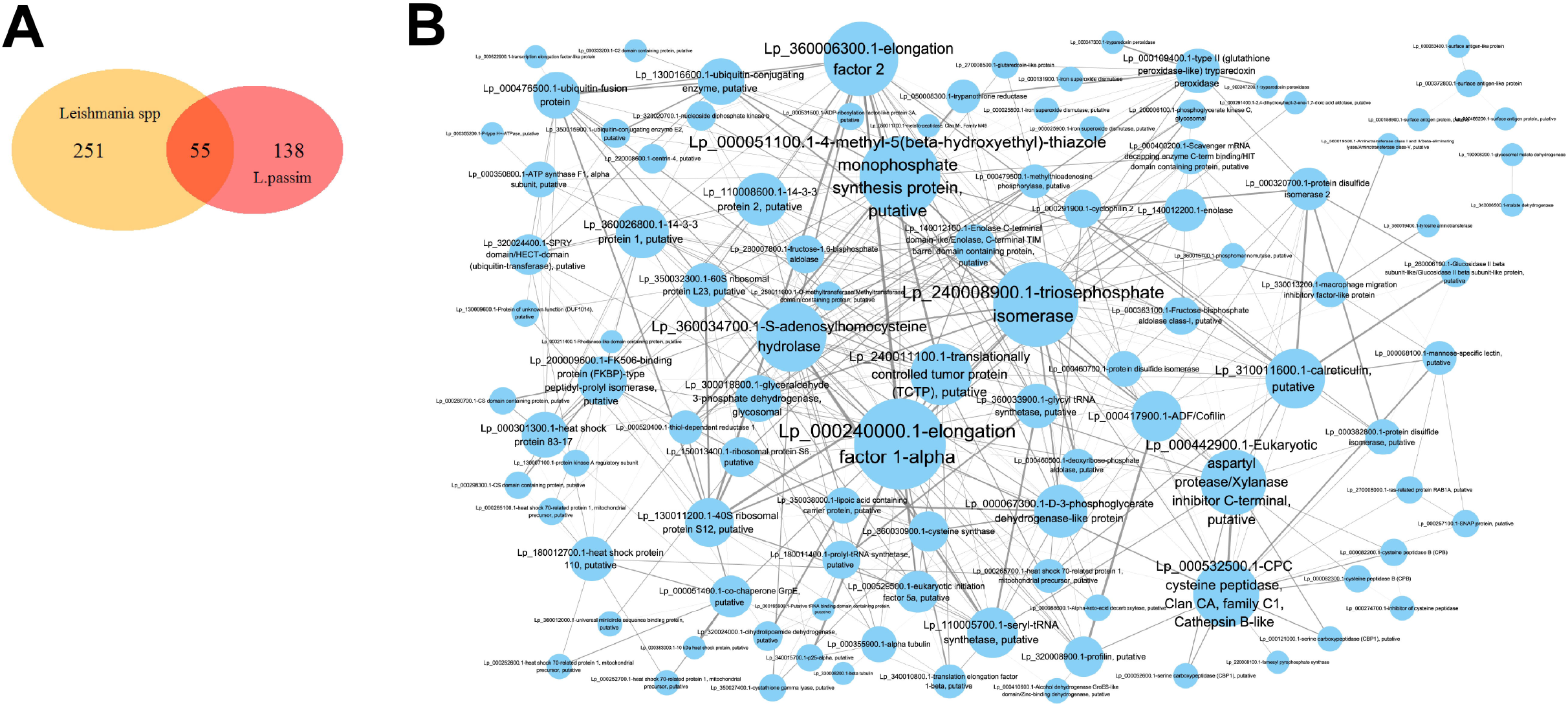
Characterization of *Lotmaria passim* exoproteome. (A) A Venn diagram to indicate 55 proteins are shared between 193 *L. passim* exoproteome proteins and 306 exoproteome proteins common with seven *Leishmania* spp. (Pissarra *et al*., 2022). (B) Protein-protein interaction map of *L. passim* exoproteome predicted by STRING. The size of the circle and the thickness of the line are proportional to the number and confidence of interactions, respectively.

**Figure 2.**
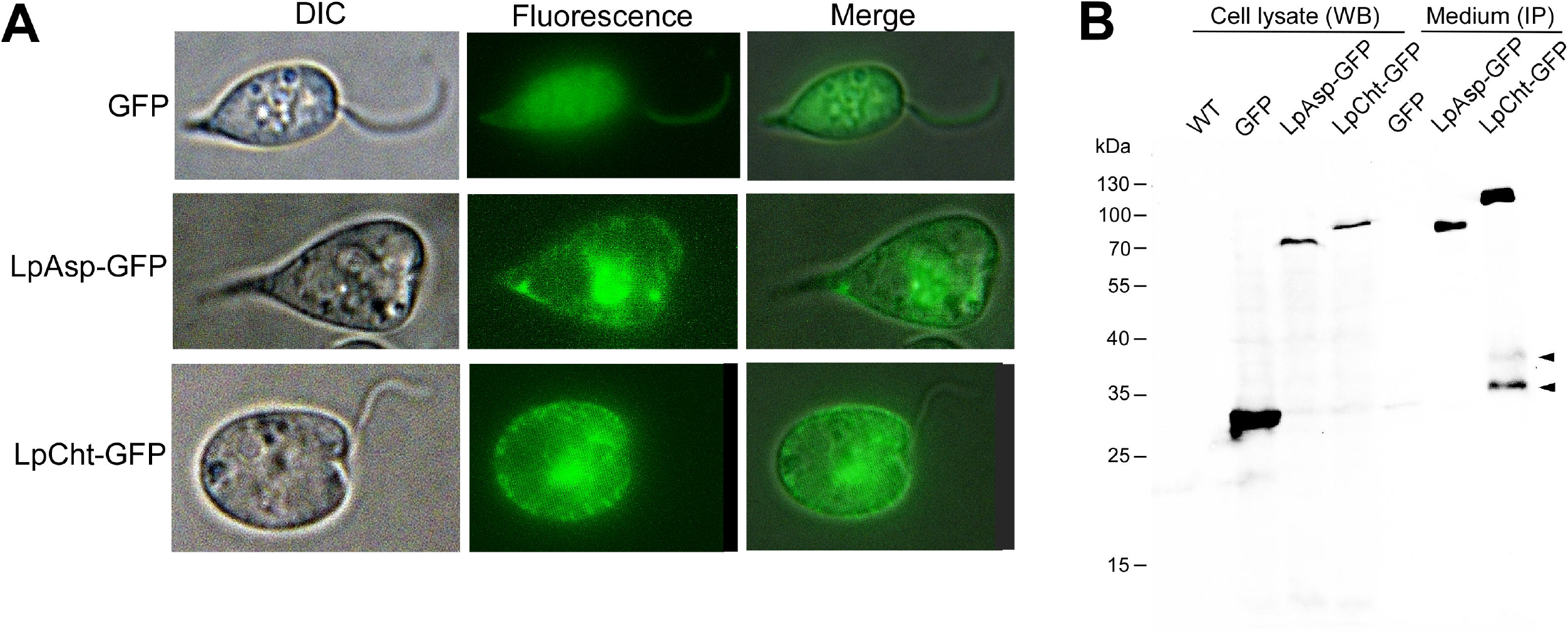
Secretion of LpAsp and LpCht. (A) *L. passim* expressing GFP, LpAsp-GFP, or LpCht-GFP fusion was viewed by visible (DIC) or fluorescence (Fluorescence) light. The merged images are also shown (Merge). The fluorescence signals for LpAsp-GFP and LpCht-GFP were intensified by longer exposure time than GFP. (B) The amounts of intracellular GFP, LpAsp-GFP, and LpCht-GFP were measured by western blot (WB) of lysates prepared with an equal number of parasites (Cell lysate). Wild-type *L. passim* was used as a control (WT). The amounts of the extracellular proteins were measured by immunoprecipitation (IP) of the equal volume of conditioned medium (Medium) followed by WB. Two cleaved LpCht-GFP proteins are indicated by arrowheads. The molecular weight (kDa) of the protein marker is indicated on the left.

**Figure 3.**
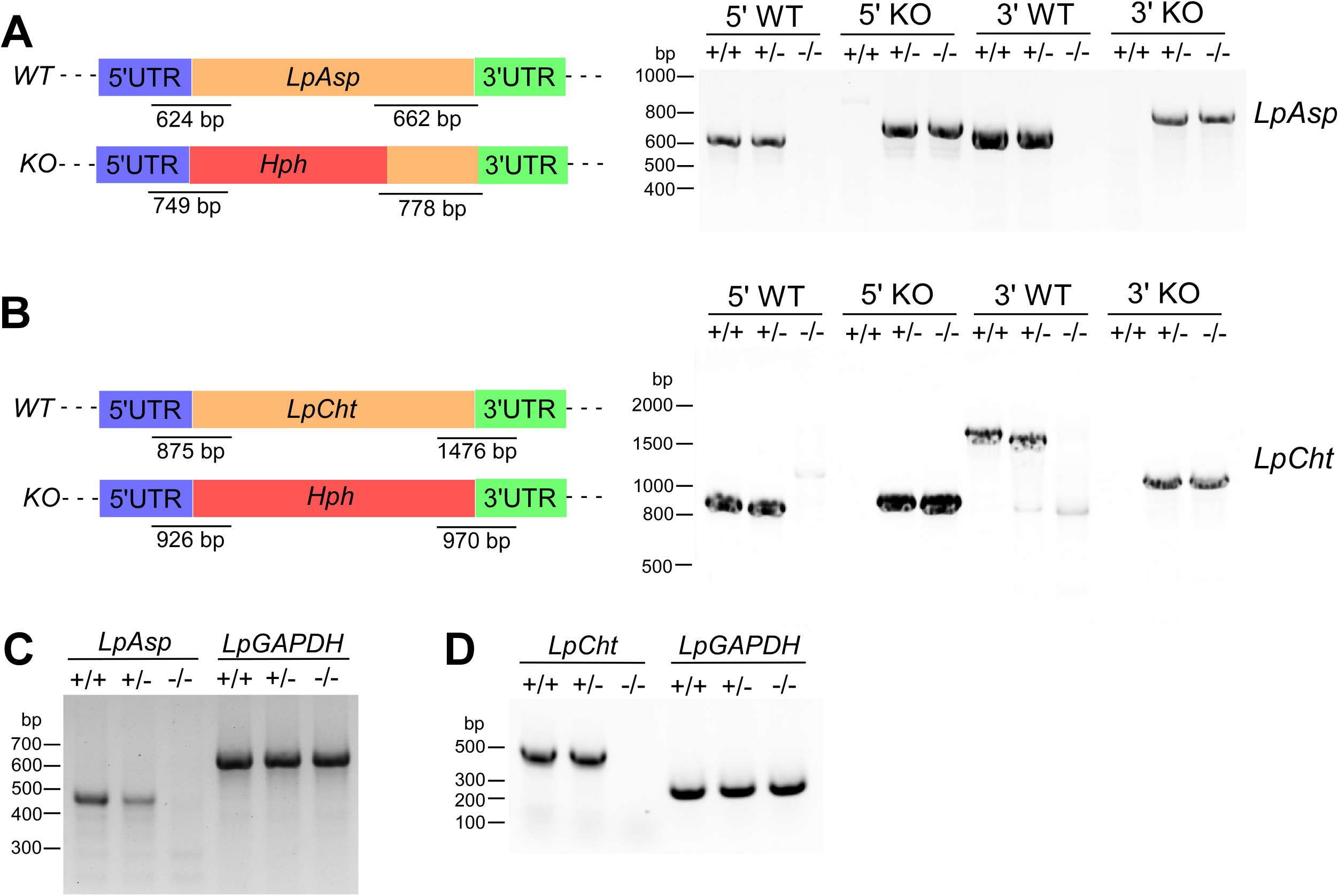
Deletion of *LpAsp* and *LpCht* genes by CRISPR. Schematic representation of wild-type (*WT*) and deleted (*KO*) alleles of *LpAsp* (A) and *LpCht* (B) generated by CRISPR/Cas9-induced homology-directed repair. 5’ and 3’ untranslated regions (UTRs), open reading frame (ORF), and hygromycin resistance gene (*Hph*) are shown in blue, green, yellow, and red, respectively. The expected sizes of PCR products to detect *WT* and *KO* alleles (not in scale) are also shown for each gene. Genomic DNAs of wild-type *L. passim* (+/+), heterozygous (+/-), and homozygous (-/-) mutants of *LpAsp* and *LpCht* were analyzed by PCR to detect 5’*WT*, 5’KO, 3’WT, and 3’KO alleles. Sizes of the DNA molecular weight markers are shown on the left. (C) Detection of *LpAsp* and *LpGAPDH* mRNAs in *LpAsp* heterozygous (+/-) and homozygous (-/-) mutants together with wild-type *L. passim* (+/+) by RT-PCR. A primer corresponding to *L. passim* splice leader sequence was used as the forward primer. Sizes of the DNA molecular weight markers are shown on the left. (D) Detection of *LpCht* and *LpGAPDH* mRNAs in *LpCht* heterozygous (+/-) and homozygous (-/-) mutants together with wild-type *L. passim* (+/+) by RT-PCR. Sizes of the DNA molecular weight markers are shown on the left.

### Characterization of *LpAsp and LpCht* mutant parasites

We inoculated wild-type, *LpAsp*, and *LpCht* homozygous mutant parasites in the culture medium at 10^4^/mL, and then counted the number of parasites every day using a hemocytometer. The images of cultured parasites were captured after 5 days. The number of rosettes present in the three different images was counted with the individual parasites.

To infect honey bees with *L. passim*, we collected the parasites during the logarithmic growth phase and washed them once with PBS followed by suspension in sterile 10 % sucrose/PBS at 5 × 10^4^ /μL. Newly emerged honey bee workers were collected and then starved for 2-3 h. Twenty honey bees were fed with 2 μL of above 10 % sucrose solution (10^5^ parasites in total) containing either wild-type, *LpAsp*, or *LpCht* homozygous mutant. The infected honey bees were maintained in metal cages at 33 °C for 14 days and then frozen at −80 °C. These experiments were repeated three times. We sampled eight honey bees from each of the above three experiments, and thus analyzed 24 honey bees in total infected with either wild-type, *LpAsp*, or *LpCht* homozygous mutant. Genomic DNAs were extracted from the whole abdomens of individual bees using DNAzol® reagent (Thermo Fisher). We quantified *L. passim* in the infected honey bee by qPCR using LpITS2-F and LpITS2-R primers which correspond to the part of internal transcript spacer region 2 (ITS2) in the *ribosomal RNA* gene. Honey bee *AmHsTRPA* was used as the internal reference using AmHsTRPA-F and AmHsTRPA-R primers (Liu *et al*., 2020). The relative abundances of *L. passim* in the individual honey bees were calculated by the ΔCt method and the Brunner-Munzel test was used for the statistical analysis. All of the above primers are listed in Supplementary file 1.

## Results and Discussion

### Identification and characterization of *L. passim* exoproteome

We first identified the exoproteome of *L. passim* promastigotes by mass spectrometry (MS) analysis of the conditioned culture medium and the list of proteins (193) is shown in Supplementary file 2. The KEGG pathway enrichment analysis shows that “Exosome” is most significantly enriched with the exoproteome, indicating that we successfully recovered proteins associated with EVs (Table 1). We also found that three pathways related to bacterial infection, “Pathogenic *Escherichia coli* infection”, *“Salmonella* infection”, and “Legionellosis” are enriched. *L. passim* may use the proteins associated with these KEGG pathways to interact with microbiota present in honey bee hindgut. The results of the GO term enrichment analysis were consistent with those of the above KEGG pathway analysis (Table 2). For example, in the Cellular Compartment category, GO terms, “extracellular region” and “biofilm matrix” are enriched. The enrichment of the GO term, “cell surface” suggests that *L. passim* exoproteome contains a significant amount of cell surface proteins shed into the culture medium. With the Molecular Function annotations, the GO term “icosanoid binding” is enriched, suggesting that the exoproteome proteins may bind with various metabolites in honey bee hindgut. In the Biological Process category, we found the enrichment of GO terms such as “positive regulation of membrane permeability”, “positive regulation of immune response”, “regulation of cell-matrix adhesion”, and “response to host defenses”. The proteins related to these GO terms may have roles to interact with the host as well as the hindgut microbiota.

**Table 1.**
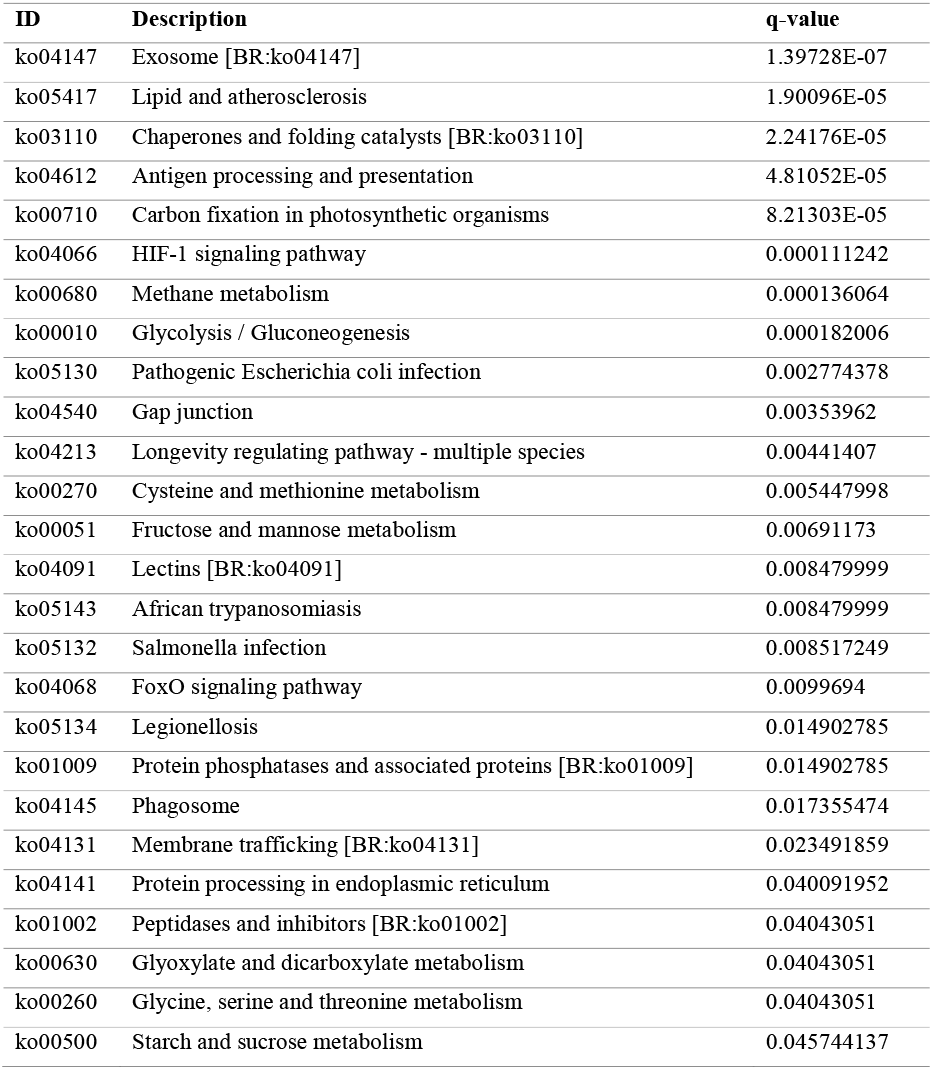
KEGG pathway enriched with *L. passim* exoproteome

**Table 2.** Representative GO terms enriched with *L. passim* exoproteome

| ONTOLOGY | ID | Description | q-value |
| --- | --- | --- | --- |
| BP | GO:0019752 | carboxylic acid metabolic process | 4.83E-07 |
| BP | GO:0006165 | nucleoside diphosphate phosphorylation | 2.88E-05 |
| BP | GO:0006096 | glycolytic process | 3.51E-05 |
| BP | GO:1905710 | positive regulation of membrane permeability | 8.20E-05 |
| BP | GO:0034599 | cellular response to oxidative stress | 0.000120144 |
| BP | GO:1904592 | positive regulation of protein refolding | 0.000120144 |
| BP | GO:0050764 | regulation of phagocytosis | 0.000764768 |
| BP | GO:0050778 | positive regulation of immune response | 0.00343486 |
| BP | GO:0001952 | regulation of cell-matrix adhesion | 0.018152884 |
| BP | GO:0052200 | response to host defenses | 0.026506521 |
| BP | GO:0048870 | cell motility | 0.032949856 |
| CC | GO:0005576 | extracellular region | 3.34E-07 |
| CC | GO:0048770 | pigment granule | 1.79E-05 |
| CC | GO:0009986 | cell surface | 2.42E-05 |
| CC | GO:0005788 | endoplasmic reticulum lumen | 7.96E-05 |
| CC | GO:0070062 | extracellular exosome | 0.000346854 |
| CC | GO:0062039 | biofilm matrix | 0.000431219 |
| MF | GO:0051082 | unfolded protein binding | 1.51E-05 |
| MF | GO:1901567 | fatty acid derivative binding | 9.28E-05 |
| MF | GO:0005504 | fatty acid binding | 0.000157262 |
| MF | GO:0031685 | adenosine receptor binding | 0.000157262 |
| MF | GO:0050542 | icosanoid binding | 0.000157262 |
| MF | GO:0016209 | antioxidant activity | 0.002016583 |

Surface antigen-like protein is the most abundant protein representing 10.9 % of the exoproteome. Meanwhile, tryparedoxin 1, GP63, mitochondrial heat shock 70-related protein 1, NADPH oxidase, and β-fructofuranosidase represent 2.3, 2.8, 3.1, 6.7, and 7.4 %, respectively. These proteins were also abundant in *Leishmania* spp. exoproteome except NADPH oxidase (Pissarra *et al*., 2022). Furthermore, we compared 193 *L. passim* exoproteome proteins and 306 exoproteome proteins shared between seven *Leishmania* spp. (Pissarra *et al*., 2022). As shown in Figure 1A, 55 proteins are common, indicating that they represent the core set of exoproteome proteins with conserved functions in the Leishmaniinae lineage (Supplementary file 3).

We further identified 157 potential proteins secreted by a conventional pathway in *L. passim* by bioinformatic approach. They contain an N-terminal SP but not any transmembrane domain (Supplementary file 4). Among them, only 30 proteins are found in *L. passim* exoproteome, suggesting that most exoproteome proteins are secreted by non-conventional pathways. However, it is also possible that the amounts of some proteins secreted by conventional pathway were not enough to be detected by the MS analysis. GO term enrichment analysis of the above 157 *L. passim* proteins demonstrated that they not only include secreted proteins but also ER and Gogi luminal proteins (Table 3). These proteins contain the N-terminal SPs as well as the amino acid sequences to retain in either ER or Golgi (Murshid and Presley, 2004).

**Table 3.** Representative GO terms enriched with 157 potential proteins secreted by conventional pathway in *L. passim*

| ONTOLOGY | ID | Description | q-value |
| --- | --- | --- | --- |
| BP | GO:0034975 | protein folding in endoplasmic reticulum | 1.57E-14 |
| BP | GO:0006457 | protein folding | 1.26E-08 |
| BP | GO:0036500 | ATF6-mediated unfolded protein response | 1.5E-07 |
| BP | GO:0061635 | regulation of protein complex stability | 6.67E-07 |
| BP | GO:0006986 | response to unfolded protein | 8.52E-07 |
| BP | GO:0031204 | post-translational protein targeting to membrane, translocation | 1.11E-06 |
| BP | GO:0071287 | cellular response to manganese ion | 1.11E-06 |
| BP | GO:0034976 | response to endoplasmic reticulum stress | 1.11E-06 |
| CC | GO:0005788 | endoplasmic reticulum lumen | 7.01E-20 |
| CC | GO:0034663 | endoplasmic reticulum chaperone complex | 2.63E-11 |
| CC | GO:0005790 | smooth endoplasmic reticulum | 1.51E-08 |
| CC | GO:0005783 | endoplasmic reticulum | 2.13E-08 |
| MF | GO:0051082 | unfolded protein binding | 4.16E-07 |
| MF | GO:0031685 | adenosine receptor binding | 5.74E-07 |

Among *L. passim* secretome, we prioritized studying of two hydrolytic enzymes, LpAsp and LpCht with N-terminal SPs by the following reasons. We previously reported that GO terms associated with proteolysis are significantly enriched with *L. passim* genes continuously up-regulated during the infection of honey bees (Liu *et al*., 2020). *L. passim* exoproteome contains multiple peptidases in which *LpAsp* mRNA was shown to increase 12-27 days after the infection (Liu *et al*., 2020). Furthermore, LpAsp is one of two proteases interacting with the large number of exoproteome proteins (Fig. 1B). LpCht is of significant interest because it would be necessary for *L. passim* to release from the PM and to modify the cuticle lining of the hindgut (Schlein, Jacobson and Shlomai, 1991).

### Secretion of LpAsp and LpCht

To confirm the secretion of LpAsp and LpCht, we transformed *L. passim* with plasmid DNA to express the GFP-tagged protein. As shown in Figure 2A, GFP is uniformly present in the cytoplasm; however, LpAsp-GFP and LpCht-GFP fusion proteins are highly enriched in the specific compartment (most likely ER) as well as the plasma membrane. The fluorescence at the plasma membrane is patchy, suggesting that protein secretion may occur in specific regions. In kinetoplastids, protein is considered to be secreted through ER, Golgi, and then Flagellar pocket (FP) (Field *et al*., 2007). The above localizations of LpAsp-GFP and LpCht-GFP in *L. passim* suggest that FP may not be the only site for protein secretion in kinetoplastids. Western blot of the cell lysates of parasites expressing GFP, LpAsp-GFP, or LpCht-GFP indicates that fewer amounts of fusion proteins are present in the parasite cells compared to GFP. In contrast, more fusion proteins can be detected in the culture medium (Fig. 2B), indicating that LpAsp and LpCht are secreted from *L. passim*. Because two small LpCht-GFP proteins are present in the culture medium, LpCht is cleaved at two sites in the C-terminus after secretion.

### Deletions of *LpAsp* and *LpCht* genes by CRISPR

To uncover the functions of LpCht and LpAsp, we deleted the entire and 1-674 bp of open reading frames of *LpCht* and *LpAsp*, respectively by replacing them with *Hph* gene using CRISPR. The gene disruption was confirmed for both *LpAsp* and *LpCht* by genomic PCR using the specific primers and the homozygous mutant parasites lacked the wild-type alleles for both 5’ and 3’ ends of the gene (Fig. 3A and B). Furthermore, *LpAsp* and *LpCht* mRNAs were also absent in the homozygous mutant parasites (Fig. 3C and D). These results indicate that LpAsp and LpCht are not essential for the viability of *L. passim* under culture condition as we expected. We also found that the growth rate of the *LpAsp* or *LpCht* mutant parasite is comparable to that of the wild-type (Fig. 4A). However, there are more rosettes (clusters of cells with their flagella toward the center of the cluster) formed at the stationary phase of culture with *LpAsp* mutant parasite relative to *LpCht* mutant and wild-type parasites (Fig. 4B and C). Rosettes were observed with *Leishmania* in the midgut of insect vectors, and specifically express surface poly-α2,8 N-acetyl neuraminic acid (PSA) and PSA containing de-N-acetyl neuraminic acid (NeuPSA). Thus, the rosettes were considered to represent *Leishmania* at the distinct stage perhaps to initiate mating (Iovannisci, Plested and Moe, 2010). There has been no evidence to show *L. passim* undergoes mating; nevertheless, loss of *LpAsp* changes the physiological/developmental state of the parasite even under the culture condition.

**Figure 4.**
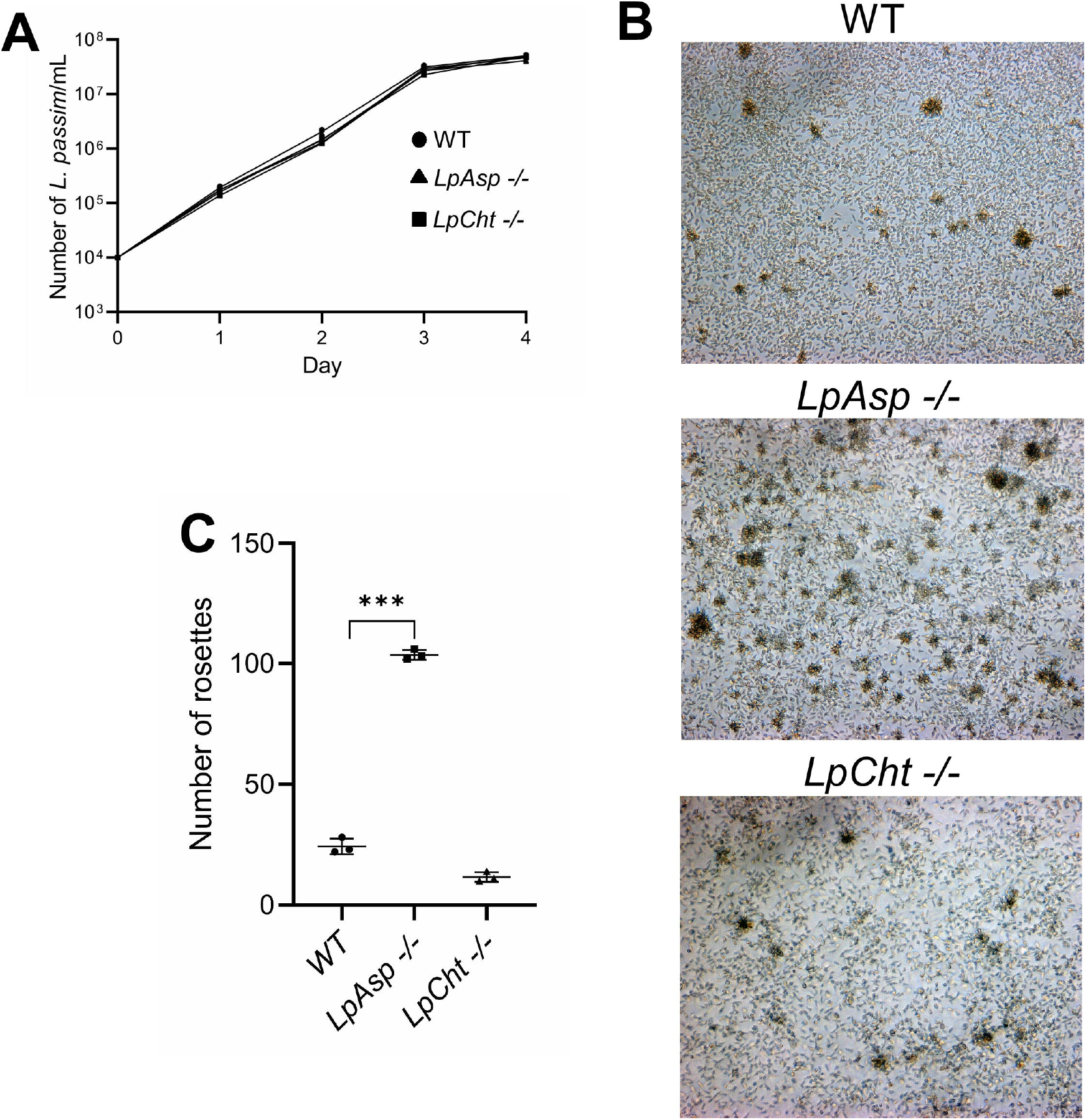
Growth and rosette formation of *LpAsp* and *LpCht* mutants under culture. (A) Growth rates of wild-type (WT, circle), *LpAsp* (triangle), and *LpCht* (square) homozygous (-/-) mutant *L. passim* in the modified FP-FB medium at 28.5 □ were monitored for four days (n = 3). (B) The images of parasites in the medium were taken five days after the culture. Rosettes of various sizes are observed. (C) The number of rosettes present in the three different areas was counted with the individual parasites (n = 3).

### LpAsp but not LpCht is necessary for the efficient infection of *L. passim* in the honey bee gut

We next tested if LpAsp and LpCht are necessary for infection of *L. passim* in honey bee hindgut. Wild-type, *LpAsp*, or *LpCht* mutant *L. passim* was infected with the honey bee, and then we quantified the relative number of parasites in the honey bee gut after maintaining in the cage for 14 days. As shown in Figures 5A and B, the infection of wild-type and *LpCht* mutant parasites was comparable; however, the number of *LpAsp* mutant parasites in the honey bee gut was less. These results demonstrate that LpAsp but not LpCht is essential for the efficient infection of *L. passim*.

**Figure 5.**
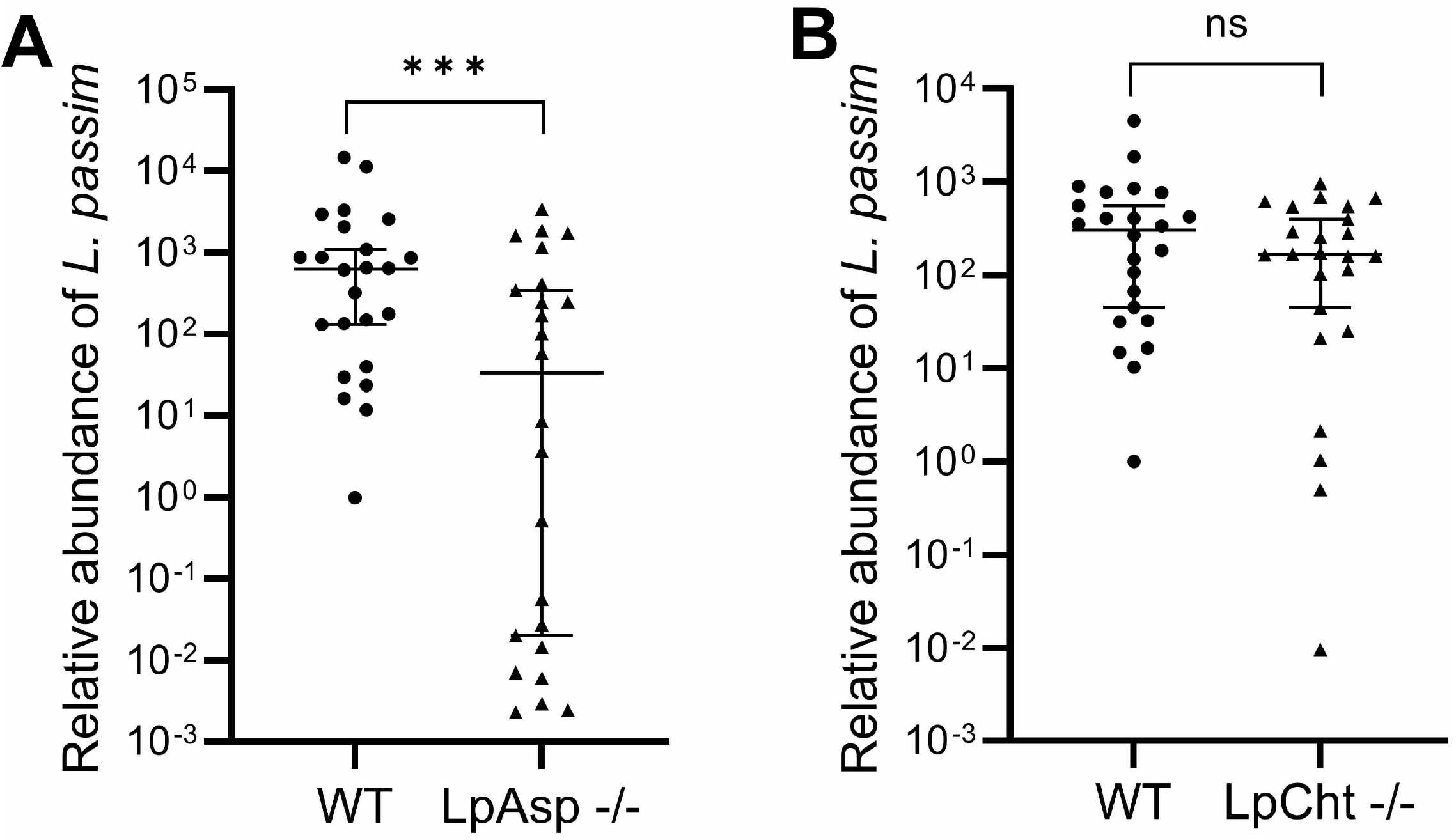
Infection of *LpAsp* and *LpCht* mutants in honey bee. The relative abundance of *L. passim* in individual honey bees (n = 24) at 14 days after the infection was compared between wild-type (WT) and *LpAsp* (A), or *LpCht* (B) homozygous mutant (-/-). One sample infected by the WT parasite was set as 1 and the Median with 95 % CI is shown. The Brunner-Munzel test was used for the statistical analysis. *** *P* < 0.0029.

*L. passim* is likely to be surrounded by PM in the midgut when ingested by a honey bee; however, the release from the PM using chitinase may not be essential. Perhaps, the release is specifically required for parasites infecting the midgut of insect vectors such as *Leishmania. L. passim* could migrate to the hindgut through the direction of gut flow in the honey bee. Attachment of *L. passim* to the epithelial cells of the hindgut may depend on chitinase to disrupt the cuticle layer. However, a recent report has shown that the cuticle layer of *L. passim-*infected hindgut appears to be intact (Buendía-Abad *et al*., 2022). Chitinase was biochemically characterized with *Leishmania* (Joshi *et al*., 2005) but the effect of loss-of-function on the infection in sandfly midgut has not been tested. Although chitinase is well conserved among trypanosomatid parasites, the role of infection in the insect host or vector is not yet fully understood.

We have found that LpAsp is required for *L. passim* to efficiently infect honey bee gut. What is the potential substrate for digestion? Protein content must be relatively low in the hindgut because the most of nutrient is adsorbed in the midgut. However, there could be some peptides/proteins derived from the gut epithelial cells and microbiota. The digested products would be necessary for the efficient infection of *L. passim*. The attachment of *L. passim* to the epithelial cell layer in the hindgut appears to require remodeling of the flagella (Buendía-Abad *et al*., 2022), and LpAsp could be required for this process. Anti-microbial peptides (AMPs) would also be the potential substrate for LpAsp. We previously reported that *AMP* mRNA expression (for example, apidaecins type 14 precursor) is up-regulated by *L. passim* infection (Liu *et al*., 2020), and thus the parasite may utilize LpAsp to inactivate AMPs. Finally, LpAsp may digest the specific *L. passim* exoproteome proteins to modulate their activities that are critical for the infection. Whether the inefficient infection of *LpAsp* mutant in honey bee hindgut is related to the enhanced formation of rosettes under the culture condition remains to be answered.

We identified the exoproteome of *L. passim* promastigotes and found that 28 % represents the core exoproteome proteins common between Leishmaniinae species. Abundant exoproteome proteins are also shared between *L. passim* and *Leishmania* spp. These results suggest that some exoproteome proteins have the conserved functions for trypanosomatid parasites to infect the gut (midgut or hindgut) of various insect species irrespective of monoxenous or dixenous life cycle. The other exoproteome proteins would have functions to infect the specific insect and mammalian host species. We also proved that two *L. passim* hydrolytic enzymes with N-terminal SPs are indeed secreted and showed at least one is involved in the development of *L. passim* under culture condition and essential for the efficient infection in the honey bees. Because the content of exoproteome is likely to change under different physiological conditions, characterizing *L. passim* exoproteome in honey bee hindgut should be of significant interest. Exoproteome represents an important resource to further understand the mechanisms of how trypanosomatid parasite infects the insect (and mammalian) host and exerts the pathogenesis.

## Acknowledgements

We thank Yuchi Ma for his contribution to study LpCht.

## Author contribution

TK conceived and designed research strategy and wrote the paper. XY and JS performed the experiments. XY, JS, and TK analyzed data.

---

**Supplementary file 1 List of primers used in this study**

**Supplementary file 2. List of 193 *L. passim* exoproteome proteins**

**Supplementary file 3. List of 55 *L. passim* exoproteome proteins shared with seven *Leishmania* spp.**

**Supplementary file 4. List of potential 157 *L. passim* proteins secreted by conventional pathway**

