## Supplementary file 1 for "Aspartyl protease in the secretome of honey bee trypanosomatid parasite is essential for the efficient infection of host"

| Primer | Sequence (5’→3’) |
| --- | --- |
| LpAsp-5 | TTTCTAGAATGCGCCGTTTTCTTGTAGCTTTC |
| LpAsp-3 | TTTCTAGACTTGGCGTAGGCAAAGCTGAGCGA |
| LpCht-5 | TTTCTAGAATGAGCTGCCTTCGTCAAAAGCGC |
| LpCht-3 | TTTCTAGAAAGGTCGCCGTCCAACGCCGCATC |
| LpAsp_For | TTGTGAGGGCGACCTCGTGCTTGG |
| LpAsp_Rev | AAACCCAAGCACGAGGTCGCCCTC |
| LpCht_For | TTGTGGTGGAGGGCCATTGTGATG |
| LpCht_Rev | AAACCATCACAATGGCCCTCCACC |
| LpAsp5’UTR-F | CAACAGCATGCTCTCACTCGCAT |
| LpAsp5’UTR-R | CTTTTTCATGATTCTGATCGATATGAAGAAGCC |
| LpAspHph-F | ATCAGAATCATGAAAAAGCCTGAACTC |
| LpAspHph-R | ACGTGGCGTCTATTTCTTTGCCCTCGG |
| LpAsp3’ORF-F | AAGAAATAGACGCCACGTGGACGGAGAGCTG |
| LpAsp3’ORF-R | TTACTTGGCGTAGGCAAAGCTG |
| LpCht5’UTR-F | GGTCTGCTGAGTGTGCAATCTG |
| LpCht5’UTR-R | CTTTTTCATTCGCGTTGAGACGACCACACCTAC |
| LpChtHph-F | TCAACGCGAATGAAAAAGCCTGAACTC |
| LpChtHph-R | CGAGCTGGGCTATTTCTTTGCCCTCGG |
| LpCht3’ORF-F | AAGAAATAGCCCAGCTCGAACTGTCGAGGCT |
| LpCht3’ORF-R | ATGCCGCTTAACAACGCATCGA |
| LpAsp5’UTR-Outer-F | GTACTGGGTTCTTCTTGCCCAGA |
| LpAsp-72R | CTTCACCTCTTGCTCGCCGACAA |
| LpAsp-555F | GGTCGAACCTCCGATGTTTTCCA |
| LpAsp3’UTR-Outer-R | AGAGTTAGTGGAGCGGGCTTACT |
| LpCht5’UTR-Outer-F | TGCTTCGTTGGAGGTGTAACCT |
| LpCht-108R | CGAAAACGGAGCTGCGCACAAAA |
| LpCht-756F | CCACTGGATGGCGTACGATC |
| LpCht3’UTR-Outer-R | CGGGATAGAAAGGGAGACGTTCTT |
| Hyg-159R | GCAGCTATTTACCCGCAGGACATA |
| Hyg-846F | CGTATATGCTCCGCATTGGTCTTG |
| LpSL-F | CTAACGCTATATAAGTATCAGTTTCTGTACTTTATTG |
| LpAsp-260R | CCGGTATCGTAAATCACCTGGAAC |
| LpGAPDH-F | TGAACGGCCACCGCATCCTG |
| LpGAPDH-R | GGGCCAGGCAGTTGGTCGTG |
| LpCht-1248R | GTCTTGACCCAGCTCCCAAATC |
| LpITS2-F | GGGTCTTTTGTGATCGGGATAA |
| LpITS2-R | CAAAAAGATGCCTAACGTGAAGAA |
| AmHsTRPA-F | TAGCGTACATGTGGTGCTGT |
| AmHsTRPA-R | GCTAGGCTCCACGTAATCCA |
