## Supplementary file 2 for "Aspartyl protease in the secretome of honey bee trypanosomatid parasite is essential for the efficient infection of host"

>Lp_000191400.1

MASYKPFYPVTPTNPKVWMDIEIGGKPAGRITMELFKDVVPKTAENFRALCTGEKGFGYAGSPFHRVIPGFMCQGGDFTAGNGTGGKSIYGNKFPDETFSGRAGKHFGPGTLSMANAGPNTNGSQFFICTANTDWLNGKHVVFGQVVEGYDVVKAMEAVGSRNGATSKPVRVSASGQL*

>Lp_000021500.1

MINEVDQDGSGTIDFPEFLTLMARKMQDSDSEEEIKEAFRVFDKDGNGFISAAELRHVMTNLGEKLTDEEGGGRDGPPGRGGGCRSNQLGGVREDDSEQVEPKSKSRKAARQ*

>Lp_050008600.1

MTSITAEKRFDNPHRDASGNYPCTEYDDAYQWRGLPPRPKTSNDNDHEDDKAPINPSMYKTTNRADYKPYDQDAYKRQQPEDDGDEDEKAPVDPNMYKTTNRDDYKPYDQDAYKRNLLDEEPSVLGEKAKGAPVVPEQSIAVSEKARKPLERESSSVARGPAVEEAVDVPVARRAPIDPNMYNSTYDDDFRPYSPQDYTAQEQEPEASPAPRHAPIDPNMYNSTYDDDFRPYSPQDYTAQEQEPAPAPAPRRAPIDPNMYKTTNRDDYKPYDQDTYRNRPTEQPLGSSSTPRGVTPKPSESPRVRRAAADPAAYDSTYKKDYGRPEPEAQEAKGGDELVPPKSSPAGQQPPPQQQQQQPSSSPAAAAQKVDQPKEAKASPRTRRAAADPAAYDSTYKKDYGRPETESKPSEEDATAKASAVPALKIPQQQQPEGKKRSSMPASARSSVRNDSAALSSRQPAQVSENKPVKKAAAQARPRPRPKANYETEYSKNYRAYSNVPTPAATPRRVPACVHHVDPNFYTTTMQTAYAAPPSPAPVRRAYTRPTVTVRHMDPSMYVSANTAAYS*

>Lp_000173500.1

MTSAKPNIDALLKPLVIGHHSMANRFVMAPLTRCRSGLEHIPNDEMVKHYSDRASMGLIICEATQIQKGYSTFGREPGIYGPAQVAGWKKVTDAVHAKGGLIYCQIHNGGRATGPCNGSGGLKVIAPSPVAIVGHDSPALFNQSGVKEHYPTPVELTKAEIEEYVQLYATAAHNAIAAGFDGVEIHGANGYLIDQFLKTSSNKRTDEYGGSIENRSRFLFEVVDAVIKKIGNERTALRISPLNSFNGQSDENPEALTKYICTELNSRKLAFLDVMRGDFFTPARGADKWAREVYTGVMFTGMGFEIDEAAKTVENGEADAVVFGVKAIANPDLVARAIAGAELNVPDVATFYTHESKGYNDYPMLACTECTAA

>Lp_320008900.1

MSWQAYVDDSLIGSGNMHSAAIIGTADGSYWAYGGNYIPQPEEVKHIQKCLNDFTLVQSSGVTIYGVKFFGLQCGSDGDSKYIFFKKGAAGGCIYTTKQAFIVAVYGNPGDASSLTQDLKKNSAHAATVNPADCNTTVKRIADYLIKLGY*

>Lp_000173800.1

MSEISYENGQPAYTGDTVVKCFKDNGNGLLFRIVNNDEHKWAFYNDTTNYNMTVKVSFGKDSKIEAIGNTKMAKDEETGEFKCELHIAPTTTEMFIQGEPNGYKISFEADPIPKA*

>Lp_000343700.1

YTHANGLDALYFAKATYCEEDAIRSWTCGTTCMQHSNFRVLAVIRNPGLSLLSYVGVEDEKQRVVVAFRGASARNQPVGSVAIATVQLDKRLGCGENCGVHAAYQSYYKTMRPLVRRYVIGALQWNPSYSVLVTGHSMGGGLSLLAAPDLQTQIDRYGFSPRPLVHLYTFGALRVSNKAYSAWAVNLLSNAAHFRVTHGDDPIPRLPPLPSLFGYVHVPQEIYYPGNDNVFVVCDDSVSGESKSCNNAAKNRDNNDHLHYLG

>Lp_150013400.1

MTGEISEKAFPLATDRLSQTILDLVQEASNAKMVKKGANEATKALNRGIADLIVLAGDTNPIEILLHLPLLCEDKNVPYVFVPSKTALGRASQVSRNVVALAILQGENSPVAAKVQAVKLEIERLL*

>Lp_180012700.1

MSVFGVDFGNLNSTVAITRYGGVDIVTNEVSKRETTTIVSFLDEERFIGEQGLDRYVRNAQNTVFLIKRFIGMRMDDPQLEIERKFLTCGVKGDDEGRLMFSVTYCDEEKCFYPEQVLAMMLQRLRTYVNEAATTDPRVKADVRDFVITVPSYYTAEQRRLMYQASEVAGLHCMSLINETTASAVDYGIFRGASLQELEADGQVVGILDIGYGATDFSVCKFWRGNCKILARTFDRNNGTRDCDYCLYQHMVNEVKSRYKVDVSENKRARLRLLQACERLKYLLSANQSAPLNVENLMDIDVNIPAFDRGTMEELCRGVLDRVRAVIERGFVEAGVTREHFHSIEMIGGGCRIPMMKRLVEDVLGRTPSFTLNASETTARGCAIVAAMLSPKFQVREFKISELPTYPILLGYHADNPRSPSEVPFLPNVNKVVRLLGAADSYPKKLDVRFPFSGPWKLYAFYDYENELVKEMVLPGQYVIGEWEMGAPAKPKGSVHEMKVRIHIKPDGLLEVERADAIDVYEVEEAAPKPTEDAAAPAAPAAPPASGAAPAEEDQPQVPAEQANEEAPVKTVKKTKEAALPVSVRPNIDIIGHKSACIAGFRKAEADMHERDCRIISTREKKNELESYILDFRPRISSGGALADYTPADAAADFVRQCDADEQWLYEDGEYASYDDYEKRVQALRAIGDAAYNRLRSREDAEFAAKGFKTRMAAAAKKAMDFVGKKEYITEAELQAAAATADQACAWADSQLATMQAAPKTSESPITAADFDKKAAEVEAGINKVLAKPAPPKPKEEKKEEDKKAEEDVAPQSAPADGAAASANEESHAPRADVDLD*

>Lp_360015700.1

MPGKTILLFDVDGTLTPPRKLETVDMKEALAKARAAGFTLGVVGGSDFSKQKEQLGATVLEDFDYVFSENGLLSFHEGKEFHRNSLLKFLGNEKVVAFVKRCLVLIAAMDIPVQRGTFVEFRNGMFNVSPIGRNCSQEERDEFEKYDNQHHIREKMIADLKSSFPDYPLAYSIGGQISFDVFPKGWDKTYCLQFVEKEYEHIHFFGDKTFPGGNDYEIFSDPRTIGHSVKTYKDTIEILNKMVAEKQ*

>Lp_360012000.1

MSATVTCYKCGETGHMSRDCPKAAAARTCYNCGQTGHLSRDCPNERKPKACYNCGSTEHLSRDCPNEPKSGADTRTCYNCGQTGHLSRDCPNERKPKACYNCGSTEHLSRDCPERH*

>Lp_000442900.1

MRRFLVVACALLTLAVVGEQEVKHISLKRAMNAAVAHSLGMPLVPTRRFKGPQGEVVIHDYLGVQFYGAISLGTPAQEFQVIYDTGSSNLWIPAHNCSLSCLLKKRYQPSSSSTNKPDGREFKIMYGSGPVSGHMIADTVAIGNFSGPQGFAGITDASGLGLAYALSKWDGICGMAWPSISVNGVEPPMFSIAKANPGFANKFAFYLPQKDSDEGDLVLGGYDSRHVDGELVRVDLTTKTYWTVDMTGASIGGQNISAAVTVIVDSGTSMMTVPKALHSTIMTMLNAKAVINGQYSVECSAVPTLPAIVLTIGGKEWVLRGKDYIIGDDSTCIVGIMGLDLPGPIGPAWILGDVFMKRVYTVFDADDASLSFAYAK*

>Lp_000382800.1

MKRFATLLVLTAAFLAALVSADEDPGANMPGVVQLTKSNFNELVGKKQAALVEFYAPWCGHCKSMAPEYATLGAAFQRSKNAKDLIIIGKVDTTEERDLGDKFGVTGFPTILYFPAGSMKPVTYDSERTADAFAKFLTGKVPGLQLNVLKEMNYATELTATNFDQVAKDPTKSVLVMFYAPWCGHCKALKPKYNQVAKIYENDADVVIARIDADNAKNKAIASQYNVHGFPTIFFFPKGEKTEPEEYKSGRDVEDFLKFVNERAGTHRLANGDLSWDYGVVEALSKAAASVARAEGEEAKATAVEAVKAAADKLPESTSTTYYVKVAERIAKKGSDYVSTELARLQRTLDGSMTGPRRDNMLIRVNILTAISKEL*

>Lp_000195900.1

MQYVIQSSSILTAALQHAMKDVLGLADAFMETSAAGPHSVTMNGRTYRGLEAVLLALRMQATTAEQRAFFGDSEDAEQAALVNQWVAVAALLEVDAVRASNDATASVAKAVYTDVEKILATTPNGSAQHLTGSPRATIADLLVYAAAFNHPLHAEALPTTLAWAAHAQQDAYLAPLRSSVVTAHHKAQKAAGASAAAAGRAAEVTYVKPSEEEILRRRAEKEKAKMAKAAAAAGKNGGTSAPSAADDKKASASPSTKGKKAELDSTNLDIRVGQLTNLRRHPDADRLYVEDMVLGSETRVVVSGLVQHYQQADLEGTYCLVVCNMKPRPLMGVTSHGMVLCAEKGDTVRLIRPPAGAKCGDRVLFGANYDATVAAAPPPEPVPGNKMGEVLSHLHTNNKGVLCWKEEPALLASGEKVTIAEMPDCLVK*

>Lp_170007400.1

MSGDLASLAGKYTLTRVDDKGSCEGRVTLEIAVDPNDANKLRLHTKVVNALGCTATLEEGKLKGYIISTQMMGSAGQMKVEHILSRGLAEGMEYKLEDKVLKMTCKTGSLEWTVV*

>Lp_080007000.1

MPDVSRAATRHHHLEAFRALSSLVKSSSDTTRHCSASDGEADDDDDDDKDSRLSVSVVDVERRNDNDVQLRRATPRRTSKGSGRSSSDAGRRRRLTPPCQDYGTGATTTSSIATTASTVACALQQEQPEARLSLADHDDSHSSCQFGRGLSSSSVNEPHQSVQRHPRICHQHQRSFTSPQPSFGVSRVTSPEEPRGVKQQKGEEEDEEEVPLCCSPLDDVYGHTLRITSVSSQATSTVTATAVKTSPSSSSSFPSSSVSSARASCVLASIEDTAQRARQVASIERSLAYEADDEAEQDGYVSMSVTQAMTSKMGADADTQSGSTHEPQLNPSGAAVTETRVASPAAEVTAVHESIEIRDAADAEKKKDDFSPRLSIAERQGGGLLMLSSYRCRRTRSIHDFSQGDNDDEEDEREKEDRFDEGDSGAPHMRKRWYGDSEERCRDESGTSSPLSPPVPRCDTSCFALSSLPSHSPLVRISTSAIMEEEQKAQEKTETITSSLALYNGPG*

>Lp_000486200.1

MTSLSYLYLNNNTLSGTVPEEWAGMTGLSSVYLKDNHFCGCLPEKWRSSYSPSVSADEALKSDRCAVFNRCTSGSESGDSLEGMTDEEVSTLKFLRSFVALNPSLASIWTANYYCDWAYVSCSSYSPSLDFSPYSSPYSGTLVLPELGDDVNGSAVIFTDIKVRSMGVRVTGTLPASWGRLTRLETLYLDGNGLTGTLPTQWGKLAALTALRLDEYRFSGGLPMSWAALARLPSLCVNYSALSGTLPAEWATSTKFWSLYLQNNALSGTLPVEWTSMTRLATLNLSNNSMTGTLPSCWGTMSYMSTLNLSGNAFSGTLPAQWGSMGSLRSLNLSGSRLTGSVPASWGGLGQLQSVDLTGTGLCGCVPAEWAGKAVVADAALTGTDCAVANACSKVVSGSQSESSSSSRRVASSWASGISSSSNTSQEVVCEVEHCVRCHAGNGTSCAECARSYEMTATLQCGGGDAAMRGLMTEGHMWAALLVVATVLVFCAPL*

>Lp_290009000.1

MSGLKKFFPYSTNLLKGTTANVALPTLAGKTVFFYFSASWCPPCRQFTPQLIEFYKAHAEKKNFEIMLVSWDEAVEDFKEYYAKMPWLALPFDDRKGMEFLTSGFDVKSIPTLVGVDADSGKIVTTQGRTMVVKDPEAKEFPWPNVEQKK*

>Lp_250011600.1

MLHHGPHHRKGDKPCTAAIKPEYYQYLLQYIGETPLQAELHRKGEELERPMMMASADEYQFLGWLCETLGVKRAVEVGVFRGVTTLAMALHLPEDGLIRALDISREFASIGFEAWKAAGVDKKIDFIEGPAAASMQRLLDQGEAGTYDFIFIDANKDQYPEYYELSLQLVRKGGVIALDNTLLHGDVVDADESKELARTTRKTNEIIKADPRVSAVMSTIADGVYFVRKL*

>Lp_000274700.1

MSGALTIHDNNKTVQAKVGHPIHITLQGNPTTGYKWTRLNYEEKDMLSDDDMEVVAEYKQSPSAPGMVGTGGSYNVTVTPKRPGKHTVELVYARSFEGPKPDNQKYTLHLDTH*

>Lp_000079700.1

MVAGTLPSSWGGLKKLISLLLYGNALNGTLPSAWGGMTKLETLRLEKNTLTGSLPSEWGSMAKLDYLYLNDNALSGTLPVSWAWMTSLNFLSLKNNNLSGAVPEEWTGMTELFSVYLEGNHFCGCLPEKWRFSSLWVSSEGALKSDRC

>Lp_360034700.1

MTDYKVKDISLAEWGRKAIELAENEMPGLMELRREYAAKKPLAGAKIAGCLHMTVQTAVLIETLKALGAELRWSSCNIFSTQDNAAAAIAKAGIPVFAWKGETDEEYEWCIRQTLKGFSGDGLPNMILDDGGDLTNLVIDHFPELVPKIHGISEETTTGVKNLYKRLAKGKLPISAINVNDSVTKSKFDNLYGCRESLVDGIKRATDVMIAGKTCCVCGYGDVGKGCAAALRAFGARVVVTEVDPINALQASMEGYQVVLVEDIVSDAHIFVTTTGNDDIITSEHFPHMRDDVIVCNIGHFDTEIQVDWLQANAKEHVEIKPQVDRYTMVNGRHIILLAKGRLVNLGCASGHPSFVMSNSFTNQVLAQIELWTNRESGKYPHGDKAGVFFLPKSLDEKVAALHLAHVGAKLTKLTPKQAEYINCPVNGPFKPDHYRY*

>Lp_000051100.1

MRVLVVAADYSEDIELVCITDVLARAAISVTLASATASKHIVLSRGIRVECDALITEVAAGDFDAVLLPGGMPGAETLGKNETLKALMHEMRSQNKLYGAICAAPAMALGPMGLLEGVETVAGFPGFEEKIPAGVKYSESAVVRSGNCLTSRGPGTAIFFALAAVSILKSPELAEKLAGMLLVDKMSEMDAVRALK*

>Lp_000051400.1

MRALSFRTGVLARSTAAMLCSQRWCAAGTNAEKPAEDKEAAKNAEEVVPIETLKKIEKQLEEATAKIEELKKEILYRAADAENARRIGREDVEKAKLYGITSFGKDMLEVADTLEKAIESLAAFSQEELRDNKIIESIHTGVKLSSKVLLKNLSKHGIEKMGVSVGTKFDPNMHDALVSTPPTEQAPSETISSVLKDGYTLKDRVLRAAQVGVSH

>Lp_000234500.1

MSIVVAIATLSVLIGLATVYIYTRKMSAAEEQLKTAEAEGAAQPPEVPISGEAEALAAADDAAAAPATTASQSSASAKKNKVNSYYYWHGHEKERAKLGDVAPMPTPHLVSRDDNVNIVVPAISVAKYSWCDGDKFVSVYVETAPPASGEALEESTIDASFTCNAFRLAFTTVDAAGKRRAKQMVTRLSKHIDPERCSTKVRPKTQEILVKLAKKVPSMWIDLEGNASDCGSADDEPLEKYKSDENDE*

>Lp_360019400.1

MSSASHFPEVQSSKHAQRTLQPLTELTDKMKPSHSTKSLIKLSMGDPTADGNLVAPQILVDEMVDIVKSKDFNGYPPVAGYNEARQVVADYWKKFCGTQERKDQIKWKNALLTSGGSHAIVLAISALCNEGDNLLVCAPAFPHYKTVCDSYGVECRYFLLDSAKNWEADLDAAAKLVDSKTRGVLFVNPSNPCGSNYSRKHVGEIIEFCEKFSLPLISDEIYAELVFKGEVFTSIADFDTDVPRLVLGGSAKHAVTPGWRIGWLILVDRKGVAKNWMNGIDRLSQLLTGSNSIGQMSLVRALTKIPQDHVDSVVAQLEAGAKVYNRLLEHDIGITFDAPRASMFVMLKVDLSYFKDIETDMDFYEKLLDEENVQVLPGEIFGVHGFVRATTSRPAAIINEAVDRIIEFCQRHKK*

>Lp_000088100.1

MAVRDRLVLLAVCLVSALLIAAAVAAPDGSGKVEPPCIGVDLGTTYSVAAVWQKGEVHIITNEMGNRITPSVVAFTETERLVGDGAKNQLPQNPENTIYAIKRLIGRKFADPTVQNDKKLLSYKIISDKAGKPLVQVTVNGAKKEFTPEEVSAMVLQKMKDISETFLGEKVKNAVVTVPAYFNDAQRQATKDAGKIAGLNVVRIINEPTAAAIAYGLNKAGEKNILVFDLGGGTFDVSLLTIDEGFFEVVATNGDTHLGGEDFDNSMMKFFVDGLKRKQNIDISNDQKALARLRKACEAAKRQLSSHPEARVEVDSLVEGHDFSEKITRAKFEELNMGMFKNTLIPVQKVLEDAKLKKSDIDEIVLVGGSTRIPKVQQLIKDFFGGKEPNKGINPDEAVAYGAAVQAAVLMGESEVGGKVVLVDVIPLSLGIETVGGVMTKLIERNTQIPTKKSQVFSTYQDNQPGVLIQVFEGERQMTKDNRLLGKFELSGIPPAPRGVPQIEVAFDVDENSILQVSASDKSSGKREEITITNDKGRLSDAEIQAMVEEAAQFAEEDRKVRERVEAKNSLESIAYSLRNQINDKEKLGDKLDADDKKAIEAAVQVALDFVDENPNADREEFEEAREQLQKVTNPIIQKVYQAAGGAAGEEPDAMDDL*

>Lp_320024400.1

MYIQDPSGANRDTMELDTMPLSQLYAVGYRMYRKKHQPDPSQNFALPTAPPDFDGDSAIDLQTVVRHVRSLANAYNSVIPVPTPFAIYEEVILDAQETLKTTTEEANQSTLFINSEVANTPLPECTDNTMNDLFVYSSTFGAQLQSDALMMSVNLLNGGQAHLRSLQLLYHRLLSDVNLSRALAPSVMSELLVQLDAVAKEAAQPSAEVLERCAIAGLLVGRRHHLNEVLRYTEILEQFTLSSASIVPAEVPHAREIVSSFSLHAADMCPITSPLDCFQDDPTNVKVMLGEGTSVQRFCVSPNQKFLAAVGSQRVIVVSLSTGTACAERDLPDVAGRCWNVTFSEDSTHLICTDASGLRHAVLPTDLSAQDDHHTETVSIVTDYRGGALETRPAPLLPGGSVSSLWLLEPVMADSVSFPVASPQDCVECISLCAHAYHDCSGTLLRIIPDSGETLCVELSANLIRMRQGSTMCQGVLPSSSQWYFIHVQWSRGVWSVNVNSSIIELSGPHYTQSQTMKLKTLECLHCVGHVGSMAVWSGVNGPSTVRTFFRERTVDSSVTPLFVLPMDEGTGPWLKETVASQCVSLNCKVFEWDPAVVVPVTFAHARAPAAAAVPLDTQLLLSGRILKSAYRFFIVLPPTEKGSDDMVAEVERETNRVTRVYQCPRSCSENAYALLADESELLVYQQRFSTFSSVRLFTRRRMENDLVLCNAGAASVAEGSALWLLEQIYRSIYTESYELERIPSENTPLSFATLRSLAQSLRMVKTPAAPLKVLAFMSLALVHFRRLASLNDSRISDIAASLKASCSVIMELFTEKTFFEVAYKLYQMATAWSLTVEDQTEMLLHMESGEKSDLQWSYFSRERVYTVIHHIVQNRPADIVSAAKNLLRQCERESADESSNSKPVIHLMTSFVLSVMTTLKSKPRDTFEVYVGKLIDVFVEFMCKNPSLLVPSRVDEQPNSVLLTGVLPLFVAVANLCPAAISVEAQEKLLELCARLRAVETCSAQTVSFTTHEAYDIYVPPKENGGIWRLYLDFSTAKNVSLIKEASSPPMFVLQTDSKTGKPDNTVMDETSSFRKLEGGVLHVHGKGENNFTAIAQYELVLDVSLASVMRECILRLLLAASAELMAKPAMTDIAMTPLFRCGLTTSVLAGNGVVLSGPRRENTDMLPRRVLEDDGEGKEFVDEVCKAMRGSVVPTMRPAIQALLAILLQCGLTRCQAEDELKLRWFSGEWGAKLVGAAKEPTMAFLTELAQWMLDHVYHESDATTPFASSARRSSVERRLDNSLNGSRSLSERRNSTSSRRSVVKMYSQTECLEQVAHLLSKGITTTIIEKVLIGRARAAQSIEKGLKVLSKVLQVNQESLSEEILTDALRVLANFTYTENFQHFTEVLRGAGADAEAAVRGAFHETLHSFFHVLKTSTNALSYRAALTQAAHRGRALVLFQAILCLPWDEADCEQFRKNFETDKDILKFLENQMDSPTGLGVLTSPWRVRHEVSRSCEEVPADQEQAIRTELLCAADFASLSRVFPNGFSVAIPGVQTARSQLGVEIEKSAGAAVLIRADEGWQLSWKEGVPRVYYYEVKIISPINKFDIGVQLPREEQTVRAENINFSYSSTGEVQGADLPQWPPFVEGDTVGCGVVATSQQLFYTLNGRFLSYGGRVAVSNTKSDNALLYPFISFDDGSMVRVAINFGTMPFAYDYRQLHPALTIAMGPTWYRVTSTAEMVLHYLTARVCAVQESKNQSARDALSQACTVVCRNVNSVTSSLMNIGVGGSANSNPTTRVKQAVALSVAEYSIMMLISTIRHIATHGLLPSEHVSNMVFDAVSILMLLPLHSVQIATIGLLPDVIVHVKGVPDSKAAGNLVSTLFENANSVTDSEEFIPFVPTWCECDSIAMAIVNVQHAHMLPEAQRSIVLGNVLPRTGMVSFSVKVTRRNMAKGHSLKGGYFIGVSVAGLSPLTPTSNSQSWKAVNPPIVWAIQDTSPQLPHATNPTVKLNNFQRTFGNADVIRVVVDRDKRCIDFYREGEFLKTLFSDIPEDIDLVPFVQLYNDDAAAAISTGEMTAPITGPTLLGAASVDVLRSMLTLDPFQDLVANRLCEELKSKKYPKVTLPIFNSVPDPRLLQLRSSSGEEVTVTVSRLNDLRCKFSLGGKTNYEHLYNMRTPAKPRTTCNFDTVSSTSRLTGLGRCIDELIAATERMVIPRITVEALCCIEQDERSKIVLEEKDHWDFLFHGLRVGSFITVLRRLPSVPTMHPPPIPQDFTFSAALSNPGFVLPSHYCGRLATVPNGAKGVSTPFIAIAEPPISNNAQVSIRCQLIRGNQGQILGGGYYFGVCTSSFHWKRKDLNSHATEVWALHDMDDSPWRLRHLRCDATFPLAADPRCIIVSGDIIRLQIDRPSRTIQAFRTPVNGEELPLGIIFDNLPPDGMLYPFVNLFNTDAVAVLLPSNSSAPALRLPCQKLQYALCTADERKTCDGCMTSHADSRITNCWYKCNECADYTLCSNCFLTCVHSNHSFTMMDGSPIAYSTTTPTTIERGMEVTIPATTAMYLRSSGCRMSEAKMNCVAEAVEDNALCTWGLVGIDDAETFSVVIDTIDGSPLSTDAPVFVGLGPAQEIVQATREKLRNKCLSNISTIATFCSDPTLRQPMEHKSRDPYGFQRGARITLRYDATTGKITILRDWVVVGTKSVALKDSENSVQPIGCFVLLGKKGTTASIFPERATTYTTTVEDVTGNILKTTGRDGSVWFVTRDNCRLPLTPCRGIYPHSSLGYVLIDQHLAQCCRLSFEGREFRVKVLETEEVITVPANHFLTNERSAISEEWFLSRRERNEGATVEEGFVISRLLLTLSCLCENESLSHLVSAHQDRLLSMTKRLATVKIQNDAPVEFLEEIRASVASTANRRGWWLEERHIAPFVPVLNTDERKRRFHSGTLVHTLHGGNDVFEVVSRRGESLTVRNTRTETLSRLSYRSCVAMKQCGGRAWYTRKNGLPCSLYEHTVRLNDQARAAENVPLQAEWLGELTFARAAKSLSVLLAPTGIGRADIASGTTEETRMLVIYEHKRYGRVVRLYMRPYDADLQVKLQSFVAEDEAKEATPRTFQEKVDFIVGAAAASSADAATPAAPAAEREVFFEMTGYLDAEGLRLNGTFQCNQGKGGSFTLNSARRFALLPAVSDKWSVEPLPDCVDVAIPPEPYHGMRECTHRLVVLLARHLYLVICSQIERKPTDFLEHIFNFTNHPLADLLVSRDADHHMLSVMLSETNMRIVDPAIKPWDAISLARLATNSLLLSPEAINNVPEHLLWDCLHGVVTAAHRCGRYRRHEIIRSITTFVESRLDYTDIYVQLFSTLFSYVERWVVTLCEDAEVKKDVAAGVDLLLALDKPLSEKLRNIPVAPLHCLIDLWRSLSTGVALPRVVDKLHMAVRPDAPRVDTLCTFGDYVRHELKVGKISSSCGTEGKGHYYYEVTLPDRITSAYAIGWGTAAHNEVPGQHVGSDRNSFAYNGSDINSREGKEEYKIPTESVPGSVVGCLLNMEDRTAAWSLNGVVGHFIAIPISVGSAPLFAFASIGTCSGMKVRLRANEFVYAPDGYTDLSGYFARPLVDQYDRALRSEMPLQTYAFYAQLASFLSDTDDYFDEQAERNRTASVSLPPEDPYTLFLHSYPLLQPFPPATLQRFSQLIAVTESCMATANRFVDLDEETVTGTLSQAFLSMKGLIRRSFRQRLLVNVASLEPSTSAPTIFVRATDLYSNLPRTTESALQHSILAQLYRQIGFFTEEQWAVTPLFKVNLHISGSGHTPVDMGGPYRQVWTFLGFEMMQHPDKCYPNSDFHRNPLFVFVNNSQRVSLVPDSQANSAYDLNLFTFFGKIMGHSARAKTPLDIDFSPFFWKFLVDDELTVKDYYTHVDSVVEATVQDDNFLLSGVADELIPGFAESVEWLDADDNDEVSAQQRRTIAERCLVHSMDEQLSAIRCGLWSTLSRRVVRCLSWKDLENLVCGESNLSIPDLQKYIRVQLTGSREAYFWQIVEQLSMEQRASLICFASGQRRLPLIRPITVAENNESVDHLPRAQACGSVIMVPQYTSLEMFKEKLVIALQHQMEMELA*

>Lp_000121000.1

ATMPAAPKLFVALAVSTVAVFTLCCTLAVGAAHHGYPSCDLSVVQSSGYIDIPGVGGTQKHYFYWLFGPRRWPRDGSRPPVIMWMTGGPGCSSGIALAIELGPCKVNETSGELYRNAYGWNDEAYLLFVDQPTGVGFSYGDKANYVHNESEVAEDMYNFLQGFAKRFLSPSITGGNDFYIIGESYGGHYVPAVSYRILQGNQRGDKPKINLKGIAIGNGWTDPYTQQPSFAEMAYNGCKEKLGAPCITEAAYEEMLSLLPECLNKTRECNNAGLDYGASNAECVQAKAVYADYEGYYFATGLNNYDIRKPCNGPLCYPMNHTLDFYENPLVRASLGVSDAAKWSTCNAEVGELFEYDFLRNFNYTFPPMLAAGIRVLIYAGDCDFVCNWIGNKAWVTALEWPGKAAFNAAPDVEFSVNGRAAGQERTYGNFSFVRIYDAGHMVPMDQPEVSRDTVSRFLHNKRLA*

>Lp_340015700.1

MDTSELQKHFEAFASFGTAPSKDMDNSHFSKMLKECKIIGKLFTSTDADLLFSKVKAKDARRMTFIEFKDKAIPEIAAKLKKTNEEVEEMIASAAPKSSGTKAEAVRFHDDKSSYTGVAKQGGPTNVDRNAGSLAGVVDRRQATIDNRGTTAHQI*

>Lp_000342300.1

MPNKEYPTEDPRPRENDPGNSEPHEEESVCSSEARDRKDLEECPGVDLRAVPSEHSGMVVMIFYEIFAFISFVFMLTASCPIPWMNDGSGRKWTVWKDVDKKRWKNHPCDHKRALFQGMEAFAICGCIFSLVCFIVGLLQMLGIGHLGVTLLFSFINLIVLITDWSLLVNQYHKYNCPGELSYVSHINRLNAGFALVFCSFGLMFFGTVALMYWTYATFSLSETQRDKYSTGAFNSTLISAALLIITTVATAQTLWEQYYEKYTVKVTYWHVEFYDRETGLSEYWSLDAYKCSQLKSQIHAAAAFNIISDVFLFVTMLCSIGAVYNRFCKWLTIGFGCASWVFLLICWAIGVGARYKTFCKNGVVPPLGGVPLNTEKQRVSFTGYVITEGLGMIISAWCIETANLIYFGLRG*

>Lp_360049900.1

MKSLYPMARRSAASYQLHNRTIPEPYDYLEDPSNAETKAFVHQQNEAFANYMKSSQGLRDKIVERVTAMLNYARTSNPSQHAGKYYFHYNTGLQNQSVMMQSNSLDDKNPSVFLDPNTLSADGTTALKSHGWSRSEELFAYSLSDKGSDWQYIHVRNAKTGEQLQDKLSWAKFSDISWWKDEGFFYERFPELAEGADQGAETDSAQNQCVCFHKVGTPQSEDVVVLRVPEHPGWNYMAEVTDDDQYLVVSPMDGCEPHNLIWIAPLPATAAELKEKPLEFRKIVDSFVGEYRYLGNEGTTFYFTCTKDAPRKKVISMDLETAAEADVVPEKKSVLNIAALVKDTLLLVYIEDVKDVMYYCKLHGGASHMTKLDLPVGTVTSFFANYHKDFVSFKISSFMLPGRSFVMDINDPPGTLAIFRDDTVDGFNADDYMTEQRFYASADGTKIPMFIVHKKGALSAESPVLLYGYGGFNVSLMPAFSPSRIVFLQNLGGVLAIPNIRGGGEYGQAWHDAGRRAYKQNCFTDFIAAAKFLHANNIGSPATTAIMGGSNGGLLVAACANQAPAEFACVVCQVGVLDVYKFHKFTIGHAWISDYGDPDKEEDFKVVEKYSPIHNVRAGVQYPAILVVTGDHDDRVVPLHSLKYIATLQHANPELGGPFLARVEVAAGHGFGKPTSKIVEETSDMYAFMAKNIGATWHS*

>Lp_340007900.1

GDSLFVSSASDCSNTIATAGDGCPAAGSHAFSVSSASLQVSLTKPVFSVPPTYWCISNSGGIATSIGMLQMNVVQTIPMYYPKTKLAAMTFNEATPLGSIIRFYYADGTCQAPVVDYAQFILDENRTVEVSISVNEVGVCAIISSPVKGDTTSEMTLRNTLFSVPVLSITPTTGVRFSNVSVKTNSSSIYYCALSDSKICDDLPYPDYLQNDHNGTLILKVTKPAGNYYFCVAEGKKLYFPATSMFSVVEYGVQPNTAYTQWSTKIYPSINATSDMQVALFSSKDCSGIPVKDWATFTDASWSVSDAGTYYACVRASSSTMSDGYYFANAYTVVPLPTLTARQSVVVREFNLSLSFSVSTAEMPLAAIFVALSFNAACASFYRRAKALNGETLSFGIGPDAADTLYWCVSNPIVAGTADNEGTLVVSYAYVGSTSVRDFQVSYDPLRTNSPSTIVLDKMAALSSGMQVAFVPSARYTCADVSGKANGAVVVATVSASNTLESVTFPAAGQWRLGAQLNGSGTALTNLQSVSVYGDASVSPRGLVPKVTTALKLSALQPLQPVFITDAAACSNDSSALAAGEADSSGAASFILTYNGTGTLLVCGKYTGVEDGTTASTVKTMKAATVVSANPHIYPTVAELSAGKQELRLVGGGAHMLVGRRLVLTTKSTACPSSLTLPPFALVLSPLRLGSNAETPATTLTLHDNMMDTQFHACIKTNDTYVDVGMITVTRASLKTPVTSSPSPIALVAGQPSLLTLPSNYTSVALVDSYVVVDADADCTNDLGSTTVYTSGSINPNTGEASAFIAPSPASDSTTATVDLRLCVAQQSQLLNSTYGYLDGGALVSSTFTAVSKYALSDLNGGVTGWPVLSYASLYLVSCSGSSCDASAASATCRTASPQYTVSEATNTTLKPSMGTYLLCQRVTDGATTTVVGSNSTVQVLNAFTMSTTADRNQLRAFVGFDAVVTGGNALVVDVVVQPLAVPCGTSAATSQSFRFTTNQQRSITITALTAYTPIHFCAKPSGQHADAFEVMTATLNSFVSPTYIAFPSLSSTSTISIPQPSMTSLQAMLAREPDCGSSVVGGRLTNAVGGTFTFTVSPCGENSNLDVVHFCDLSSGAKVYRGPLLMLRNGNCSANDGSASVRAVTVAPGAPITDFGVNTHFLAVLGISKTASCRTLIAAASVAAQGYVPIADENAVFYVCTYVITNPSVTFTTQQATLTVRNYVAVPSTARSPLSIISSGPARVLVTLNTALPSGYTFFSIGTGCLASISDAPGLNTSAASAVYSQASLCGTVSVCWQSASDSAPLAVAQFASVTGPVLQSSTLAIVRGASYTATFARDSCSSATTFSNAKLAFLSADQCATRLDGTGAGVDALVFTTSVTSDLVTGLSTANLCANSTAGIVTVLAGVPVARDRVYPVVFTSGVAEAKIFIPSFARAFFWLTSADTSCRATSAQTTSLPSFTTDANGYGLLTVVSANGMPLPVGTYALCYYSSGFAMALESIEVVAPTFFDVRGTSFVTGVASKMLMQDDLQASDLIEGFSTTSNCSTLSTSYGTWARVSATSINVTATRAATGGAYLCARLPLNLSVAALPNAWSNTAQSLTFAPSTLHLPSTGFDVCTRYTLTQCTPPSGGGDDATNRLTVVYGDCCNSSDEANAVGVGSMAGGACSLSFDEAKVRGYPAGTVFSLCAWNSFDAAVCATLNYVTVNTNCARDVTEKRSGLSLGAVVGVVVACVCGGLALMAGLLYLLCYFRRRAMEEDKASESATSPTARRSRSTTARKFARPSGQT*TCRLRTTTAAVRGDLTSAAASRRCAAPSSRRWPGAEAGGTAAAMMKTTPSLDSWREPLDTNFLDGCEELAEKYKDFINGVRRDDSESVADFPIAAFMKRSKLDTEALVALRKVCTQSAAVSDLLRPSVMTLQRHELLASMERTNPAAKTLYLFFEEQNIARIAIEISEETDFFNLRALFRSYNVMLHAQQSPPSKSQPAPADGGGNGESVFLMENVPNARGRVVEPFASPHWSVALPSADATPAQTRERQWLYRHRSELEFPWVTNNFRTLVGYRKGNGERYNYMDWSLTLLDLQAFHPLHRWWSPLLPCVSPSPVDACDARFAFMHANSTVPFVKRYLTLFKVEYLERTQI

>Lp_050011700.1

MSQKALYVTPRAVPYCTLAIAGAFKDLTPKQRRYAHHMMMAGWCGTPVVAEQLSPESRPLLRFFYDFLSAQPLEMLKANVTKAGVAAEAVDQFLEYFAMVYSNMGNYISFGDSKFIPSIPQETFAAIVAKAGAGPGAVDAALLNAIYDLHEDKLTLDFPPKGITRYYSPNITREDAVIANDFLASKKIDGVNTRVFKEEDGTLVIRVAAAEERTVPAVAFNGRAIAMYFGDFKEEMARVVAELRQARAYAENETEVRMLDHYIEHFLRGDVNEHKASQKEWVKDVGPTVETNIGFIESYRDPSGVRAEWEGFVAVVNKEQSKVYGALVAQAEKFIAQLPWGRAFEKNVFTSPDFTSLDVLGFASSGIPAGINIPNYDDIRQNIGFKNVYLANVVSAVTFKEKINHITDADWEVYKRSLLAATSVNVGIHELLGHGTGKLLTEGDDGQLNFDKNTIDPISGHPVTSWYKPGDTYSSVFGGMASSYEECRAEAVSLYLCLQPELLKIFHLATEQEQQDAIYICWLNMVRAGLCGLEYYTPETQQWRQAHMRARFCILEALARAPNPIVHIAEDEKEGLLITLDRARIATDGRQAIGELLVNLNVNKATANAVRGMEYYRNMTTVNDTYVRYREIVMARRKPRKQYVQPHTRIVGDTVEVEEFAGSVEGVVKSFLTRHQEVPL*

>Lp_000025800.1

MPFAVQPLPWAYDALASKGISKEQVTFHYDKHHKGYATKLNAAAEANPDLAKKSLLEIIKTVKGPAFNSAAQIFNHDFYWRCMSAHGGGEPKGKIADAINESFGSFAKFKEEFTAAANGHFGSGWAWLVKDTTSGKLKVYQSHDANCPLTEENLKPILTCDVWEHAYYIDYRNDRAAYVNAWWNVVNWEFANKCYESSSGSSCTNSNL*

>Lp_000025900.1

MPFAVQPLPWAYDALASKGISKEQVTFHYDKHHKGYATKLNAAAEANPDLAKKSLLEIIKTVKGPAFNSAAQIFNHDFYWRCMSAHGGGEPKGKIADAINESFGSFAKFKEEFTAAANGHFGSGWAWLVKDTTSGKLKVYQSHDANCPLTEEHLKPILTCDVWEHAYYIDYKNDRAAYVNAWWNVVNWEFASEQL*

>Lp_140009400.1

MATSVDLTRAPSATGKSVARTMVAIFKEEADAPRVLGESDAFIAYTLKNRRAIRLICRTNNDRGAFRRHEGVVHELRFVNFHSNVAASASDVDFFVWVAVSETTADGTASEKADLSTRLTVKPYFKLVDAVTIQRFHFFINPANNMPDLLILYNKTAAVLQSSKLISEYKSDMLEARLGQQSRELRGLPQEASMQSLCSTFTGGWLAFTAEATKVMACTLRNTTTPAWSGCEGEEIVQLSFVGANATTTTDSAAAAAAATAAAATAGSNDSSALLAICSHAKVCLWKLTGVAEPVLLRTFVFGRIITMLSSRNAFVVFNTEGNVARVQVRDEANMDAVVYALNANIRPTSVCYHEAAHFATILVDSVMELLLFQLATDPSSNNNNSSSAAAAAAAAAGAGSSKSSPAAASGATAATAATSTTASMVNTPFVDPSIISTGIPFRPNLNSNNNVARSTPDAAAAAAPSAPLSSSSPSAGPTGVAAAPAATSAAAINANPMAAMAAFQSARAALQNASLPPSLDGLVASAVYQSDESVRRSIEGLQNVLKNITQILQLTPDQLLRDHQQLVSLALEAQMTELQQQQQSSTAAATAPSAAAPASAVSARHSEAFNSYALAKLLAPVASEIAIGVTRGVRDTLRVELDAAVNAAFGANVHASQKEVLRRRLDEALQGSARVLGEDVASRVDRYIDQELSESLARVNASVSALEEQNRRLQAALHQVANSGVVQEVEAMRAELAQLRAAIQRGDAISGDGAAAAGGNGASSAAAPSPQTVLETAEKYVKEGRITQGLAYIVMAQNARCVVQLLSRLSEDDVANVVSDAGTTETTWSKLLTQLCEPEAADGVTELETVASFLFDVLSDHEDLILKKTAKAQQIRSDVRHFIARARSKIDSSEGRRVLKDLEKSIQ*

>Lp_000397700.1

MHAEPFSVENCLRYLAERGYSMQSPDLRQVAQLSAGKRATHSLCEGLNGVYSNGIHIGSIFLILGVSFLGTAIPIAGKFIPVIGRYPFVFVIAKSAATGVLLSVSTIHLIHEGMMAFREPCMNSTLKKYDPVYFLFALIAVLLMQLLDIQLGEVAERWMKAQLAAEAESGAMADASNDDKQQQQEDQEQERGDDGFAEAEVGRPAHGLDLDAIECDAMVGSASGVPDEPTTAEQSKRGDSATELATVELQAHHHTHSHPQLGHEKDGECDAHGHQHLNVQPPRDMGSIRRVIAAICMEFGVALHSVFVGLAIGLTTDSELKPLLFALVFHQLFEGMSLGSRLADADFAGSLEIVLAFVFSVSAPVGMAAATIAVSVSRNAMSGSGFVTMMAILDSFCGGILLYLAFTLLLGDFAADLRHHCADGLPNRLWKKIGMFAALWIGMGVMALVGNWL*

>Lp_000018000.1

MVATASLTLLLLFCEAAHVAHGWAFVPVVHNFAVKETTASVVQSRLVCQEGGGYIAAEPTALLHQSVVQTVRAAGAADTWYTFLGGDSHANAALNCPKPIAIDELSGASYGCYWRWNQGRWTEMDDARALPLYIGGVGVTFFIGNTYKVPLSSSVTFYGQTGGFPIFFDTAAVADRDKRPGMLFGKNLVAVGKSDTSTWSDNLESGGYSYAGYTYNSVVGVSRWNADAQTASKADFWTVCQAQTPSRTMYELENTSSGLQENWWVIFFVILFVVMLVVFFIIACCQDDEDMDEPPEDAPVWAQEETQQTKRTKSFVSTRSFRRNDYGDEDDDDDE

>Lp_000307600.1

MPTLKEKNPASPTALVESAQTPGPAEVSAEELHVLLNSADKTAVMVVDVRPAEDFGAFRIPGSINKPFQELNYAKLISEVAPHAKQEGFLLVFVSAQSPDVDDLAAREYMNEFAKTHSHPPGDGAVSILLGGVCGYVQRYPS*

>Lp_000372800.1

MQFRSIPRLAAAAMVALVCLLALSANAASGPISDSDTHYFVLLWAKKYPLNYVWSGDNLCKWEGISCDTVKQQVTMLLPKVGLTGTIPMWGSKSGFTPANVKVISINLAKNKLTGAFPAHYGQLTKLQELYLLDTSLSGTIPQAWNNLASLTIVDVSNTKACGGVPAWDQTSMPSLQYMHFTNNSLMHGTVPASVMTFSKVSFNTSGCHLCGCMPAGTSPYIRVQLARDQPQLATSTCATSNTCSASDLTCMTPKTKPSKGSSHSEASAPVLALSSLVAVVASVVAALAV*

>Lp_000417900.1

MAMSGVTLADNVRGAFNDLRLKKSRYVVMKIGDDGKKIEVTEVGARDVSYADFTSKFTAETPCYAAFDFEYADAGANRDKLILIQWIPDTAKPRERMLYSTSRHALNAVSEGYLPIQANDASELNAEEIVRKVKQHRSA*

>Lp_130009600.1

MPSGPKTNKYTNRHSEEARLRDEERRLDARAAREAAKEDAKWEETDAKVLKKMEKQREQDRKQAEKAARDAEKKQQLADEEREMNAKVPAKVTKRQIQKDLSKMLADYDRQRDAIRGVQHGAVVEPAQKEEEETPLPSGNVNRQRGIVPFAGGNQGSDVITASGKTGDVLAALEGKTTGEAEIPDNRHIGKRARVLYKAFYAEHYDAVKEEKPGLRRAQYNDIIWEMWQKSPSNPFVLRNEKVAQQRLDEERRWMEGNSGDDEEGEEDEEKE*

>Lp_000082200.1

MTASKLLVAVVAAVCVVLAAASVPAYGLHVTSSAAAQFEEFKRAYGRVYATLEEERLRLRNFEHNLETMRVHQARNPHATFGVTKFFDLSEGEFRKHYLNGASHFKAAKERAAKLPAVSADVSGAPATKDWRDDGAVTPVKDQGSCGSCWAFSAVGNIEGQWKLAGNPLVRLSEQQLVSCDTVDAGCNGGLMLDAYDWLINNANGNVYTEESYPYVSGTGEEPQCNMSSGLVIGASIEGHLSLESDEDVMAAWLAEHGPLAIAVDASAFMSYQGGIITECEGVQLNHGVLLVGYNTSGSVPYWVIKNSWGTDWGEDGFVRVRKGTNECLITEYAVSAQVSGKTHAPVTTTTTTTTTTVGPQPTVVEHTTCNDYSCSRDCSTTRVPVGECRKGRNGGSLSLECGEDQVVERSYKSSDCSGTPKYYVTPANQCMSVWWGSFKDVCV*

>Lp_000082300.1

MTASKLLVAVVAAVCVVLAAASVPAYGLHVTSSAAAQFEEFKRAYGRVYATLEEERLRLRNFEHNLETMRVHQARNPHATFGVTKFFDLSEGEFRKHYLNGASHFKAAKERAAKLPAVSADVSGAPATKDWRDDGAVTPVKDQGSCGSCWAFSAVGNIEGQWKLAGNPLVRLSEQQLVSCDTVDAGCNGGLMLDAYDWLINNANGNVYTEESYPYVSGTGEEPQCNMSSGLVIGASIEGHLSLESDEDVMAAWLAEHGPLAIGVDASAFMSYQGGIITECEGVQLNHGVLLVGYNTSGSVPYWVIKNSWGTDWGEDGFVRVRKGTNECLITEYAVSAQVSGNSTTTTSSPSTTESPSSLVVEHTACHDHHCHLNCTTMQVPVGKCMEGYQNTSFSLVCGVDQVVKLSYVSSDCTGLAKYEVVDANKCVKFRYGSFKDVCVPA*

>Lp_000309200.1

CTPYITAANGGVACLSKNPINLRFLLQMAHENDYTANGVLEDKATYAFQPCVDSIAALSLKGGPFLAAGYAVTSGTTTSYYFTKNKSDKYLSVYGFGMSNQTTVPEPGISTGAVLIAKNGADACNDASMMGVVTVLSLEIGMHKGFFNYIGQSGQTTKTVANPHYNATLCGESSSGSSHTTTTTTTTTTTKANADASSSEGIDYCAKTITVVVPANNVQPARAGFVPTCDAKDVCLMGSNDTYTCIGNVTGKKNCGICTNNATALAGNAALTVWVSYIGTDRKGNVMTSSGDSPFNYLNYVRNQAFETVTSKFNSLIHGNFSD

>Lp_130007100.1

MSAEDTPVSLFLQACKQEGVKTPNPQLVAFFERHKTFDDIEEIDLSNNYVGNRGLLAVLDVIEHLSQFRFLNAADQKLYNSDLNEEAVKGNAVIDRLVEVLKVHPKVNALDLSSNPISNYAGRKLLSLAQVNKRMCRIVVNDTRIDFELRNKITKQCEENTTNLWNEEQEGSVTADTAFGEGPSWVPTQVSADLTALGGGRARRQTVHVEGIDPEAAKNFVAPVHEKSTEETELICELLRHNVLFSFLSSKDIQTVAGAMYREEFVKDECIIEFGQKNCNKLYVIQSGQADIIKEGQKVFLKTEGTAVGELELLYDTPAVATVKVCTEQLVAWVLDRDTYRNLVMGSCIRRRETYVAMLAKVPFLQSLDNYERTQIADALTSDEFQPGDYIIHYDEEGEWLYIIIEGTVEVIGRDAQGNETKVCEFTSGDHVGELEFLNNHRTVADVVAVTPVTTAKLNRRHFEMCMGPVMDILKRDSSSAKYEYYQNVLQNQHRKPEATEEL*

>Lp_130011200.1

MAEETTRVEVPEVEENAVVDTAPESLEDAVRIVIQKSLEANGLVRGLSEVARALDCKTAHMCILADDCEDEEYKKLVTALAKQGSIDLINVEEREKLAQWAGLVRRDMSGEVTKTLKCSSVAIRDFGERTKALDFLLSQLH*

>Lp_330019100.1

MFRRNFCRLSGLNKVCSIDEAVAGMKDGIQVAFGGFGCAGVPDAVVKAMVKKGVKDLTLFTDSAGIDGFGLARLIEANQVRRMCCSYVGMNKIFTKRYLEGDVELEFTPQGTLAEKMRAGGAGIPAFYTATAYGTQRQTGGQIVRYDKEGNPCMISEPKETRKFGERWYVLEPTVRPEYAVVKALKADKSGNLVFRGTARNFNIPAAQCGKTVIAEVETIVENGQIHPDDVHLPGVYVNHVVQASYEVPIEKRTVSGSSEKKSAVDPNDDRQKIARRAALEFADGMYVNLGIGIPTEAANYIPAGVSVTLQSENGLLGMGPFPTEDKVSADWINAGKQTISFLPGASCFDSATSFAMIRGGHMDLTMLGALEVASNGDLANWGIPGKLIKGPGGAMDLVASGSRVVVVMSHCNKNGEPKILEKCSLPVTGLNCTSRIITEKAVFDVIDKQLVLKEVAEGLTVDDIKKCTGAHFHADSVKPMTYAPVRA*

>Lp_000254100.1

MGRSAFALLGVVACLAVLLATVPGRAAYTPNGATSEWLEDWLKAIPNLRTIWLNPVICSRAGIQCSEATQTISIRLDGVTSSGFNFIGSLPEAPSRASELQVTSISVRGKTRISGTVPASWSTIKSLTGLDFSQTKLSGTIPDSLGSLNRLVVADFSNSFFCFGMPNWNRTGMPSLTQVSFKNNNMRGTFASSWSTFSAALSLDLEGNKLCGCMPSSWQTPNLISAAKAMDSSSVNGCSRVCTEDSLNYCPKPRIKASSAAARAATLSATLVALAVAIASFAY*

>Lp_000095600.1

MSDLRQKLKDLKAPYPEPRPDQCRYVIFLEPKGDEVELNNYKVELMPGRMEKVDGANLHRLGGKIEAKTIEGWGYDYYEVKMGQMASTLMMPLGAAAELKPRFVPMYVDQLYRYNSKLPIVVYMPKDSELRYRIWSATGASEGTPAPKA*

>Lp_270017100.1

MVKETGYYDALGVSPDAGEDEIKRAYRKLALKYHPDKNTDPGAQDKFKEVSVAYEVLSDPEKRQRYDQFGKDAAEAEGGGMDPSDIFAAFFGGGSRPRGEPKPKDIVHELPVPLEAFYCGKTIKLAITRDRLCAQCNGSGSKVAGVSATCKDCNGRGVRMVMRQLQPGFVQQMQVSCPTCKGKGTNLKEEDKCMGCRGQQILKDKKIFEVVVEKGMHRGDSVTFSGEGDQIPGVKLSGDIIIILDQKPHPQFTRKGDHLLMEHTISLAESLTGFSMNVTQLDGRKLRISTTPGVVVDPSNMYSVSREGMPVAKTGGMERGDLIIRFRVVFPKTLEEGSFAELRKVLGYPPQPTAEVRSELHTLQESHIDFEKEARRNAYDDDGDQQPRVQTAGCTQQ*

>Lp_180011400.1

MSIEECKGFDEVKATFAELGQNPTILHHDEKASVDDVLGLLKEQLGITAAGTKTLFLKSKKGELVMVTALREVTTDLKVIQGMTNTKELRFANSEVLHENLKVVQGCVTPLPLINNLDKRNITVLFDQSIATSPIPLAFCVCRNDHTIVVTFEQLKMFMDKIGYPYRMVDFSNPMTSPTSANSSNTKVEKKYADPAVKKAAPAAAATASGETKLGIAVKREDNFSAWYIDVITKAEMIEYYDVSGCYIIRPWAFYVWKCVQRFLGGKIERMGVEDCYFPMFVSRNCLEREKEHIEGFAPEVAWVTRAGDTDLEQPVAVRPTSETVMYPYYAKWIRSHRDLPVRLNMWNNVIRWEFSHPTPFIRTREFLWQEGHCAWAKAEDCAKEVLDILECYAAVYEHLLAVPVVRGKKTEKEKFAGGYYTTTVETYIEAVGRGCQGATSHNLGQNFGKMFDIKFQDPENNEQTLIPWQNSWGLSTRVIGVMIMVHGDNRGMVMPPRVASTQVVIIPVGITKDTTDAEREELLRSCRRLEGELCDSGVRAKCDLRDNYSPGWRFNHWEVKGVPLRVELGPKELAENKLAVAVRFSGEKRSIAWNPQTRTTIGALLEDVQMQMYNRAKATMEAHRVKLTEWSEFVPALNRKCVILAPWCGSTECEEQVKKDSAEESKAAQAQDAREDARAPSMGAKTLCIPFSQPGPVEGHTCICKSCDKPATTWVLFGRSY*

>Lp_000332800.1

MMRYSFLRLAAAPAAAAAAAASSDAKMVSLHKLLIGEVQFRGNAPLKECNIEHNFGANWRADLADYAKGLPAEQKKLLERQVARVGLTRYTTRELSEYCGEGPENVDKVAREANVAQAKAYAAKHGADKLEAYVMAEAKNAGWSEADAKKFIDAVKAAK*

>Lp_000400200.1

MASNCIFCKIVRGEIPCAKVAESAKAFAFMDIGPLARGHVLVIPKAHAACLHELDLESAADVGLLVAKVSRAVAGPEGTTQYNVLQNNGKLAHQEVPHVHFHVIPKRDMATGLQIGWDTIKVSPHDLAEDAKQYKDVIEKL*

>Lp_360006300.1

MVNFTVDEVRALMDYPDQIRNMSVIAHVDHGKSTLSDSLVGAAGIIKMEEAGDKRIMDTRADEIARGITIKSTAISLHYRVPKEMISDLDDDKRDFLINLIDSPGHVDFSSEVTAALRVTDGALVVVDCVEGVCVQTETVLRQALTERIRPVVFINKVDRAVLELQLDPEEAYQGFVKTLQNVNVVIATYNDPCMGDVQVSPEKGTVAIGSGLQAWAFSLTRFANMYAAKFGVDEMKMRERLWGDNFFDAKNKKWIKQETNADGERVRRAFCQFCLDPIYQIFDAVMNEKKDKVDKMLKSLHVSLTPEEREQVPKKLLKTVMMKFLPAAETLLQMIVAHLPSPKKAQTYRAEMLYSGEPSEEDKYFMGIKNCDPKAPLMLYISKMVPTADRGRFFAFGRIFAGTVKSGQKVRIMGNNYVYGKKQDLYDDKPVQRTVLMMGRYQEAVEDMPCGNVVGLVGVDKYIVKSATITDDGENPHPLRDMKYSVSPVVRVAVEAKNPSDLPKLVEGLKRLAKSDPLVVCSIEESGEHIVAGAGELHLEICLKDLQDDFMNGAPLKISEPVVSFRETVTDVSSQQCLSKSANKHNRLFCRGAPLTEELALAMEEGTAGPEADPKVRARFLADNYEWDVQEARKIWCYGPDNRGPNVVVDVTKGVQNMAEMKDSFVAAWQWATREGVLCDENMRGVRVNVEDVVMHADAIHRGGGQIIPTARRVFYACCLTASPRLMEPMFIVDIQTVENAMGGIYGVLTRRRGVIIGEENRPGTPIYNVRAYLPVAESFGFTGDLRASSAGQAFPQCVFDHWQEYPGDPLEPKSLANTTTLAIRTRKGLKPEIPGLDQFMDKL*

>Lp_000183400.1

MSGLAQYLPNIEKLRRGDSEVDVKSLAGKTVFFYFSASWCPSCRGFTPQLLEFYEKYHVQKNFEVIFCTWDEEEDEFEGYFKKMPWLALPFSQRDVVQNLSKHFHVETIPCFIGVEADSGSVVTTRARQKLVKDPEGEQFPWKDVQ*

>Lp_140012100.1

MAIEKKACNSLLLKINQIGTISESIAASKLCMAHGWSVMVSHRSGETEDTYIADLSVGLGTGQIKTGAPCRGERTAKMNQLLRIEEEIGAAAKYGYPGWA*

>Lp_000052600.1

ATMPAAPKLFVALAVSTVAVFTLCCTLAVGAAHHGYPSCDLSVVQSSGYIDIPGVGGTQKHYFYWLFGPRRWPRDGSRPPVIMWMTGGPGCSSGLALAIELGPCEVNETSGELYRNAYGWNDEAYLLFVDQPTGVGYSYGDKANYVHNESEVAEDMYNFLQGFAKRFLSPSITGGNDFYIIGESYAGHYVPAVSYRILQGNQRGDKPKINLKGIAIGNGFTSLLIQYPFYVTYAYDFCKEKLGTPCVNASTRDEMLSMMPQCLELIRTCNSFPTDGDPSCLAATQHCSIIENLFSRSGLNPMDITKRNVGNLGYSMNHTQAFFADAKLRAQLGVADGAQWSTCNDEVTKLFDKDELRNFDYVIPTILSSGVPVLIYAGDLDYSCNWIGNKAWVTALEWPGKAAFNAAPDVEFSVNGRAAGQERTYGNFSFVRIYDAGHMVPMDQPEVSLYMVSRFLHNKRLA*

>Lp_000333200.1

MGKVELTVCAARKLHDGQLVGLPDPFVRLTTGDKKYKTKVVKNNLNPEWEETFRFNIADEMSTQIRLEVWNKNTYDDDLMGYYTLSLGGLTKGIVKDQWYILEKSKTEAELHVRVLAVDFGAPPKPEEQWMVTTDISRDPVKRAIEDGTWRPGQKMPPPPSGPPQPQAYVQQAPVPVQPQVQYAQQAAPVAQPNTQPVQYVAAPPQQQPIQYVQQPQYMQQQPQYVQQQPQYVQQQPQYAQQVPPQPGYYPQQQPPQPGSYPQQQPPQQQPNQQGYYTQAPPQGYAPQPYYQY*

>Lp_330013200.1

MPFIHTTVSTKLDAAKRANLTSAFKAISTQALSKPANFIMTAFNDESPMSFQDTTEPAACVRVDVYEEYPPNAPAEMTPVITAAVSKECGIPADRIYVFYYSTPYVGWNGSNF*

>Lp_330005100.1

MPITVNTVSGGSITLDASAENMYDFHPGQIVHFTKSLRNGKVALVRGTHDGLIWFAVFNTVADAATKEALDAPVDTVSCRGKEELIRQYGWMIDDTSNPFAQAQFE*

>Lp_000298300.1

MSASGALVPPISWAQRPEYVLLTIALQDTTNVKIEIKEDGKVLHFSCSAPEDKHYEYTLHFYAPILSEESQHVVRPRQIELKLKKKFRTSSEEAAEDEVEWPRLTEEKTKHANITIDWSKWKDEDDEGAGADDLGDFGLGGGEGDAMDGQYSEMLSQLMQAQGQHDAEAHAGLPPGSIPAFGSAADQKATGGAEVVVLLFF

>Lp_000520400.1

MSKDAGALKLYVAAGSPLCDLVEIVAHEKKVAYERVVVGLREEMPQWYKEVNPRETVPMLVVGQRRQVVLESTLIARYVDNISTPAGSLMGTSPLQRHRIEFFLSQVGDFTGAAHDLLRDPLNGEKRKAVEDNAVYVDGLLAANQTTGPYYFDAEFTMADVALVPFLVRLKPAMMYYAGYDVFSKAPHIKALWAAAVQRPSVRETAPTPEQCVEGYRHMVPENAPAMGANGGYVLYGNRLCPFVDRARLACALRRFTPYFVEVALHPQPEWYKYINYRETVPALFTPGGEAVHESQLIANYVDSVATEGAALLPRGDADKEYEVAFFVDNASSFVASMYWFTSNPENNEAKGDLLWAAGELEKQLAKHPFGEGPFFGGAQMNLGDVALLPFLVRTKAFTPELTGGYNLFSEFPLLGKLVEAGLASPEGKEVFAPLKVYLENTKARRAKQH*

>Lp_320020700.1

MSSERTFIAVKPDGVQRGLVGEIIARFERKGFKLVALKMLQPSTEQAQGHYTDLSAKPFFNALVQYFSSGPIVGMVWEGKNVVKSGRMLLGATNPADSLPGTIRGDYAVDVGRNVCHGSDSVESAQREIAFWFKPEELVSWTSHSVGQIYE*

>Lp_000363100.1

MSTTTTVLQAQLPAYGRVRTPYEKELLETARILATRGKGILAADEPNEAYDARFAPMGLENNEANRRKYRVILLETEDLNKYIGGVILSKETVEQQCSNGMMFTEMLRKNGIVSGVNVDGGLRPFYEGAPGEEMTAGMDGYVERVQHYYKLGARFCKWRNVFKIQNGNVSESLIQFNAETQARYAALSQLNGLVPIVEPEVMLEGTHDVETCQRVSEHVWATVFAALHRHGVILEGMFLKPSMVVAGSESGIKCTPEEVALATVQTLSRVVPPAVPAIGFLSGGLSELQSSEYLNAINNAPVARPWHLTFTFGRALQGSALKAWKGKDELIPDARRAFLHRARMNSLASSGKYDPKLESEA*

>Lp_000083100.1

MPPRTRRLIHAAILAVVVTMFAATVQHARATLTAAQQSATLAFLQKFPDEFGALKNSWTGTDYCSWEGIGCYNDDVSIDLYDCNLTGHLPELDNSVDGSQVMVTSIGMSNSPNLVGGFPDSWARLTNLRYLDLSSTGLSGAIPDAWNGMSSLETVKISNTYACKTLPNWNITSLRSIDLSNNAFSGSLPSAWGSMTGLRDVDISGVYPCGCVPGTWTSSVLLHAAATLGSGVSSGDCATVNTCHNNDDDHCLRYNATAAGMDVRMRHTLAFLRKFPEAFETLRDKWTGTDYCSWEGIRCNEFKYVNLSSMGLTGRMPELDSDVDGWHVTVTSIDMSNNPNWSDDFEEDWGKLRHLQFLNLSHTALHDEIPNEWSGMRALQEVYITNTGACKSLPDWTNPSLRTIDFSRNNLQGSLSTTWSQMPALTSVDISGNNFCGCVPGTWTSSVLQNAAEAAGGSLLSSTCATSNACTIAKLRCTAAPVSTTAPTPQPTTPTSAPTSNATLAFLQSFPVAIPGLASSWTGTDYCSWEGISCDANGYVSIDLTGRDLTGHMPSMEDHIDGSQVMVTSIDMSNNAKITDNFRNDWARLSNLRSLDLSHTALRGAIPDAWNGMRSLQSIKVSHTNACKGLPNWNISTLQTVDLSNNKMGGTLSSAWAGMG

>Lp_000167500.1

MSAADFEAAVKYVRSLPKDGPVQMDNNTKLEFYSLYKQATEGDVKGTQPWAVQVEARAKWDAWNARKGMKPEEAKEAYVKRLIAVTAEKGHPWNPA*

>Lp_000078700.1

MTFTFSVISTTGQRISISVLGPDNTVGEIKAKLEETEGIPQKMIMLVHSGRKLEDNATLEECNIHAGVTINMVLALRAG*

>Lp_000347200.1

MSCGAAKLNHPAPEFNETALMPNGTFKKVSLSSYKGKWLVLFFYPMDFTFVCPTEIIQFSENAKRFAKLNTEVISCSCDSEYCHLQWTSVDRKKGGLGAMEIPMLADKTKAIARAYGGLDEETGVAYRGAFIIDPTGKLRQIIINDMPVGRNVEEVVRLVEALQFVEEHGEVCPANWKKSDAAKKTEGDAAKKDH*

>Lp_000083400.1

MAHHLCHTIFAAVLALVVLLLTATAASVLAALTAAQNSSTLAFLQLFSSSIPDLSSSWTGSNWCSWTYLDCTNTSNVTLIIDGAALTGSLPALTSEVTGSSVALHTIALMNMNVTGSFPESWGSLTALRVVNLGNTNLFGTLPRSWNAITGLTSVYAARSGACGNLPNWTHSSMLNLDLSDNYLRGTLPISWATMAKLENLNINGNHLCGCVPESWTARVLEYAAVRSLGLRSHAPNCRSTAKCNAAQECSRAAPDYGDAVAAPVDYAVVAPPPVVVVVTAVFAL*

>Lp_350011300.1

MPSVDPSVHIKDQPIFHIRPPKNWINDPNGPYRDPVTGKIHLYMQYNPNGPLWGDIAWYHVTSDDYVKWTRPESPVAMYADKWYDRWGVYSGTMMNNNHSEPVAIYTCTEPENIQRQCMASPPKSDLVGKRTLNSLVKSARNPILTEDDVPGLVGLGNFRDPTEWWEDPANPGHWLIAFVARINDADGDNAHVVVFSTEDPTFQSGYTFSHSLYVYKYDLDRMFECPDFFSLAPGGEHYLKVSTMPSHRDYVIYGSYQPNATTGKYDFVEDPDRSFTFIDYGPFYASKTFHDPILNRRVMWGWTNDELSDAQIQSQGWSGVQNMLRSVEYDSTEKKLRTYPIPELKGLRASHLVSSRSLALSNGVVTLLAAGTNATRHHEIIVTFTLSSMAPFDGTKYYTDATAPEFGVMFRGNSDFSRYTSVSVKMPATTAAPIADSAQDTAWAPIKVFPTTATDPATNCSAECTKERTCVSWTYTSSPSPTCALYWKTSGRVQNSTAQSGTVNLPQLYMDRTVSGSIGSTQPLMGRAPVKQTNANEVQLHIYVDDSIVEVFKDGGWEGVDDGLLLPAGRCRADVHGTLHKEPRRRCRNGLSGRLHDGHCMGSGSGPERVAQLHEQPLRLFVLLN*

>Lp_000173400.1

MTSAKPNIDALLKPLVIGHHSMANRFVMAPLTRCRSGLEHIPNDEMVKHYSDRASMGLIICEATQIQKGYSTFGREPGIYGPAQVAGWKKVTDAVHAKGGLIYCQIHNGGRATVPCNVSEGLKVIAPSPVAIVGHDSPALFNQSGVKEHYPTPVELTKAEIEEYVQLYATAAHNAIAAGFDGVEIHGANGYLIDQFLKTSSNKRTD

>Lp_000355900.1

MREAICIHIGQAGCQVGNACWELFCLEHGIQPDGSMPSDKSIGVEDDAFNTFFSETGAGKHVPRALYLDLEPTVVDEVRTGTYRQLFNPEQLVSGKEDAANNYARGHYTIGKEIVDLALDRIRKLADNCTGLQGFMVFHAVGGGTGSGLGALLLERLSVDYGKKSKLGYTVYPSPQVSTAVVEPYNCVLSTHSLLEHTDVATMLDNEAIYDLTRRSLDIERPSYTNVNRLIGQVVSSLTASLRFDGALNVDLTEFQTNLVPYPRIHFVLTSYAPVVSAEKAYHEQLSVADITNSVFEPAGMLTKCDPRHGKYMSCCLMYRGDVVPKDVNAAIATIKTKRTIQFVDWCPTGFKCGINYQPPTVVPGGDLAKVQRAVCMIANSTAIAEVFARIDHKFDLMYSKRAFVHWYVGEGMEEGEFSEAREDLAALEKDYEEVGAESADDAGEEDVEEY*

>Lp_000009600.1

MSAYQKNNSVEVRRAECARLQAKYPAHVAMVVEAATSSKAHFLALPRDATVAELEAAVRAEKAMGGVGGPCFASD*

>Lp_000369000.1

MSAYQKNNSVEVRRAECARLQAKYPAHVAMVVEAATSSKAHFLALPRDATVAELEAAVRAALGISAKKMTLAGGGCTPAASATVGDLSDVCTQDDGFLYVAVRAEKAMGAFTGPCYA

>Lp_000038500.1

MIKECDAIIADLSPFRSLEPDCGTAFEVGYGAALGKVLLTYSSDTRTMVEKYGGMEAQGLAVENFDLPFNLMLADGTPVFGSFEAAFEHFLQHHAAT*

>Lp_000068100.1

MLCSLVAALHGGHRGVWTVALFALVLFLSACQLVALAVVDTVPRLTPEQQKRAVTNIINHHSFSPPLLRHYYGDGEIPHWMISGTTVITDNYIRLTANEKSQTGHLWNTEPLDMNAFEITFGFRAFQPFGGMGADGFAIWVAQLPRFDGNLFGRPTNFDGFGILFDSYDNDIRRDNPMVTLVVNDGSTTKKFTPNNDFLGEGLASCVFDYRNIFAPNMATARLRYNKGTLSLFLSRNNEVSEQQCFSASNVDLPVGKSYLAFSGQTGEVAEVHDIIFVHLSPLANTTYDHDVQQPAQDELDAKTQLYDNVAMNNRRSTEPSTQVPLQQQQQQQQQQQQQQQQQQQDIEARIRAEAERRVAELERQRLEAQQQQKAQQQAQQQPTQQPQDAAEQARAEADRRVAELERELAEMKRRENRRPERVIEEDDEDDEVDEDEVDEDDVDGDATQPRRRRRVRARRPRRNTVQYEG*

>Lp_360030900.1

MSAPIDKVKEVAESIEQLIGQTPALYLNKLNHTKAKIVLKMECENPMASVKDRLGLAIYEKAEKEGKLVPGKSVVVEATSGNTGVALAHLGAIRGYKVIITMPESMSLERRCLLRIFGAELILTPAALGMKGAVAMAKKICAANPNAVLADQFSTKYNAQIHEETTGPEIWEQTHHNVDCYVAGVGTGGTLTGVARALKKFGSHARIVAVEPAESPVLSGGKPGPHKIQGIGPGFVPDVLDRSLVDEVFCVKGDDAVETSLKLTRSDGVFCGFSGGANVYAALKIGERPEMEGKTIVTIIPSFGERYLSTALYKSIRDEVSSLPVVDASELQD*

>Lp_000382100.1

MSDSYRLIRNATTIFEYGGKKFLIDPMLAKKGAYPGFEHSANSHLRNPLVELPMRVEDILKGVEAVILTHTHLDHWDEAAVQAIPKSLLFYTQNASDAALLKSQGFTNVQVMGDDGTDLGNSITLHKAACQHGPDDLYAVPPLAEALGQVAGLVFQRPGGKTVYFVADTIWFRGVEDAMKKYKPDFVVLNTGKAKMDGFAPIIMEEEDVVRTLAIVPNAVVVAQHMDAINHCLLSRKQMREFVEAKGVSSHVVIPADGELVPVS*

>Lp_000214300.1

MAWLNHPLRRMAVAAALLVLCVVDASLVGARTASDYTAAQQASTLQFLQGFVTANPSLSAVWTGTDFCSWSYVTCVWYSNSLDFSTSKSPYSGTLVLPELGDDVDGSAVVFMEIKVRSIVVRVTGTLPASWGRLTALETLYLDGNGLTGTLPSAWGGMTKLETVNLSNNSVTGTLPSSWRAMAYMSSLNLSRNAISGFLPPQWASMSSLRYLYLSTNALTGSIPASWGQTWTFLFTAAFEGNKLCGCLPSQWENNNIFVTITVDPAVRAAECAVKNA

>Lp_090007800.1

MSSDNNNDAAAAQPPVAVKKPHRVTFGFVEGEDRGPNPMNPPRYHDDPYFWMRDDARTNPEVIEHLRKEKAYFEARSADTLGLRDEIYAEHISHIKEDDTSAPYVDGPYRYYTREVKGKSYKIHCRVPKDKTPGDAAAEQVIIDVNQVAEGKAFCDVMQVEPAPPTHDLVAYSVDWSGNEVYTIEFKKVSNEAEKVPDVVTGTNGDIVWGKDSTSFFYLTKDETLRDNKVWRHVMGQQQSDDVCLYEEHNTLFSAFIYKSADSNTLCIGSVSSETTEMHLLDLRKGNTYNTLETVRPREKGVRYDVQIHGTEHLLILTNEGGAVNHKLIIASREHPADWSNVLVGHREDVFMENIAVRARYLVVAGRRAGLTRIWTMMVDPQDGVFKAGEGLREVEMDEPIFTVHLVESQMAEYEEPTFRMEYSSLATPNTWFDVKPEDHSRVAVKVREVGGGFAASNYKVERRFATAPDRTKVPLSIVYHKDLDLSKPQPCMLYGYGSYGLCMDPQFNIKHLPYCDRGMIYAVAHIRGGSEMGRAWYEIGAKYLTKRNTFSDFIAAAEYLVESKMTTPAQLACEGRSAGGLLMGAVLNMRPDLFKAALAGVPFVDVMTTMSDPSIPLTTGEWEEWGNPNEYKYYDYMLSYSPIDNVRAQEYPNIMIQSGLHDPRVAYWEPAKWVSKLREYKTDNNEILLNMDMESGHFSAKDRYKFWKESAIQQAFVCKHLKSTVRMIVHK*

>Lp_190008200.1

MVNVCVVGAAGGIGQSLSLLLMRQLPYGSTLSLFDVVGAPGVAADLSHVDNAGVTVKYAAGKVGVKRDPALGELAKGVDLFVIVAGVPRKPGMTRDDLFKINAGIILDLVLTCATSSPKAVFCIVTNPVNSTVAIAAQALKKLGCYDKNRLMGVSLLDGLRATRFINEARKPLTVSQVPVVGGHSDITIVPLFHQLPGPLPDEATLAKIVTRVQVAGTEVVKAKAGRGSATLSMAEAGARFALKVVQGLTGAGNPLVYTYVDTDGQHESEFLAIPVVLGKTGIVKRLPIGRLAPSEEKMLKDALPVIQKNIAKGNEFAKANL*

>Lp_300018800.1

MAPLKVGINGFGRIGRMVFQSMCEGDVLGKEIDVVAVVDMSTDAEYFAYQMKFDTVHGRPKYTVEVAKSSPEVKKPDVLVVNGHRILCVKAQRNPADLPWGKLGVEYVIESTGLFTNKAKAEGHVKGGAKKVVISAPASGGAKTIVMGVNHHEYDPATHHVVSNASCTTNCLAPIVHVLTKENFGIETGLMTTIHSYTATQKTVDGVSLKDWRGGRAAAVNIIPSTTGAAKAVGMVIPSTKGKLTGMSFRVPTPDVSVVDLTFRATRDTSIQEIDAALKKAAKTYMKGILGYTDDELVSSDFINDARSSIYDSKATLQNNLPGEKRFFKVVSWYDNEWGYSHRVVDLVRFMGAKDRASSKM*

>Lp_000211400.1

MQRLTYAAMKALVGKKQAGELPNTYIFDVRGTDEFAGGAIPSAVNVPLDQLGAALKLSTDDFKNQYKVPKPSKADHIVTYCLRGGRAERAAHALAEDGYTNVDIYPGSWTEWSEMEKNA*

>Lp_000277400.1

MAHMDRFMQVYAEVQDFLLGDAVRRFEMDENRKRYLKQMLDATCLGGKYNRGLCVVDVAEAMAKDSQADAATMERVLHDACVCGWMIEMLQAHFLVEDDIMDHSKTRRGKPCWYLHPGVTTQVAINDGLIVLAWATQMALHYFADRPFLAEVMRAFHDVDLTTTIGQLYDVTSMVDSAKLDANVAHANTTDYVEYTTFNHRRIVVYKTAYYTYWLPLMMGLLVSQTAEKVDKTTTHEVAMVMGEYFQVQDDVMDCFTPPEKLGKIGTDIEDAKCSWLAVTFLETAPADKVAVFKANYGFDDQAKVAVIKKLYAEANMLERFAKYEEGVVTKVEKLISALEAQNAAFAGSVRVLWGKTYKRQK*

>Lp_270008000.1

MTAEYDYLFKLLLIGDSGVGKSCLLLRFADDSYTDSYISTIGVDFKIRTLNLDSKVIKLQIWDTAGQERFRTITSSYYRGAHGIIIVYDTTDMESFNNVKTWLSEIEKYASENVNKILVGNKCDLVTKKAVDTQMAQDFADSLGIPFLETSAKNSTNVEEAFIRMASDIKARLAVSGEQAKGAPRPNIGNPQPQKKDDGCC*

>Lp_000070600.1

MSTPSHFKKLVVTSLSKDFRHSTEVVEAHLPDEVPEKMVRVAIKYAGVNASDLNFTNGSYFKNAKLPFDCGFEAVGTVVKVGAGVSNVKEGDVVTVLQYGSFAEFLDAPAQTCVVVPALKPEYIVLPVSAMTAAVALGEVGHPKKGEVALVTAAAGGTGQIAVQLLKHVYGCTVIGTCSSAEKADFLKKIGCDHVINYKTESLDDRLHELAPKGVDVVYECVGGQTFNDALRHIAVHGRVIVIGSISSYKSGQQVPFSHPSGTPLPTLLLVKSASLNGFFLPQFHDVIPKYMKEFLSAVEEGKVHLFVDKKEFKGVSGVADAVDHLYTGSSYGKVVVQIQ*

>Lp_000339200.1

KEVRCGFDELEARTIGTRVSGISRVVQPPLAQRSADAATRLAAWQPIRIRVFTEDLQNPSKYCTKAGDVRPDFQGSTATCTKNDVLTDGKKTVLTELLIPSAIQLHEDRLNVQRVSGSIVVDPSISSDSICGQFTIPASHTTTGVSGADFVLYMSAGPTSGNTIAWAITCQYFGDTGRPAVGASNVSPKYISADPQTVRVIAHELLHALGFSVTTFRARGILSTVSLRGKGASPVLASSNVVARTQAQYGCSTQTFMELEDKGGAGTAYSHWKRRSAKDELMAGISGVGHYTALTIAAMEDTGFYKGNYAKAEPMSYGQNAGCGLATSKCVVNGVSQFSGMFCASTGSDMTCTSDRTAIGHCAIATYTSALPSYFQYFADATRGGGDALMDYCPYIQEYSNTNCTIDTDTLLGNAYGVNSRCFDVPTGFISDGYYIASQLAICASVQCDASTSTYGVKVSGASSYTTCTPGARLSLSSLSSSFWSGTLTCPSYDDVCVGQVDAKQYAKYA

>Lp_000237500.1

MPISTTEAFAKRHVTREDGVDVLPRKMIPVAALEAGYCLSSPIVNEAAAGATYPDQMTAAEFTALCEKNQSAFISAEDMAKAVVVVAPAGVITRGSLEEVMGKSSKKEDFLSEDEVEALFTTLDKDNKGAITAEDFMRALYGDEGAYRLAERRKLDAADAERRKAEAAAQELARKEKEEEEKRNGQAAEAKRKEEERLKADAAKKEAEKAKKEKGNANDEDKGKANDVKGKGTEKKKKASACC*

>Lp_350015900.1

MALRRIQKELKDLEKDPPANTSGGPVNENDLFNWKATIIGPEDSPYAGGLFFLNIHFPSDYPFKPPKLQFTTKIYHPNINNNGGICLDILKDQWSPALTISKVLLSVCSLLTDPNPDDPLVPDIARQYKTDRAAFNKTAAEWTRQYAM*

>Lp_300016600.1

MRPPRIVYSLVIRNRSRQGVTVSVTYTDIGNNKHRERVVVPANSVATVGEKTTKSGMAYFGMEITKVEIDGASVRGSASSLSAPFPAVHSPMKDYPIEIVPKNGALTLVATNLE*

>Lp_000383000.1

MFRFTVPALKKLQPLGQRVLVKRTQAAKQTKAGILIPEQVAGKINEGTVVAVAAASKDWTPTVKVNDTVLLPEFGGSAVKVEGEEFFLYDESMLLGVLTA*

>Lp_000390800.1

MAGLPVATTKQRGGVALMLLAAVVVIVACAAPATAQTATTVPPIIQCPATFEHCKECHTVEGMQICSKCDDTYAPNTMGECAPYNGECEVPNCKICFSDSKTQCTDCNSAFQVTDKFTCEAKSTSSTAGPTTAAPTTAGPTTAAPTTAGPTTAGPTTAAPTTAAPTTPAPTTPAPTTPAPTTPAPTTAAPTTARPTDPPCAVAACALCSIGMPNVCVACTQGYSLMRGGACKPTGSCSVARCAQCFVDDDTVCQTCSKGYSLTVSGGCRRGGHSAAVFHGPTVLVTAIVAAVTYALTSL*

>Lp_000532500.1

MTCYTKSTVCLVALLSVLLATTVSGLYATPKKSPLLQEDFIAEVNKKANGQWTASADNGHLITGRSLDEIKQLMGVRDIRNHALEPRVFMADELARDIPESFDSAENWPQCKTISEIRDQSSCGSCWAIAAAEAMSDRYCTIGKVTDRRISTSNLMSCCFVCGMGCNGGFPSAAWTWWVWVGLTTETCQPYPFAPCAHHTNSSKYPACPSTIYDTPTCNSTCDSSQNEFVKYKGAKSYSVSSEEGYQRELMAGGPFEVALDVYADFTAYKSGVYSHVSGERLGGHAARPVGWGVLNGVKYWKIANSWNSDWGDQGYFLIKRGSNECGIEDSGVSGTPATN*

>Lp_000320700.1

MKQSFLLLAVCVLFLCVVSAEVQVATKSNFDKIVSGDLTLVKFYAPWCGHCKALAPEFEKASTTLKGVATLAEVDCTKETELASRFDIKGYPTLLIFRSGEKTEDYEGPRTAAGIVAYMKAQVGPAVTMVGNAEQLEELKKEDLPLCLVKTASADSALAMTMTKVANSLRTQLNFALVTDAAISPDDAMESVTVYRHGVEREAYAGASPVTVEAAKQFLSEAQLDFFGELGQDSFQTYMEANKAKPLGWFFVDKDTSPELKKEVAAVAKKYRHKVLMSWIDGDKYRQVSTQLGMDKDVKFPAFVIDFERRHHVMPSEAPLTASSVSEFMEKYVQGNTEETLMSESVPEVETVEGLTTIVGKTMAKYADGSKNVFVLFYAPWCGHCKKLHPDFEKMAKELEAKDVIIGKIDATANDFDRTKFIVSGFPTMYFIPAGGKPVSYEGGRSAAEMKAYVLSHMVDTPAGASTSATAAAPPAEDKKEDREENNDDL*

>Lp_000290200.1

MSLFFRRFFLVALLVAVAAVVMANAECDANCKSCVFGVCVSCKSGYYFEDKKCTPCNSEHCAHCEIAGHCISCEPGYKLQYVTGSDNITVNKYGKCVNAASFAGVPTALVAAAVAVVYAAIA*

>Lp_000171300.1

MSLFFRRFFLVALLVAVAAVVVANAECDANCDSCLFGWCIACKKGYYPGANGCTSCNSEHCAHCEAFGRCIACEPGYKLQYVTGSDNITVNKYGKCVNAASFAGVPTALVAAAVAVVYAAIA*

>Lp_000325400.1

MASYVPGDRVWVYIEGEDWWPARVLSDEEFGPRTPGQDIAVQFYGGIDTPASLYELNSHSEAAHICFFETSSEKAVTSNPELEAAIRHATEDADANPLKSATAMTAAPMPAAAAKRAREMDTFNPTSHSSGSAEARSAAAMAGLVHLPGDKLQRLADKITQAVEAQSLTKVRAALCQLDGVDVYLTELEDTKIGIAVGSVLSQPALKPLWPLARAIVSFWARHLPTETLAAIRSVKQQDVTAAGKGGVDGAASPNEPASPVTTQSPLRQRLGSQAVAGGSAFDTVANTPSAATEPKTSSLPPLKSNTFLRNVRQRLDNPDNSTRYDDATVDAVARRIADDITDLDDRQLLLIRLGEPDMDFLRMKLLSGEWTPKKYLEQPMEIFLTEKQKSEEAQRVAAKVKAVEAAANAGLNETELFQCERCGHRKCTYFEMQVRGADEPTTKYIKCLNCKNAWSQE*

>Lp_350027400.1

MSAHQHLVSDFKSGSGSWLPQAQGFDTLQVHGGVRPDPVTGAILTPIYQATTFVQESIEKYQAKGYSYTRSANPTVTVLEQKLCSLENGDYCTAYATGMAATTTAISSVMSAGDHAIVTNCSYGGTNRACRVFFSRLGMEFTFVDMRDPKNVEAAIKPNTKLVISETPANPTLTLIDVAAVSAICKAKGLVHMCDNTFATALIMRPLDLGADITLISTTKYIDGHDMTVGGALVTKSKELDDKVRLTQNILGNAMSPQVAFLQLQTVKTMSMRVTKQSQNAQQIAEFLETHPAVERVVYPGLKSHPQKELADRQHRNNLHGGMLWFEVKGGTEAGRRLMDTVPRPWSLCENLGAAESIITCPSVMTHANMTSEDRMKVGITDGFVRVSCGIEDGNDLVAALKIALDALAQ*

>Lp_000088600.1

MPAQEYNVGCYLLDRLVEIGCGHLFGVPGDYNLRFLDDVTAQQGLEWVGCANELNAAYAADGYARCRRIAAVLTTYGVGELSAVNGIAGAYAERNPVIHIVGGPATEAEHSHKMMHHTLGDGLFQHFSLMSEQISCVTGRLTAENALTEIDRVIRGVLHHKKPGYILLPMDVAIVPAFPPTVRLAPRLAEFSQESLDAFKVAAEEKIKNSKRTSALVGYLCDRFLCHKEAQALVDDAHIPFAHMLLGKGTLNEQSPNYLGCYFGSTCPDSVRLTVEDADVCVMLGVKFHDFGTGYFSQNIHLDHMIDIQPFEATISGRVYPQLPMKDAILAVHEIAMKYCDNWPRNELKPPRFPAPESDAYSMRHFWNEIQDNLMEGDIVVVDQGTSSSAAAGLVLPRNGKCIVQCLWGSIGFSLPAAFGAQLAEPNRRVILSVGDGSAQMTIQELGSFLRHGLHPIVFLVNNDGYVIERVIHGWNEAYNDIASWNWCALAKAMSATENQADTATVKEVGTIAKMMKETQKERSALMFREVMLGRHELPIVSLAWKPN*

>Lp_360019500.1

MSAASEWSVACSAQANRIVNPIRAISDAAKISESAKPLIKLSVGDPTLDEGYLPISRVQQAALQEAVNSCKLGGYCPAVGTDEARAAVAEYWKRYFVNTPTRKAHIMAENVVMTSGGSHAIQLAITAIANPGDNILVPSPGFPHYKTIADTYEIECRFYSLDSEKNWEANADEIKSLCDGRTKLLVMTNPSNPCGSNFSREHVEMLVHTAEELKLPLFSDEIYAGMVFQYEENPSKVFTSVADVDTDVPRVILGGTAKSHLVPGWRVGWLLFVDPAGRGADYYRGIKNEASLIVGPSTLAQASVPAGLLKTPPNYLAHVTSIIEESALTFFKAINESSVWSDPERRMLIATRPQASMYVMVKIVLERFDPAVVSNDVTFFRKLFEQENVQVLPGHVFHAPGYFRVVITRPDAVLEEAARRMIEFCECHRREDKPQTTAEDSADASAAAAAVANDDIENKAEAEAGKPKEEAFEEKDFNDEL*

>Lp_350038000.1

MRRLFASVSVAAAATLRCYATKFYTDSHEWVEQADGDAIIGISTYAQENLGDVVYVSLPQVGDKVSAKDVVGEVESVKATSNVYSPVDGTVTAVNEKLKDEPGLINQSPEDKGWLVKMKCTEIPKGLMDAEAYKKFLE*

>Lp_310011600.1

MSQRTVLVAFVGVLVLCVCLARAEIFFHDEFNSLDGWVQSEHKSDYGKVKLSAGAVHVDAKKEQGLQLSEDAKFYAVSKKLPTPVNNDGKNLVISFSVKHDQKLECGGAYLKFFSELDQKDFNGESPYWLMFGPDACGSMNRVHIILSHDGENHLWKGSMRPTKDQATHVYTLEIAANNSYQLYVDGAYKAGGSLEKDWDIVAPETIPDPDEKKPEDWVDDQMMDDPADTKPADWDDEPATIADPNATKPEDWDDEEDGEWNAPMIPNPAYRGEWSPRQIANPAYKGVWAPKQIPNPNYKPEPNLYRVPAPLQYIGIDVWQVQGGSIFDNIILGDDLQEVLKLVKSTYGAMAEKEKKLLKVIEEEKREEASKAKAEAEKAAAAEEAASEEEEEKEDEDDL*

>Lp_110008600.1

MSTVFKAPEKREELVYTAKIAEQCERHDEILFCMKRVVKMNPKLSSEERNLLSAAYKFIISARRACWRSMSSMAHKEDNHKGKTASLFNGFQQQVEKELAEICSDILELLDKYLIPAADNDESKVYYYKLKGDYHRYFAEVETGAETQKNLALEAYKKASEFTASLKPTSPIRLGLALNFSVFYYEILRSPDKGCQLARQAFEEALSDPEVLDEEQNKESALIMQLLRDNLALWTEDSRPEGQDDGTAMEELE*

>Lp_140007600.1

MADERFDSMLLAIAQQQEGIDGILDTFFSFLSRKTDFFTQPEMARASVQKVMDRYLALAAAQQQQRQAKEAAEVQQRQHEAAGRPTSSRVEVLDDEEEMNAAAAAKAAQAAAERKAQLEKTQQALDEARAKKEAEEQAKKAAAAEGEAEASADTDDAPRGLPPTADNGFAYEKYIFSQSLQEAEVRVPLPATNVRGKQLNVVITADHLTVGMKGQPAIVDGELFAKVRAEECMWTIEDGSTVVVTLYKQNTMEWWKTIFRGDPEIDLQKVMPENSKLDDLDSDTRQTVEKMMYDQRQKMMGKPTSEEQKKQDMLRKFMEAHPEMDFSNAKFS*

>Lp_000476700.1

MREIVSCQAGQCGNQIGSKFWEVISDEHGVDPTGSYQGDSDLQLERINVYFDESTGGRYVPRAVLMDLEPGTMDSVRAGPYGQLFRPDNFIFGQSGAGNNWAKGHYTEGAELIDSVLDVCRKEAESCDCLQGFQLSHSLGGGTGSGMGTLLISKLREEYPDRIMMTFSVIPSPRVSDTVVEPYNTTLSVHQLVENSDESMCIDNEALYDICFRTLKLTTPTFGDLNHLVAAVMSGVTCCLRFPGQLNSDLRKLAVNLVPFPRLHFFMMGFAPLTSRGSQQYRGLSVAELTQQMFDAKNMMQAADPRHGRYLTASALFRGRMSTKEVDEQMLNVQNKNSSYFIEWIPNNIKSSICDIPPKGLKMSVTFIGNNTCIQEMFRRVGEQFTGMFRRKAFLHWYTGEGMDEMEFTEAESNMNDLVSEYQQYQDATVEEEGEFDEEEE

>Lp_000476500.1

MQIFVKTLTGKTIALEVEASDTIENVKAKIQDKEGIPPDQQRLIFAGKQLEEGRTLADYNIQKESTLHLVLRLRGGIMEPTLVALAKKYNWEKKVCRRCYARLPVRATNCRKKACGHCSNLRMKKKLR*

>Lp_000257100.1

MSSADEMFREADKKTKKTFFKDFEGALDLFTKAAAKYKLDKDFMRAGDAYSRAGDCAVRLNDKPGASFAFADAANAYKKVDAAKAKNMLDMAVRLKIENNRLGDAARLLLEFAAGLEEQGSSMEALPYYEQAMKYFDAEDQKAQSQKCMLAMAKIYGENDNFDKSLMYYERVANNMLGGPLKFQAQDYFLRAMLCRFAMVTNDNRFEGSEECRDALQQYLSADIYLKNTRESEFLQLILDAVTDNDVEKFEQGVSLLQDIRKLDDWKTHVLLVVKHNMESLA*

>Lp_360026800.1

MTETIKWKNAAIQDEIVPKQNEIKLPDDLVELVYMAKLAEELERFEEMLQCIRKYVKLNSELDTEERNLLSVAYKNVITPRRNAWRLITSIESRENAKENTATIPLVVNMRTELEAELSALCDDLLSLLDTYLIPAAQGGEAKAFYLKMKGDYHRYYAEIDSGDGQKQAALAAYQKATDVANSSLAPTHPIRLGLALNFSVFYYEIMKEHEKGFQLARQAYDEAVTELETLDDEAYHESNTIVRLLRENLNLWTDDPA*

>Lp_000403100.1

MLAKCFYLLAVLTTAVSFVFSDVTFYPQGTHVQFGDSNLKIRNLVPLKLDWPSTNISSTAEALSFSLDNPFSFDFTVQVFTFFFWMPSKMQVTLTGGAAYTKNGNGACDSQSLVREYCDIKARVVIRGIETNLKIVDVAIDICDVVESMFAPYGQQSFPAVTKGATDISRIIYFQKFAVLGGLSEVLHLPCSSKFVAKNVMEVSTGLVLPLFFDFDTSNNITSGAIDTVTTITNGIAKLLNVSIYNNLTDATTTIAYKGGNVTMTSLKNVLDQIMTKKARAALKVYVPAGASLVYDVVVNDLQCRFFNVICSIPASNGIQVMNSRFTGLGDFNSILGNALGKSIDEMMTNITFGAVKKTSILTGGRYYLPLI*

>Lp_340016800.1

MLRIDFSIAPDAQAALEESGKRTSTAVAIAIVIEKEVLKLLASPVAASGSGLEEDLKATRKVLEAKQPSGAYVIVRTSPSSQYVVVYVSDAASAKERMLYSTGTSRVAGATPHAQKRTLQISSLSELQPSLFAAESKKIREDLMTESERNEAAIARMEVAPQPVALPGVAIKMTAEADEMLTRFAKGSVEVVTYKITSEQLQLDRTMTKLGGDLANVKDLLPEAEPRFVLVRYPSPKTSRAEFVMVYVCPPTCSPKVKIQYASSAAAFREQASRHEIKFVQKVETDTRDTLVEDVKSAFEPFSAGGGSAASQDARPRPPVAGAVKAHRMLI*

>Lp_160011700.1

MFLFFFFLLEELCRLVWSSAPPSSSRLFPSPVCPPCVASLYTLASLRCSFFRARTQFSLPRESHYSHCAKLGCISSKSTQIGERECKTAAERKAAWEGIRERLPRRKTPEDKERRIELFKKFDQNNTNRLTLEEVYEGCVSILHLDEFTTRLRDIVKRAFNKAKEMGTKDRGAGSDKFVEFLEFRLMLCYIYDYFELTVMFDEIDTSGNMLIDAKEFKKAVPRIQEWGVKIEDPDAVFKEIDSNGSGQMTFDEFAAWAAAHKLDADGDPDNMAA*

>Lp_000025400.1

MSAGGAEFKAELRAGKPKFGVFLNSASPLLAGQFSHSGYDWLLIDAQHSPVDSLTVAQMVAAVRVGHAKVMVRVSSTRDRAGIQSSLDSGADGVLIPYVNNAKELEEAVSCCYYPTKGTRSVYQPQQCMNAAGLLGYVPEANKNVVVAFQVETASCIENLEEIMAVKGVDIAFLGQNDLCMSMGLYDGRYVFPQMYFSPELQGATEKLIATAKKNNVILGLFLFGTDRVGEFLEKGFTFISIGCELHHAMTQAATHVKALKEISAAKGKPWTNQPSALV*

>Lp_260006100.1

MRRVILAALLLLGIAWPSTHAWLLGVQREHVHHYDAAKTSNSFQCLDGSKTISFSAVNDDICDCPDGSDEPGTPACATLRNGVVARLPDGWLFQCTNKGFSEQKISHSYVNDGICDCCDGSDELNSGVSCPNRCAEVEGELARKMAEEKKRMQTASANKAAMRAEVAKHREELAKTLPLRKAERDRIVAEMPALEEKNNTLQKALEPVREELRVKYAEWEATKEEREAAEEKSSCIKWRQTGKCKADGPREPNQDKDCGIVIPGKDSGFCECAAGNADHASQDDGTNDVADSDSNEESTIHYEFACGHPHLTCMHVCEHNGEAAVGAVYEKPKDPNSYTTPEATAAEEALTARRSKLAELEKTIEESSKILNSTTLSTEELLRTLEGKTFTMEFQDYTYAVTMFKDVYQRGKGQSSGGSLLGEWKSFAENTYAMWGKDAYDLSQMLYDRGARCWNGVTRKVDIQLVCGPENKLTHVEEPSMCIYRMVFETPVMCDD*

>Lp_000131900.1

MPKELEAAIVKQFSSVAEFKTAFEQAGASNFGSGWTWLCVNPKTKGLEIDNTSNAGCPITSGLRPIFTADVWEHAYYKDFENRRPDYLKEMWKVVNWKFVAQMYAQATK*

>Lp_200009600.1

MEDEAVSSNHSSVSDDSQPPMEVDYPLNEEVEVPDTDGGLHKTVLVEGTGARPVKGAKVSVHYVGTLEDGTKFDSSRDRGEYFEFTLGRGQVIKGWDKGVATMRIGEKAILKCSPEYGYGVSGSPPKIPANATLLFEVELFAWSREVDISPQKDKSLMMDVLKDGVEFENPAFESSLTMDLLIYVGKFDPEDKEKKHVPVKVLSNWSIVVGVTPLPPYLEDFLYKMRKQEAAACRVRSDLIHDAVPAFDIPSSADRNHLDVTYVVEVSAMSRSKTYDFTGKPKIAEGEKRKDAGNDAFKAGNLVLAEKYYRRALEFIGEDYGYDDADKPECHRVRISVMGNLAQVLLMRTKYSESAQFSQKVLDLDANNTKALFRLAKAQDGLQEWDAAAKSVDKILAADPNNADAAALKSKIQQAQKAYDQKQKSVFKKMFS*

>Lp_000183300.1

MAGLDKYLPDIEKLRRGDSEVDVMSLAGKTVFFYFSASWCPSCRGFTPQLLEFYEKYHVQKNFEVIFCSWDEEEPGYNVYYAKMPWLALPFSYSDVVQDLTKGFRVESIPTLIGVDADSGRVLTTRARYMLGDDPEGERFPWKDVR*

>Lp_000240000.1

MGKDKVHMNLVVVGHVDAGKSTATGHLIYKCGGIDKRTIEKFEKEAAEMGKASFKYAWVLDKLKAERERGIAIDIALWKFESPKSVFTIIAAPGHRDFIKNMITGTSQADAAILMIDSTQGGFEAGISKDGQTREHALLAFTLGVKQMVVCCNKMDDKTVQYSQARYEEISKEVAAYLKRVGYNPEKVRFIPISGWQGDNMIDKSDNMPWYKGPTLLEALDMLEAPVRPVDKPLRLPLQDVYKMGGIGTVPVGRVETGVMKPGDVVTFAPANVTTEVKSIEMHHEQLAEAVPGDNVGFNVKNVSVKDIRRGNVCGNSKNDPPKEAADFTAQVIVLNHPGQISNGYAPVLDCHTSHIACRFADIESKIDRRSGKELEKNPKAIKSGDAAIVKMVPQKPMCVEVFNDYPPLGRFAVRDMRQTVAVGIIKAVNKKDGSAGKVTKAATKAAKK*

>Lp_000206600.1

MQHVVFFKLDPEKFAAEFPGDSIYEDFETMRRANIPGLLEFNFSAKNTTAWEGYQDATQGYTHAFISRHTDAEALHIFADHPDHKVLQLRVFKCLAAPPLRMELNWHPPAQ*

>Lp_000109400.1

SSIYDFQVNGADHKPYNLAQHKGHPLLIYNVASNCGFTKGGYEAATELYNKYKDRGFTVLAFPCNQFMGQEPGTETEIKQFACSRFKADFPIMEKVNVNGEKEEPVYHYLKNAQRGVLGTTAIKWNFTAFLVDKDGHAVHRFSPGTTAAAIEKELLPLLGGGA

>Lp_000347300.1

MSCGAAKMNHPAPEFNETALMPNGTFKKVSLSSYKGKWLVLFFYPMDFTFVCPTEIIQFSENAKRFAKLNTEVISCSCDSEYCHLQWTSVDRKKGGLGAMEIPMLADKTKAIARAYGVLDEETGVAYRGAFIIDPTGKLRQIIINDMPVGRNVEEVVRLVEALQFVEEHGEVCPANWKPGSATMKPDPKESVSGYFSKL*

>Lp_000460500.1

RSTIMTDLHMTSRPGYTGQLNGRILERVHFLARELDLPHLEEKFPHLKVTDGSWRKVASFPDDADLGEFIDHTQLKADANDAAFVKLCDEAKAHHFKAVAVNCAQVKRCVALLEGSGVRVGGSVSFPLGQTTTAVKVAEAKDEVENGALDMDMMLNVAELKNKNYRYVYEDIKAVCGACPAHVTTKVIFETCLLTEEEIIDRAILCVAAGATFVKTSTGFSTGGATPEAVDVMLAVVGNAALVKAAGGIRDRSTALQYVRAGVRRIGTSAGVAI

>Lp_000355200.1

MSAKNSQEKEPNAFEDKPKSQSDAEENAPQKPQRRQSVLSKAISEHDEEATGPVGDLLPPSKGLTSEEAEELLEKYGRNELPEKKTPSWVIYVRGLWGPMPAALWVAIIIEFSLENWPDGAILFAIQIANATIGWFETIKAGDAVAALKNSLKPVATAFRDGKWQQIDAALLVPGDLVKLAAGSAVPADCSINEGIIDVDEAALTGESLPVTMGPEHMPKMGSNVVRGEVEGTVQLTGAMTFFGKTAALLQSVESDLGNIHVILGRVMISLCAISFVLCMSCFIYLLAKFYETFRRSLQFAVVVLVVSIPIALEIVVTTTLAVGSTHLSKHKIIVTKLSAIEMMSGVNMLCSDKTGTLTLNKMEIQDKCFTFEEGNDLHSTLVLAALAAKWREPPRDALDTMVLGAADLDECDNYEQLEFVPFDPTTKRTAATLVDKRTHEKFNVTKGAPHVILQMVYNQDEINDQVVDIIDTLASRGIRCLSVAKTDQQGRWHMAGILTFLDPPRPDTKETIRRSKEYGVDVKMITGDHVLIAKEMCRMLDLDPNILTADKLPKVKDANDLPDDLGEKYGDMMLSVGGFAQVFPEHKFMIVEALRQRGFTCAMTGDGVNDAPALKRADVGIAVHGATDAARAAADMVLTEPGLSVVVEAMLVSREVFQRMLSFLTYRISATLQLVCFFFIACFSLTPKDYGSTDPEFQFFHLPVLMFMLITLLNDGCLMTIGYDHVVPSERPQKWNLPVVFVSASILSAVACGSSLMLLWVGLEGYGPVHYYNSWFHHMGLAQLPQGKLVTLMYLKISISDFLTLFSSRTGGNFFFYMAPSPVLFCGAIISLLVSTMAAAFWHKTRPDSVLTEGLAWGDSNAEKLLPLWVWIYCVVWWLVQDIVKVIAHHFMDWIDLFGCVSDTAGSGPIKPYSDGVNEEEAKKHAEKTGAAQLVSKSGTEKVAPEKKDVSPKAGSAGSSPKNALEDLNAHEGEDSVLFPSCLLKSCEFKAAND*

>Lp_360033900.1

MSAASALPSKLEGFVRTEFEDTCRRRFFYGLAFDPYGGTAGLYDLGPTMCAMKSNMLQFWRQHFVLEESMCEVDTTCLTPEEVFKASGHVTRFNDVMVRDTVTGECIRADKFLEEWGEAQMGKDGVTAEQKDEFLHLVHDAAGMNPAQIKAVMEKHAIKSPKGNPFSDPFPFNLMFATHIGPEGDRVGYMRPELAQGIILNFKRLMDSGNAQRMPFAGACIGTAFRNEIAPRAALIRVREFTLAEIEHFVNPNDKSHEKFASVRDVEVWAWSRVRQANNEEPTRMKIGEAVDKKIIDNETLGYFIGRVALFLESIGVRFYRFRQHQTTEMAHYAQDCWDAELLTSYGWIECVGIADRSAYDLTQHSNASKKDLCAREEYDEPRVERQLLRNLTKGLIGKTFGKKAADVMAYLNTAPAAACEAIRAAHAAGEVATVKLPSGEEVSITDKMVAFEEKDVKVTGYSYTPSVIEPSFGVGRILYCLLEQSYWVRRDEASGKNDKRAVFSFTPLLAPQKVALLPLMVKPEAMSTIAELRAELVAHGVSVRVDDSSVTIGKKYARVDELGIPFAITCDFEDDGAVTLRERDSASQVRVPLDEVASVVVELCNPRKPRSWESVVAQYPAQGTADAEKRNLFERVMDDSYGL*

>Lp_350014900.1

MKSVFISGGNRGIGLETARQMGKLGYHVVISARKEDAAKKAVETLSQDGVKADYVIMDTVDTSSVAKAAAEVSKLVNGSLDALINNAGYRAPLGDKEQADLEALRKCFEINVVGTANVTNHLLELVKKAPEGRIVNVGSIMGSCATALAGFEDAPYNISKAALNMYTVNLAAALKGTHVKVNCGHPGWVKTDMGGEGAQLEVSEGAETNVYLATLPADGPTGGYFHKKNRLPW*

>Lp_000482300.1

MGIPLPKPVMTQLQERYGNAIFRCGSNCVNGYRETMEDAHLTYMTDTWGFFGVFDGHVNDQCSEYLEGAWRKAIEKEPIPMTDDRMKEMALKIDQDWMDSGREGGSTGTFFVAQKNGNKVHLQVGNVGDSRVVACIDGVCVPLTEDHKPNNEEERRRIENCGGRVENNRVDGSLAVSRAFGDREYKLGDAGQLDQKVIALADVQHKDVTFNSNDFVLLCCDGVFEGNFPNEEVVRYVKEQLENCNDLAEVAGRVCEEAIERGSRDNISCVIVQFRDGSDYAAEPHSTVIPGPFSVPRNSGFRKAYEMMAEKGNTTVGALLEKRYDTLKNAEALTPEETEELDQFEEGPDAKLTGADRQEWFTNYFRKLCESSANGNSDQMERLQSLQQQAGIPLSILLSLMGEQTQ*

>Lp_240008900.1

MSAKPQPIAAANWKCNGSLASIEKLLDVFNAQEINHDVQCVVAPTFVHIPLVQAKLRNPKYVISAENAIAKSGAFTGEVSMPILKDLGVNWLILGHSERRTYYGETNEIVAQKVAAAVKQGFMVIACIGETLQQREANQTAKVVLTQTAAIAKLVPKEAWAQIVLAYEPVWAIGTGKVATPEQAQEVHALLRQWVKEKIGTDVAGKLRILYGGSVNAGNAKELYAKQDINGFLVGGASLKPEFREIINATQ*

>Lp_000347400.1

MVRSHPAPSFNHSHGHASAPPPKPAEPMHANNSAPHPPPPPPPRTEVTNIYVQRPAPAGGGGSGMLGTMAAVAGGSVIGHGISNYLYGNNNQAPTQPAEAQQLAQAAKQENNACAPQLMGYSKCLEANSESADSCKWAWDYFLQCREENPSAPAQ*

>Lp_050008300.1

MSHAYDLVVIGAGSGGLEAGWDAAALHKKKVAVIDLQKHHGPPYFSALGGTCVNVGCVPKKLMVTGAGYMDTIRESAGFGWELDRDSVKPNWQTLIAAKNKAVSDINKSYEGMFADTEGLSFHQGFGAIQDRNTVVVRKSDDPSSEVLETLDTEYILIATGSWPQRLGIPGDELCITSNEAFYLENAPKRALCVGGGYISVEFAGIFNAYKPRGGQVDLVYRKEVVLRGFDLEVRKQLTEQLTANGIHFRTSDNPAKVTKNDDGTKHVVFESGAEGDYDVVMLAIGRVPRSHALQLDKVGVEVEKNGTVKVDKYSKTNVDNIYAIGDVTNRIIMLTPVAINEGAAFVDTVFGGKPRATDHTKVACAVFSIPPIGTCGYIEEDAAKKYDEVAVYESSFTPLLHNVSGSTYKKFMIRIVTNHADGEVLGVHMLGDNAPEIIQSVGICMKMGAKISDFYNTIGVHPTSAEELCSMRTPAYLYEKGKRVEKLSSNL*

>Lp_000531500.1

MGLLTLLRKLKKSDTEPRILILGLDNAGKTSILRKLSDEDPTTTQATQGFNIKSINCEGFKLNMWDIGGQKAIRAYWPNYYDEVDCLIYVVDAADRRRLEETSAELDTLLQEEKLREAPVLIFSNKCDLATALSPEDVSTALNLHSLRDRTWSIQKCSAKTGEGLHEGLEWAVKNLKPKK*

>Lp_000479500.1

MYASPHPEPVVIAVIGGSGVYKLDCLEDQKLYDMTTPFGKPSDQICVAKVSGVTCAFLPRHGMHHQHNPSEVNYRANICALKQLGVRYIIAINAAGSLDEKYAPGDLVVCDQLIDRTFNRASTFFDNGIVAHVDFAHPMSRTLRTVTLEAMRECFPEAWEGKSKFKIHSTGTIVTMEGPQFSTKAESLANKQLGGHLIGMTTSTEAKLAREAEIAYMTVAMVTDMDAWSDAPHVDTVQVMKVVAANVEKAQKYPPAIIKALSENLYDDPAHHALASAIMTKPEALTPEIRQRMAPIIAKKYPKYAPQE*

>Lp_340006500.1

MRCSQKFLTRVAVLGANGGIGQPLSLLLKNNKYVTELKLYDVKGAPGVAADLSHINTDAKVTGYTAEDLAKAVADVDLVLIPAGVPRKPGMTRDDLFNTNAGIVRDLATAVGKASPKAIIGIISNPVNSTVPVAAEALKKVGCHDPRKVFGITTLDHVRARTFVGAAIGKSPATLRIPVIGGHSGETIVPLLSEFKQLSEEQVQQLTHRIQFGGDEVVKAKAGAGSATLSMAYAGNEWATAVLRALNGEKGVVECTYVESKAEPSCAFFSSPVQLGKDGVEKILPMPTLNAYEQKLLAKCIEGLHGNIKKGVVFGSK*

>Lp_000065200.1

MPQLDNIDGSQVMVTSIDMSGNPNWSDDFEESWARLTNLRYLDLSNTALKGDIPDSWTGMSSLETVKISNTYACKTLPNWNSGSLPSSWGSIEDLDEVDITGNNFCGCVPS

>Lp_000047100.1

MAAFPTRRCAIALLCAALVLVAACARANSIVVNGDVSQSDNAFDDNSITFDLGSVSADVVTVQLINSKVSGSGLSIVGYEDSVPSSVTTRVSMSVTSTTVTQSTIAFTGVMPPNSDIRLTATTATLATAQSLFDFSGLTLSGNVTVTVEDSSVAWPSGSTNTGSIVTYTAGATNIGISNKGALFILNATAVNGASVLHIATSSVFSITDSGVLAVDYGGCDGCSSALVTIDVPLKVDGTSMFRIMHGVVGNGAKGLLASTGYVTVSGQSLYLISDSTIDSGSFFDYHVSGNANDSTAFPFTVTSSTVSFLNLVGPSLGIPDGAYVPSTADSSSTVNGGGCTIGGTALTDTSGYLSKGLKVTQVVNSNGAAGGTCANANCVPGYSSSGAAEADGVACSCICTAKIYNPPSCTTVSDPTQNYHSATCSLANCATCSLLYPSSRCAQCNSGYVLSASYQCELDNPDNATTTTTTTTTTAQPTITSTALCSVAYCEKCSPTDGSTCTSCRNGYKLTSGACIANLNGAAAAQSALLAAVACAAAAAFYVL*

>Lp_320024000.1

MFRRNAARLASFDVTVIGGGPGGYVAAIKAAQLGLKTACVEKRGALGGTCLNVGCIPSKALLFATHMYHDAHANFAKYGIRGGENVTMDVAAMQVQKTKAVKALTSGVEYLFKKNKVTYYKGEGSFVSANSLKVKGLDGKEESIESKKTIVATGSEPTSLPFLPFDEKVVLSSTGALDLDHIPKKMIVIGGGVIGLELGSVWARLGTEVTVVEFAPRCAATADADVSKALTDALAKHEKMKFMVNTKVVSGTNNGSSVSVEVEGKDGKKQTLEADVLLCSVGRRPHTTGLNAEAINLKMERGFVCINDHFETNVPNVYAIGDVVNKGPMLAHKAEDEGVACAELLAGKPGHVNYDAIPAVIYTNPEVAQVGKTEEQVKKEGIDYKVGKFPFSANSRAKAVGTEDGFVKVVTDKKTDKILGVQIVCTSAGEMIAEPTLAMEYGASSEDVGRTCHAHPTMSEAVKEACMACFAQTINF*

>Lp_170007500.1

MLMRHPTLYGKYTVVKFDNVHFEPHATLKFSAMEGSKVRVHGTVSNTFNGVLTRVNGTLKGPVMSTMMYGSERQMALESALMDGFDSGMTYKVKDDGVTLTTRSHTIILQRQ*

>Lp_000252600.1

MFARRVCGTAAATAACLVRRASDKVQGDVIGVDLGTTYSCVATMDGDKARVLENTEGFRTTPSVVAFKGNEKLVGLAAKRQAITNPQSTFYAVKRLIGRRYDDEHIKADIKNVAYKIVRAANGDAWVQDGNGKQYSPSQIGAFVLEKMKETAENFLGHKVSNAVVTCPAYFNDAQRQATKDAGTIAGLNVIRVVNEPTAAALAYGMDKTKDSLIAVYDLGGGTFDISVLEIAGGVFEVKATNGDTHLGGEDFDLALSNYILDEFRKNTGIDLSKERMALQRVREAAEKAKCELSSAMETEINLPFITANAEGPQHIQMHVSRSKFEGITENLIQRSIAPCKQCMKDAGVELKEINDVVLVGGMTRMPKVIETVKTFFGREPFRGVNPDEAVALGAATLGGVLRGDVKGLVLLDVTPLSLGIETLGGVFTRMIPKNTTIPTKKSQTFSTAADNQTQVGIKVFQGEREMAADNQLMGQFDLVGIPPAPRGVPQIEVTFDIDANGICHVTAKDKATGKTQNITITANGGLSKEQIEQMIRDSEAHAESDRVKRELVEVRNNAETQLTTAEKQLGEWKYVSDAEKENVRTLVAELRKVMENPNASKDDLSAATDKLQKAVMECGRTEYQQAAAASSGSSSSSNSGEQQQQQQQQQQQQQQNNEEKK*

>Lp_000252700.1

MFARRVCGTAAATAACLVRRASDKVQGDVIGVDLGTTYSCVATMDGDKARVLENTEGFRTTPSVVAFKGNEKLVGLAAKRQAITNPQSTFYAVKRLIGRRYDDEHIKADIKNVAYKIVRAANGDAWVQDGNGKQYSPSQIGAFVLEKMKETAENFLGRKVSNAVVTCPAYFNDAQRQATKDAGTIAGLNVIRVVNEPTAAALAYGMDKTKDSLIAVYDLGGGTFDISVLEIAGGVFEVKATNGDTHLGGEDFDLALSNYILDEFRKNTGIDLSKERMALQRVREAAEKAKCELSSAMETEINLPFITANAEGPQHIQMHVSRSKFEGITENLIQRSIAPCKQCMKDAGVELKEINDVVLVGGMTRMPKVIETVKTFFGREPFRGVNPDEAVALGAATLGGVLRGDVKGLVLLDVTPLSLGIETLGGVFTRMIPKNTTIPTKKSQTFSTAADNQTQVGIKVFQGEREMAADNQLMGQFDLVGIPPAPRGVPQIEVTFDIDANGICHVTAKDKATGKTQNITITANGGLSKEQIEQMIRDSEAHAESDRVKRELVEVRNNAETQLTTAEKQLGEWKYVSDAEKENVRTLVAELRKVMENPNASKDDLSAATDKLQKAVMECGRTEYQQAAAANSGSSSSSSSGEQQQQQQQQQNNEEKK*

>Lp_340028900.1

MPAQEYNVGCYLLDRLVQIGCGHLFGVPGDYNLRFLDDVTAQQGLEWVGCANELNAAYAADGYARCRRIAAVLTTYGVGELSAVNGIAGAYAERNPVIHIVGGPSVKAERARLVLHHTLGDGDFEHFVRMASEVSCAVCHLNDTNALTEIDRVIAACLHHKKPGYILLPLDVALVPCAAPSAPLSRLEVPRSASILAAFKAAVDKSFQTSKAPAALSGHFCDRFECCKEAQALVDDAHIPFASIMFGKGVLNEQSPNFIGTYYGKPSLEHVRGSIEDADVLVKLGVRFHDFGTGFFSQKIEESHCVDIQPFESSVAGVVYSPLPMKDAILAVHEIAVKYCDNWPRNELKPPRFPAPESDAYSMRHFWNEVQDNLTEGDIVVVDQGTSSSAAAGLVLPRNGKCIVQCLWGSIGFSLPAAFGAQLAEPNRRVILVVGDGSAQMTVQELGSFLRHGLHPIVFLVNNDGYVIERVIHGWNEAYNDIASWNWCMVMKGICNGKAASVEVLKDVNMTAKLVTSHAAPEKALVFREVMLGRHELPIVSLASW*

>Lp_000045700.1

MTETFAFQAEINQLMSLIINTFYSNKEIFLRELISNASDACDKIRYQSLTDPSVLGDETHLRIRVIPDKANKTLTVEDNGIGMTKADLVNNLGTIARSGTKAFMEALEAGGDMSMIGQFGVGFYSAYLVADRVTVVSKNNADEAYVWESSAGGTFTITSAPESDLKRGTRITLHLKEDQQEYLEERRIKELIKKHSEFIGYDIELLVEKTTEKEVTDEDEEEKKEGENEEEPKVEEVKEGEEKKKTKKVKEVTKEFEIQNKHKPLWTRDPKDVTKEEYAAFYKAISNDWEDPAATKHFSVEGQLEFRSILFVPKRAPFDMFEPNKKRNNIKLYVRRVFIMDNCEDLCPDWLGFVKGVVDSEDLPLNISRENLQQNKILKVIRKNIVKKCLEMFDELAENKEDFKQFYEQFSKNLKLGIHEDTANRKKLMELLRFYSTESGEEMTTLKDYVTRMKPEQKSIYYITGDSKKKLESSPFIEEAKRRGIEVLFMTEPIDEYVMQQVKDFEDKKFACLTKEGVHFEETEEEKKKREEEKAAYEKLCKAMKEILGDKVEKVAISERLSTSPCILVTSEFGWSAHMEQIMRNQALRDSSMAQYMMSKKTMELNPHHPIIKELRRRVDADENDKAVKDLVFLLFDTSLLTSGFQLDDPTGYAERINRMIKLGLSLDDEEEAAPAEAAPAAEAAPAEATAGTSSMEQVD*

>Lp_000029300.1

MGCGASSENANVTYLNGKPTFKGDDVTKGFEKDNGLLFRIVNKKKKQWAYYNDTKQYEMHVTVTFNEDCDIKPLGKTRLEQQDNGEWVATVVVYPCETEMFIEGRVNGFRSKMDALPLSDEYRQRQEEKEKK*

>Lp_000299400.1

MGGWLSSLLGKKEVRILMVGLDAAGKTTILYKLKLGEVVTTIPTIGFNVETLEYKNLKFTMWDVGGQDKLRPLWRHYYQNTNGIIFVVDSNDRDRVKDAKAELDKMLVEDELRNAALLVFANKQDLPNAMSTTEVTEKLGLHALRQRNWYIQGCCGTTAQGLYEGLDWLSANIKKTMN*

>Lp_000491400.1

MSVIITKQRYNWMGYNGTETEEVDTALLRGENGTLDNYNYDHRQQIIAELKERLYHIKRDLDLLPEEEMRAEEEARQRRRDERKRREEEARRRRAEEEAARAEEEERQRIEDEERAEERRRQQEEDEQRRREEDEARRNAERWRQPRGDSDDDSDVEEEQNEALSAMPRTQEEIDYAVSGFVKGTRMLLIANDQGYVSYVDGSMHDSIDKLEAYNKEARAAAEAAVAGSSVSVPPAADFSLDDVRTSAARQQELRKKLDAYRAQQDPTYSARVRASEKNSNGQPSSAITQDDGDEAAITARLEEAKQHAEEVRKSYNAKATASGNAIAAVMASRAGAAPVSNKAAGSEAAVNDLADVDHAATAGLSAEETMVADEAPLSEKMPTRTATTASKHSSKKNTPMEDQDEL*

>Lp_290009100.1

MSGLAQYLPNIEKLRRGDSEVDVKSLAGKTVFFYFSASWCPSCRGFTPQLVEFYEKYHVQKNFEVIFCSWDEEEPGYNVYYAKMPWLALPFSYSDVVQDLTKGFRVESIPTLIGVDADSGRVLTTRARYMLGDDPEGERFPWKDVR*

>Lp_220008600.1

MAALTDEQIREAFNLFDADGSGAIDAEEMALAMKGLGFGDLPRDEVERIIRSMNTDANGLVEYGEFESMIKSRMAQKDSPEEILKAFQLFDLDKKGKISFANLKEVAKLLGENPGDDVLKEMIAEADEDGDGEVSFEEFKNVMLQMRGK*

>Lp_270008500.1

MFFTQIFRRKTSTMSTATVAELIPQHKVVMFSWVHCPYCVRAKEILKPLVNDMQVYEVDEMENGENLRKQIYDTYKHETVPAIFINGEFVGGCSDLQALQKSGELAKKLG*

>Lp_000331500.1

MAPTITVEIPYDRLIKDDCISDEYLINQLDGVNDNPPEDNLPLRKWLIREAHEALMKNHKMKEITLKPKSDKSSHMTFTLKVTGNE*

>Lp_130016600.1

MVEVPRNFRLLEELEAGEKGTGSNQNVSVGLRDTADIYFHYWNGTIVGPPSSTFEYRILSLEIYCDENYPKVPPHIRFLSKVNLPCVDPDGTVNRNKFLLFKNWDRRTTMETCLSELRKEMAAPQNRKLPQPPEGSTYS*

>Lp_240011100.1

MKIFKDVLTGAEVVCDNDRPFDVEGDIVYVVKGRYIDVGGEDYGISANVDEDAAEGATGDVAEGKERVVDVVYNNRYTETSYDKASYMAHIRGYMKQLLEKIESADEKKAFQTNAAAFVKKVLKDIEDYQFFIPEGNDEDPDNGMIVLCRWDGETPLFYYWKDGLKGERV*

>Lp_000294500.1

MAASLESLTHIITETPESEASFYSLAYVAALQQCTAWAYQATLTAVTRSTRSSGVCGDSLTECVTRLRVETERRVASQLRTSLLQEQAAAVRAWKRDGRAYTNIEADVVATPRDNGGAAGAAAAAAAVAAGPSPARHRASSVVGGGSSNSAAKQQSRRVSQRTASSSVAGPHNPLQPDALAGLPVGKRGSKETVNAAGAASSSPSSSEASVAAAVKLLREGLARSLGCPLPSSSLFADGKLQGGKSEGCSSTNAQLKGLADTVLSRRSYSPTPAHQGATLFVELEGEEGRQDLVLEWIEDVLIFALRLGFNAPQTQCLLLDSMLALQTLEDEGADNDADDDDAAWEQRVSAALEDVLCTQTCPTTTRVIDTMWQRQPVVREVPDPVQLAELEQKRLKATTQKALAALDAAQASIPLVPRTFMEDVSVEVERDATLPPVFSMAEAAAVVDFVTRSVATHRRLWRLLLTATLPSLRPVVERPLCAYVEDVPPVFLPPLSSFYPARLQDEVDAQRAVYAECAAEKASLYAALYETPLAALSMEAAATRQALMSAAQRAAADDRDVALTTQECAHVAKAFQLRLQDTVQHNTHVVRADTTASSSVLAGSHKDESSGGDAAASLANALSAAAPEPAKHKAKKDSGGTAAVHAAKNPISASNLMRQSMLSADAAAAIPLPTSAVFTLAEVEQRVDRIAVVAEHLPSVGASGVAGRGGQRKH*

>Lp_000438900.1

MAKVKLNFRITGKSITYEGLPEKRPKDLMDILAKMGVSEDVVLPTFRENIETEADYTDEQVMEMFAEAKAIREEKKRAKSGAAAPAAAAAAAATPTQANAEDQEQPQKITLSYAGKQFSYQGLPSGKRNAVLQLLSKQKDVTPEMAVDTYRTNLPEDAATEEEIWEMVQAMKNKKSRHHKDANNGGEAAAAATTTPKTPEAMPAPASSPAGYESNASYDAIDELAGMSSKQRDDLNNFIRQVLEGPTSDVAALLREDPKPRNEAEDHKAEVLTARTSGKIPVYLQEEHNTSGTKLIYATASMTYEEFLAVVEKKFGRKMALSFYEGDDLIELDDDDVLAMYLEMTVNGDPKKLRLICSNPDTRPKEVDDQITETQQTGTVAGSQVITTKAKAYSNGELTVTENRTYSGHSLAVYCCAFAPKGDLFVTASRDRSVRVWNVRTGSCMVMKGGHNGFVLSCDFSPKGNRVVSSSDDRTIKLWNTSSFSKVATLKGHEDKVYCVKYNPSGEYVASASCDHTVRVWNADTQSKVATLKGHTLAVFSCSFSNTDNGKYVVSGSDDRIIKIWDWAASKEVKSLVGHIGTVWTVAYSHNDKYIVSGSMDYELMLWDVETGTRLRSMEGHKTSVHHAIFSEDDKYVFSCARDWSVMVWRTADAEHVETITGHLSTVYHMDIKGDKLLTSSLDDKLKLWTVKRN*

>Lp_000067300.1

MKNVVVDPPYHALLLEGVNPAAKELLESKGCVVETLPSALGRDLLLEKIKDVHFLGIRSKTKVTKEILDAAPKLLAIGCFCIGTNQVDLDHANKRGVVVFNSPFANTRSVAELIIGEIISLSRKMTQRSEEVHRGVWNKAHVGCYEVRGKTLGIVGYGHIGSQVGVLAEALGMNVIFYDVVPTLIIGNATKFSHINDLLTVSDFVTIHVPETETTKGMFGEEQIRLMKQGAYLLNASRGTVVDLDALAKALRDGHLSGAAIDVYPEEPGSNKELHKTPLQGIPNVILTPHIGGSTCEAQAAIGTEVGSALAQFVTNGTTAGAVNFPQLVPPPVDKSNFRITNVHLNIPGALKDINKIAVDLGCNIGMQFLSTHKAIGYLIMDVDKDVAAELRTRIAALDCSLRTLIIR*

>Lp_340015400.1

MSSNKAPFTRLLSSIGGNYSVIDAAYPSSGNKLVNLLTGSADAKQTLSGSQSILRRLHDEKKKVVVAAPASYWSLGPAGTATCPQVGLLDTECSGSACPDEKEAAYCNAARKYITCDDRAQLYQNEIPTAFEKTISSSADMLYFQVSSIAEAASSDVEAAAQERSEVNLLDAAVGRIALALSKRTANTAENWLMIVTSDGSNTEARAPLLVVAYTKGEIVRLNAIAADARVTDVANTVKMWFQLRGVDQNRLLGICTNGAEVKNCNKTA*

>Lp_000158900.1

MAWLNHPLRRMAVAAALLVLCVVDASLVGARTASDYTAAQQASTLQFLQGFVTANPSLSAVWTGTDFCSWSYVTCSYSSDKLDFSSSKSPYSGTLVLPELGDDVNGSAVIFTDIKVRSMGKGAMVAGTLPSSWGRLTKLQTLYLDGNGLTGTLPSAWGGMASLRTLNVSGSRLSGSVPASWGGLWQLRSVDLTGTGLCGCVPAEWAGKAVAADAALTGSDCAVANACSKVVSGSQSESSSRKTKSG

>Lp_000410600.1

MQEWHLPKGGSGVASLTLHKDVPIPTPTGSQMVVKMKALSLNYRDAFIADTSFLPDTIDNVIPISDGAGEVTAVGPACTRFKVGDRVAFIITPNNYEGDQMEGFGLQPCTGGSRQGVAAEYVLAESGDAVRIPDYMTYEEAAALPCAGVTAWRALNSGERRLVPGSTVLVQGTGGVSSLAAQLAHAAGCRVIALSSSEAKMKDLLRLGVVRPGDWVNYRENPKWGEAVRALTPGGHGVDVVVEVVGLSSINESLACARRRGVVSVVGHLDTTPTEDFHLHEVLMKRLQIVGVQVGPRRDFEDMLQLMAVKAIHPVLDPKRFTLSQLKEAYEYLKSQKHSGKIVVTV*

>Lp_000350800.1

MRRFLSKCTAPAVARLASTAAAASAAPGQKSFFKATEMIGYVHSIDGTIATLIPAPGNPGVAYNTIIMIQVSPTTFAAGLVFNLEKDGRIGIILMDNITEVQSGQKVMATGKLLYIPVGSNVLGKVVNPLGHEVPVGLFTRSRALLDSEQVVGKVDTGAPNIVSRSPVNYNLLTGFKAVDTMIPIGRGQRELIVGDRQTGKTSIAVSTIINQVRNNQQILSKNAVISIYVSIGQRCSNVARIHRLLRAYGALRYTTVMAATAAEPAGLQYLAPYSGVTMGEYFMNRGRHCLCVYDDLSKQAVAYRQISLLLRRPPGREAYPGDVFYLHSRLLERAAMLSPGKGGGSVTALPIVETLSNDVTAYIVTNVISITDGQIYLDTKLFTGGQRPAVNIGLSVSRVGSSAQNVAMKAVAGKLKGILAEYRKLAADSVGGNQVQTVPMIRGARFVALFNQKNPSFFMNALVSLYACLNGYLDDVKVNYAKFYEYLLVNKDLSVMYGTATNKFFYMYVHQLNYVIRFFTLNHPVLNAEVEEMLKQHTHLFLQHYQSKMNAIKNEKEVKALKNLLYSCKRAV*

>Lp_000432400.1

STRSSCPRRTHGIMARLVRLAAGALFATLVLAAFAALGHGGERGPAHNVCIHDKLQRRVVEATAATPHTAAVLSSVGLPYVSLRPDTSTVGTTNPGISMSSTFDKFRITSWTADLSTTDSFCAYVGQTVNNHRGDNVTCTAADILTDAKRYTLVNYLLPEAIQLHKDRLQVKRVKDVWRVDGMNRTVCKEFLVPVFHRTYGVSQTDFMLYVASVPAEDGVAAWAITCQVFANNRPAVGVLNVPAKYINTDYDQYMIHTVAHELAHALGFSGAFFWPMPPWETSVTGIRGKDYAAPVVNGTTVVAKAREHYNCSTAQFMELEDKDPTGQSGSHWKIRNAQDELMAGVAGVGYYTALTMAAFEDLGFYKAVYSKAETMKWGRDAGCSFLDGKCVIDNVTQFPSMFCDKDEFVFRCTPARLDMGACGVYDYYKELPSYFQYFTVPNVGGSSEYSDCCPTVEPWDFGGCTQNASDAQSLVRYFNTFSMSSRCIDGNFIPKTTTTAQITAHYGMCTNLACNPANETYSIQVYGNTTYIPCTPGQTIALDSVSDAFEAGGHITCPPYLEVCQGNVQAIRD

>Lp_120005100.1

MPCNCRQGTALHNHLEDQLSSDGVFGLGAPINDAVDLTALQVWNCKNSAEEAQRVFNPNMGDNTLPICSDDDQELLVFAPLSSVCRIKGLSITGPANDCAPSRVKIFVNVGDITGFGSVERLVPQEQLQLADGSSEDRIVYRVNPGKFSSVSSLTFFFDESFNNNETDVLRIELFGESTGKSTHQQVATNIVYESRGNPADHRTTEDKKALFEVQ*

>Lp_000406600.1

MKQMKVAFEANKRVYESVLLTFKGVDGFDVYNCSVPFFYRGKHYIYGRVERPNLWASSYVRLFEETEKDVYTVVPGFSLQLEDPYIAKIQNEMIFGGTHVRKDKGEISTYYGYFYRGTPDELTYFTTGPDYMKDIRVVQMQDGRLGVFSRPRQGNKASIGFTILNSVDELGADAIAKAPPLNILSEDTWGGVNQAYLLSGGNIGCIGHYSYEDKDANQKPLSVYVNYSFVLDPHTREIDECKIIGTKSCYPPCEPKLPFLADCVFTSGIVMRKDGKVDLYSGVGDRYDRYEGRITIDYPFEGHGTIVGDFVF

>Lp_000511200.1

MKQMKVAFEANKRVYESVLLTFKGVDGFDVYNCSVPFFYRGKHYIYGRVERPNLWASSYVRLFEETEKDVYTVVPGFSLQLEDPYIAKIQNEMIFGGTHVRKDKGEISTYYGYFYRGTPDELTYFTTGPDYMKDIRVVQMQDGRLGVFSRPRQGNKASIGFTILNSVDELGADAIAKAPPLNILSEDTWGGVNQAYLLSGGNIGCIGHYSYEDKDANQKPLSVYVNYSFVLDPHTREIDECKIIGTKSCYPPCEPKLPFLADCVFTSGIVMRKDGKVDLYSGVGEGRITIDYPFEGHGTIVGDFVF

>Lp_000280700.1

MSHLPIKWAERHDRLFITVEASSATDVKVTFTEKTVSITGNGITAKSSEPHELKEELHLAKEIVPADSTFKVLGVSIQICAIKKEEGYWNRLVEEPTKATKGWLSVDWNLWKDEDDEAEENAAAANFGGYGDMGGYGDMGGMDMASMMAAMGGAGGAGGAGHMDCDSDDEESTESEDDKEEENKKPAADLSDLNA*

>Lp_000387500.1

LLLIAAAMTAEARLVVRMVQVAHRHGARGPLVSEDNTAEICGTEYPCGELTVVGIEMLNSVGRFVRARYNDLSVVEQALFPSTRYNSSVVYTRSTGIQRTIQSATAFLRGLFPDGYFYPVVYTQNITTDTVLNTDPVPSGMGIRWLEADIFRTALNPVVDDQLTWDQVQAAARDAYIEGLCQDYNDRAFCASNLHDVGAAFEASGRLPSMPNLEAAMPGLTELSAAWHRYAYDYNKSVELNAIQGAVSQNLAQTMLANINAHKLSPSFKLYEYSAHDTTVGPLAITFGDHTDATMSPPYGVTILVELLQDTESNDWYVRLLRGSPVLQTNGSYAFQLSGIQVHCIDSAGNQYLASTNICPLDSFRRMVDYSRPTVVNGQCTLTPEQYNNMGCPRTIASNAPVPENCFVYRFVCPENACPAGYILATTDYQC

>Lp_000007500.1

MPSPVPVNIHFLEEVGSTMEVGREMIASAAGKPFGIVAAVQTAGRGTGGRTWTSPKGNMYFTLCIPQKNNAAYFKEELVPVLSLVCGLACRRAILEVLHLDASSAKAADAAKALTTKWPNDVIFRHKKIGGTLIENDHDYFLIGMGMNIAVGPKVTDAGREATTINAVADEFGVKHTDPKELASAIWEQFFDICAAPGTTRASVVADFDAVMDKTLKLHKRLPDGRDPEELTAVSLNSWGHLKVRHTDGSTEVLSAEYLF*

>Lp_080010500.1

MTKLTVVCRAVSSALDKQTCELTCIERQPISLAVVEEQHHAYVQVFRDMIAEGYDIELIELPALNDLPDSMFVEDVAMLYNECAVITRPGAPSRRPEVDPIVDTVKKLRPDNYRIVEPGTVDGGDVLYVANSKYVFVGKSTRSNDAAFEQMKAYLGKHGLECVQCVVKGCLHLKSAVSFASEDTVLLNPEWIDAAIFTSRGFKVIEVDPKEPDSANVLSFSAEKDGKALRTIFSPVAFAGTAARIHAYAEKETKAGRPTRQVLLKVDEIAKAEGALTCCSLLSYRA*

>Lp_110005700.1

MGLDIQFFRDPDTAAAVRESERRRYAKPEVVDEIVKIDLKWRRTQFLTEATKKMINTCSKAVGEKKKAKEADGTVSEVPAELLSAAHEGTLDADKVKGLCILQLKELSKALATQVADLGKAAAEQEAERDRLVLTVGNVLHESVPVSNDEDTGNVTVRTFGDVTRRLKMTHVDCMEKLGLMDTSKTVTAMAGGRAFVLRGGLVQLQFALISYAINFLVSRGYELLYPPFFLNKDYMSAVSQLSDFDESLYKVSGDGDEKYLIATSEAPIAAYHSNKWFTELKEPLRYGGVSSCFRKEAGAHGRDMLGIFRVHQFDKIEQFVVCSPRENESWKMLEELIKTSQDFNESLGLPYRVINICSGALNNAAAKKYDLEAWFPGSGAFRELVSCSNCTDYQSRGVNCRFGPNVKGTTANNTKEYCHMLNGTLCAVTRTMCCICENYQTEEGIVIPEVLRPFMMGTEMLKFPAEPAKTEAA*

>Lp_000521900.1

MPTQPKLFEPYAIGQLQLRNRIVVPPLCMYCATDGKMGAFHHMHVGSLATSGAGLVIIEATAVRPEGRISPACCGIWDDETAAAMKAVLDEVRVYSKTPIAIQLAHAGRKASDRRPFDPDEGHLNPGEKDRWGQAGWQPVAPSAIPFDAADHPPVSLSKEEIAEIVKAFADAAVRADKIGLDGVELHFAHGYLACEFLSPLSNKRTDEYGGILENRMRMPLEVFQAVRAAFPKEKPVWVRISATEWMDAEGGWNLNDSIAFCRKLKELGCDAVHVSSGGNSIEQKIPVSVGYQLPFAKAIKEAVGINVIGVGMLTEPGEAEAALATGAADAIAIGRQLLFNPHWPYEAAHKLGATVEAPPQYWSSEPSPKLKIFAK*

>Lp_350032300.1

MGKDQANVKGCRFRVSVALPAGAVVNCADNTGAKNLYVISVKGYHGRLNRLPAAALGDMVMCSVKKGKPELRKKVLNAVIIRQRKSWRRKDGTVIYFEDNAGVIVNPKGEMKGSGIAGPVAKESADLWPKISTHAPAIV*

>Lp_360047500.1

MPSEPSNRCCPTDATAANCVYHPTGEDFYISGPLDSKAGIVIVSDIFGMLPNSKRIADVFAKEGYLAIMPDFFGSKAWDYRKWPPDFEGEEWKTFLAFASDFARHRPRVDHAIQVLRKLGVQKVGIVGMCWGAVLAFQAAADGTVNAVATAHPSFFTPATVKSAKTPVLVMPSQDEPPFDDDEAAVNAHPVEPHVYKRFDKIPHGYFGARYDPDTYTPEQMEEVEEARHMTLDFFKKTLH*

>Lp_360022600.1

MDRLARLTLTAALVVVLVELSVARAQSDSAAAPDTASSAVAGVDLEVVMVQVLHRHGARSGTPRYNTSLICTEAPCGYLTWAGVDMLLNVGAYLRNRYNNDRTVVNVPLFPSPNYDLDVSYSRSTDVLRTLQSAEIFLRGFFPNTSNLYAAVHTVEDDTDMLLNSNAQPWLKFFYSYNKPLLRTVCNPLTDKLYPDWTEITTLGKEIFLESYCSDYTTRSDCVRLLFDIAAAKMAVGELDQYPLLKANFAKIQNITRVLFAYEYHYNRSDSLMFKQGGRGQPFLKQLVANMDNQIAGTNKYKLMHYSAHDTTLAPVWGTLGDESETAMLPPFAQVLVAELLKSKSTGAMRVRVLRGAPGQSPDTNFAFDWDATWHLQCMNAWGIAYYAQNNVCPYEDFKRYIRWSEPGDARGYCYLDPDFVTIANCPTKNTVYGIPASAVVSSTCRFYRAACPSFACTPGYTLNSVSYQCVCSSDACLNVTGTTKSGTALLAREAVLKAEEEATSQTMTSLSSGLSTAAVVCVSLGTFAAGAVIAVAATVFVCLCTHRRRQESAMKRGSVQLDDVHSDEHVAPEAEVEAP*

>Lp_000428900.1

MSAVGNTVVTLNNGVKMPQFGLGVWQSPVGEATKNAVTWALQAGYRHIDTAAIYKNEADVGAGIRASGVPREQIFVTTKLWNADQGYESTLAAFEESRKKLGLDYVDLYLIHWPRGNAIVAKEGKKYLDSWRAFEKLYAEKKVRAIGVSNFNIHHLEDIFAMCKVAPMVNQIELHPLNNQAELRAYCKSKNIFVEAWSPLGQGKGLTDPTLISIGKKYHKTAAQVILRWDIQHNIVVIPKSVRRERIESNANVFDFELSAEDMAQIDAMNTNTRYGPNPDDADF*

>Lp_000291900.1

MRAVVVLAFLAAVLFTVSSFVSAEPEVTDKVFFDITIGDEPAGRIVMGLFGKEVPKTAENFKQLCTGEKGFGYKNSIFHRVIQNFMIQGGDFTNFDGTGGKSIYGSKFEDENFKVKHFVGALSMANAGANTNGSQFFITTAATPWLDGRHVVFGKVLDGMDIVHRIERSKTNSRDRPLKTVKIVDSGVVA*

>Lp_000466600.1

MPAPAAHTLFGHADLKVDARFLSNLAFMASSDKPVETPMSVGDYAMSEMCVSDSEAADFKMYLDTIRSVGQEDFVSKVVTKLRAQAAQSRDHAVAATADALESEFARLTKVFFLRARSYQVIKISTDEPVGRVVCFPDQVSDDYGNNLDEHNETGIIQSLVDRSHGAEMMPTVYVGHGQGAAMAEVCGCATGRQAIIFDASLLSTPLVYNAVRHKCVQSDPEDVDVPHEFKLIYYRTDATAKKQDAIFKTFHKAVEEVYAEEKAAEEAAAAEASKKGEAKKEDKNDERRKKEEEQQKAALEWFENPEWYQIGGIVSAAASSRFERTGRYNAKALSDMLVDYTASIRPAKDPYKPQS*

>Lp_000161700.1

MPRTETRTAHVRSRFSRCLFTGAVAVLLLLSLHGSSIAGTVRQSVTDHTNTAELTRRDMPPSLSLRSSLGTFAVPKGRTPIVSLQAACRSEVKLYCSGLTSSPLRCLVERFYRERQLGSTSVFSDVCEAWLTARDACLSFVLTHGAELCGAAAATADARECLRQIPAAFLPPKCVSSDYYEGVRLVGKLRQHQIADARLRRIREQ*

>Lp_000458800.1

MTDIDIKTYCDDAAIGKEAPQIDSTQPIKGEKVSLHPDKVYVLYFFNTFYRGADVCNDEFTTLSEELADKVEFVAISTDADIAKTEKYLAKEIVDENTKKQLRLAPAHIVYDQGKSVSKAYADVSNLTVMSCPMAFIVKGGKIAWRQQFLQTFTVNQSNFKAQLAHVLAGEELESNGPRPKVEVEEEEAEGMDGDMSLF*

>Lp_280007600.1

MSNRVQVYQSQLPAYNRIKTPYEAELIATVKKMTAPGKGLLAADESIGSCSKRFDPIGLSNTEEHRRQYRALMLEAEGFEQYISGVILHDETVNQKASNGQTFPEYLTARGVVPGIKTDMGLHPLLEGAPGEQMTEGLDGYMKRATAYYKKGCRFCKWRNVYKIQNGTVSEQLVRFNAETLARYAILSQMSGLVPIVEPEVMIDGKHDIDTCQRVSEHVWREVVAALQRHGVVWEGCLLKPNMVVPGAESGKTATPEQVAHYTVTTLARTMPAMLPGVTFLSGGLSEVQASEYLNAMNNSPLPRPFNLSFSYARALQSSALKAWGGKPEGVAAGRRAFMHRAKMNSKAQLGKYKRSEDDASSSSLYVKGNTY*

>Lp_000094100.1

MRYEKAALLLCATLLVLGAAAVVPASAQATSTPSGCADYHCDTNSSYDQLQSSGSGCLCCSSNSITNCVAAQKPCMVEHCFQCANNNTSICEVCDTHYQLVNNNCPKCDVTNCDQCMTNSSGPLRQCALCSNGFYWNAGLCFVNTGSSSAGASGANGVAQCRAAYCKTCQSVSAEKCETCLDSYMLDGTGKCVSSCTVTHCDMCYSGSTADCKLCSAGYTWQDSQCVTALGCRVGHCAVCFREKSSSASLVDNSTCATCDTGYTPSNGYCEVQRPCAVTNCAVCDTTLVDKCVRCEEGYGLVNGTCVKCADPYCRHCDINPSRCAKCVDRMIPDSVTGKCINSGDCSVANCKVCDATSAVLCNTCNDGYYLSLNRSACLQASTTTSTTTPTAPCNVPNCLTCYPNDGNVCQYCRSGYYTFNGQCVPIGNCYVGNCAQCMLRDGTKCSTCRNGYFLSSTYTCLSQHVNVNGAAAPHSLWLAAVAVAVTGVTCVV*

>Lp_310012100.1

ALSDCGHDHVAAAQEGLPITYVSLGRVTPSATQAEAADSPSLGATAALSQPPQPHHIVEVQAANSGGSDAATWAPIRIAVFTPDLEDDSRYCTAVGQYRPDYSGNSVECTSTADILTDAKKNTLINNLLPEAVKAHAERLMVHPLASDRTIVVNSMRGRYCGAFTVPAAHKTTGVADADFVVYVAAGPTSTPNSFLAWALTCQFYADTARHPAVGVIYFNPRKLPESSDVTADELALHGGDAFAWSDQLRRTATHELLHALGFTSTVFSARGVFATVPSLRGKRNVPVLNGTAVRAVAKSHYGMTDDDLFYGLELEDQGADGTTLSHWKRRSTKDEMMSPVINLARYSALSIAAMEDLGFYKGVYEKAEPMGYGYGAGIDFFDKPCLTDGTSNSPSVFCDMTSSSVHACTTDRLNIGRCYLTYYSSALPSYGQYFTGQPTYGGALPYMDYCPVIQPLSNAVCNSGSMLIIPGSVVSDSSRCFDASSLILKTTYARANAICAQVHCDNTTNTYQVLVNGASSWLSCGAGGDGYRVSPATTSPEVFMAGGTIACPRYDDVCYANPAAFGVLPPITTTTTS

>Lp_000513900.1

MSLVPKKSIDDANVKDKKVLIRVDFNVPVKNGEITNDFRIRSALPTIQKVLKEGGSCILMSHLGRPKGAKMSDAKPAKGVRGYEDAATLRPVAARLSELLGQKVEFAPDCLDAASYASKLKAGDVLLLENVRFYAEEGSKKEEERDAMAKVLASYGDLYVSDAFGTAHRDSATMTGIPKVLGAGYAGYLMEKEINYFAQVLNNPPRPLVAIVGGAKVSDKIQLLDNMLGRINYLVIGGAMAYTFQKAQGHAIGISMCEEDKLDLAKSLLKKAQERNVEVFLPVDHVCNKEFKAVDSPLVTKDVDIPEGYMALDIGPKTVKIYEDVIAKCKSAIWNGPMGVFEMPCYSKGTFAVAKAMGNGTQKNGLLSIIGGGDSASAAELSGEAKNMSHVSTGGGASLELLEGKALPGVTILTNKDAKASAAAGGDACPCGAGCRCGNRHGAHDSFRGVRLVVQILKLLAAVLIGVFIGRRMSHKLVK*

>Lp_000071200.1

MSTLVRLLTLLAVVAACVVVPVSAAVPSHLEEKLAGRTDIQLANQESAPYPVNKNNGDEEGSAYTDCTLENGVFTIQGAKTMYADTGRPTGLVQVLRMTVSDGSVVVTGFFPVVTLLNFSNVKGTVSANRPLIDATAAAFDKKLEIAVIDSSVAWSAAETLPSMQVLLGIPATLDGASSVFVLGVHLTSASAVVKAAIASGTLNMQVVNSSVVAVDYVNCTTCANGIIDVAPVPIFVLNHSMIRVSHITLNATPTVSIFMTNLSTVTVDSTSLLVVENITARSSNIFSSSVSNSGSNSVVLRYLHINSIGTALGSEATYTDVTAESGPSDLASATHVEGKCPAACLNGFTLTNAQLSCNCTCNSPYHRNYCTAMNDPLASYNPTGCTEGCIWCHNETACSMCSADYTLDTSTAVCKRNSGSCDANCVKCGASMCMECKDGYGVANNGVCVRCAVDHCKKCNNGYTDVCTECMSGTTLISNVCVPSCEEGYGLVNGTCVKCADPYCRHCDINPSRCEKCVDRMIPDSVTGKCINSGDCSVANCKVCDATSAVLCNTCNDGYYLSLNRSACLQASTTTSTTTTTATPTAPCNVPNCLTCYPNDGNVCQYCRSGYYTFNGQCVPIGNCYVGNCAQCMLRDGTKCSTCRNGYFLSSTYTCLSQHVNVNGAAAPHSLWLAAVAVLLASAVTHLA*

>Lp_000328600.1

MSTLVRLLTLLAVVAACVVVPVSAAVPSHLEEKLAGRTDIQLANQESAPYPVNKNNGDEEGSAYTDCTLENGVFTIQGAKTMYADTGRPTGLVQVLRMTVSDGSVVVTGFFPVVTLLNFSNVKGTVSANRPLIDATAAAFDKKLEIAVIDSSVAWSAAETLPSMQVLLGIPATLDGASSVFVLGVHLTSASAVVKAAIASGTLNMQVVNSSVVAVDYVNCTTCANGIIDVAPVPIFVLNHSMIRVSHITLNATPTVSIFMTNLSTVTVDSTSLLVVENITARSSNIFSSSVSNSGSNSVVLRYLHINSIGTALGSEATYTDVTAESGPSDLASATHVEGKCPAACLNGFTLTNAQLSCNCTCNSPYHRNYCTAMNDPLASYNPTGCTEGCIWCHNETACSMCSADYTLDTSTAVCKRNSGSCDANCVKCGASMCMECKDGYGVANNGVCVRCAVDHCKKCNNGYTDVCTECMSGTTLISNVCVPSCEEGYGLVNGTCVKCADPYCRHCDINPSRCEKCVDRMIPDSVTGKCINSGDCSVANCKVCDATSAVLCNTCNDGYYLSLNRSACLQASTTTSTTTPTAPCNVPNCLTCYPTDGNVCQYCRSGYYTFNGQCVPIRQLLRGQLRAAPAAPPAPKCSTCRNGYFLSSTYTCLSQHVNVNGAAAPHSLWLAAVAVLLASAVTHLA*

>Lp_340010800.1

MSTLKEVNGRLASQPFVSGFSPSTKDARIFEEMFGSNTNVIQWVARMASYYQAEREEILKGAHHKAEEKRARRRRPAHAPKAAEEDDDIDLFGETTEEEAAALEAKKKADAEKKKAKKEVIAKSSILFDVKAWDDTVDLEALAKKLHAIHRDGLIWGDYKLIPVAFGVKKLQQLIVIEDDKVSGDDLEEMIMGFEDEVQSMDIVAWNKI*

>Lp_000298100.1

MTEEVPQNVQFENGHPNFTYDHVYRCFKGNHNGLLFRLVNDSTKKWAFYNDTNDTVMCVKAHFSADSKVKATGRAKATMVPAEVGSQLQEMVVTVEVQPKGTEVFIDGVPNGFVLSFQTEQVPVRNVKFMNQRSTLPHYDKIYKCFKNEGNGLLFRLVDETAGKWYFYNDTKQFMMTVTVSFPLTEDIVPLGNTQKVEAKDDPDKVAYQITVAPGNCEEFIEGHPATFSQAFDARTIDESCLDPKDVHFKFSKPNSQIVDLSSSKIYKGFKDRENGLLFRIVDYQNQRWAFYNDTNVYVMKPTVRLFDSPKVSLAPGAKVVERNDDGSYAVTIDVPPLATVLLFDGLPTKYEVSLVAESVNKTAPEENPKFENGEPDKHIVNYDEVYRCLRDNPDVRLFRLVDTMRRRWGFFNDTKDTYYTATCKFDADGTITPLGSTKQQGDEFSVRVPPQSTVAFVQGNIKGFELSFPRSRPPAAAAATANAAATATATATATTAH*

>Lp_140012200.1

MPIQKVHAREVLDSRGNPTVEVEVRQEAGIFRAAVPSGASTGVHEACELRDGDKARYCGAGCLQAVKNVNEVLAPALVGKDESDQAGLDKLMCELDGTKNKSKLGANAILGCSMAVSKAAAAKAGVPLYKYIAQLAGTKEIRLPVPCFNVINGGKHAGNVLPFQEFMIAPTKATSFHEALRMGSEVYHALKSIIKKKYGQDAVNVGDEGGFAPPIKHIDEPLPILMEAIEKAGHKGKFAICMDCAASEAYDADKKMYNLTFKNPEATYVSAEKLQETYARWVAEYPLVSIEDPFNEDNFDEFAAITKALEGKAQIVGDDLTVTNVERVKMAIEKKACNSLLLKINQIGTISESIAASKLCMAHGWSVMVSHRSGETEDTYIADLSVGLGTGQIKTGAPCRGERTAKKNQLLRNEEEIGAAAKYGYPGWS*

>Lp_000265000.1

MTFEGAIGIDLGTTYSCVGVWQNERVEIIANDQGNRTTPSYVAFTDSERLIGDAAKNQVAMNPHNTVFDAKRLIGRKFSDSVVQADMKHWPFKVTTKGDDKPVIVVQFHGEEKTFTPEEISSMVLLKMKETAEAYLGKQVKKAVVTVPAYFNDSQRQATKDAGTIAGLEVLRIINEPTAAAIAYGLDKGDDGKERNVLIFDLGGGTFDVTLLTIDGGIFEVKATNGDTHLGGEDFDNRLVTFFSEEFKRKNKGKDLSSSHRALRRLRTACERAKRTLSSATQATVEIDALFDNVDFQATITRARFEELCGDLFRSTIQPVERVLQDAKMDKRSVHDVVLVGGSTRIPKVQSLVSDFFGGKELNKSINPDEAVAYGAAVQAFILTGGKSKQTEGLLLLDVTPLTLGIETAGGVMTALIKRNTTIPTKKSQIFSTYADNQPGVHIQVFEGERAMTKDCHSLGTFDLSGIPPAPRGVPQIEVTFDLDANGILNVSAEEKGTGKRNQITITNDKGRLSKDEIERMVNDASKYEQADKEQRDRVEAKNGLENYAYSMKNTISDPNVAGKLDESDKETLNKAIEAALSWLSSNQEASKEEYEHHQKELENTCNPIMTKMYQSMGGAGGAPGGMPGGMPDMGGMGGAPAPDTGASSGPKVEEVD*

>Lp_250006600.1

PSKMSQQMPPPPPQPQPQPQQQAYNPEPEALRNLMVNYIPTTVDEMQLRQLFERFGPIESVKIVCDRETRQSRGYGFVKYQSAASAQQAVNELNGFNILNKRLKVALAASGNQRPRHNYNQQQAMQGAAPQANYYPGNFAPAGYAQPNPYAQQQMMAAAMPQQYMMPQAGQQPRQ*

>Lp_000460700.1

MSSVKKALVLLVLAAFALSCVKADLVELNPENFKAIVNDPKKNVFVMFYAPWCGHCNHMKPTWQELADKYSVDKDDTVIARVDASAHRGIAKDYDVNGFPTLKLFTKSNKVGKQYSGPRELDAFEAFLKANVE*

>Lp_000482500.1

MSDEDHDFAHQGGGDNASKTYPLAAGALKKGGYICINQRPCKVIDLSVSKTGKHGHAKVSIVATDIFTGNRLEDQAPSTHNVEVPFVKTFTYSVLDIQPNADPALPAHLSLMNDDGESREDLDMPPDPALAAMIKEQFDAGKEVLVVVVSAMGIEQVLQTKNAAEK*

>Lp_350014800.1

MRSLTAFLLIVVVALTLVAAEMTEDDFRKMKVKDLRLFLSARGLECTGCQEKSDFVRMAYQYRSLNPAGSAEKRAIPAKKFWEAWADIAQAECEKAVKLRSNDPATEPFKSICSTIHSATDSYLMQHGRKVANQLKKTPQDLLQTSFKDIYFEAGSHLCQILADYCLASPAAQENCQSLGAVMSAMDGVSGADFKMWTTNVGIENTNPMYEIIDARDDL*

>Lp_040005700.1

MIAASVRRGVIWLLVAMAVMAGAVVALDGVPYEPVYHIRPPKNWINDPNGPYRDPVTGKIPLYMQYNPNGPLWGDIAWYHVTSDDYVKWTRPESPVAMYADKWYDRWGVYSGTMMNNNHSEPVSIYTCTEPENIQRQCMASPPKSDLVGKRTLNSLVKSARNPILTEDDVPGLVGLGNFRDPTEWWEDPANPGHWLIAFVARINDADGDNAHVVVFSTEDPTFQSGYTFSHSLYVYKYDLDRMFECPDFFSLAPGGEHYLKVSTMPSHRDYVIYGSYQPNATTGKYDFVEDPDRSFTFIDYGPFYASKTFHDPILNRRVMWGWTNDELSDAQIQSQGWSGVQNMLRSVEYDSTEKKIKTAPVPETKGLRLAKLLDLKNVAVTSTPTPIITSNTNNTLYHEIIARFTLSDASVFSATATYAADGSDAPEIGVVIRANADLSQNTTVSLRMPPYGPSVVSHYQQEEGWPAIKIFDGPDVANCSAECTKLRLCESYTYWTSTGSCKLYWRLTPMSESADAYSGLAREPLLYLNRNASRSIGSTAALSGRAPFATATPNGFELHIYVDDSIVEVFKDGGLETMTGRLYISNGADTTGVAVYAKNLNNVTVTADIEVYTMDTIWQAPVANAARNFTNSLYNLLDALIDI*

>Lp_270007800.1

MQHIVVFKFDAEKFAKEFPGNALQECIQTLRDANIPGLLDLNMSAKNVTAWEGYQDASRGYTHALVSRHKDAESLHIYADHPVHKALQVRLFKCIAAPPLRMELDVHPAKL*

>Lp_000083500.1

MVQSFHHVLLAALVTLVLALFTPRVYADFTEAHYTATYLFLSGFPYSFSSLQSDWSGGAFCSWRGVTCDNSSNVTYISVNLAGQSLSGTLPSILNEVTDRNLPVHSVDLSNNHGITGTFKADWAALRSIVYLDLSSTNLHGTIPDSWNSMVHLVTLNLSHTYACKGLPNWNISTLRSVDLSHSRLKGVLASSWGSMTNLTDVDISGNSFCGCVPSSWSSSTVLAKAAAAIGGNLVTPTCESSNKCKSSNYMCPSAAAEPISVVVVAGLLLSLFEPLLSCKTRKKRTLLPSPCPGVVLL*

>Lp_090011600.1

MASKCPVVTLSNSVQVPQIGIGTWEATDETVQNIKWAVTAGYRHIDTAHFYKNEASVGEGIRTCGVPRSELFVTTKLWNYDHGYDNALAAFERSRKALGVDYVDLYLIHWPGQSRSYIETWRAFEKLYAEKKVRAIGVSNFEPHHLEDLLANCTVPPMVNQVEMHPLFQQATVREFCAKHNIAVTGWRPLGKGKLLTEPRLVEVAEKHQKAVSQVIIRWFVQLGVIVIPKSSNEERIKQNFDVFDFELSPEDMKVIESLETNKRIGGDPDTFFPTARD*
