## Supplementary file 4 for "Aspartyl protease in the secretome of honey bee trypanosomatid parasite is essential for the efficient infection of host"

>Lp_000014700.1

MHSTSQSISNTTLRTRVSRLRMLLVVVYCLALLCGNSVAVARTLAAPKLTLYHLETRQVP

GADAVIRYVQHDVSVQLLADSDASQAASQAESQGGSQTASNPAPAAASSSASKVSSAAPA

GPAASSSSNVHSGEASRASSIAAAASSADHLSSVASSAETTAKPTSSADVSSSSSAESAE

AAHSL*

>Lp_000027300.1

MAAFLRAACVLLALFLVATTAQAASQSYVLERRVGSDAEWVNVGSFAISRVSPQAPARVS

NQQLGEQSMSMEQREQFAAADLIYYRAYPYRQGSGAPAHPVTVVFTPCSLIRGFDAIDSK

TVVLNENIRVVPGPNTTLLGLQMSSETNFFHSKMMNGDECDRSVVQKLFPTVRLRMKLGL

VHPVTASRKVNYEDLKVLIASEAERQGKKPKTQVRQVRNADGELVEEEVPVDDRSFLQKY

WMYLVLPIVVSVIQNLKG*

>Lp_000052600.1

ATMPAAPKLFVALAVSTVAVFTLCCTLAVGAAHHGYPSCDLSVVQSSGYIDIPGVGGTQK

HYFYWLFGPRRWPRDGSRPPVIMWMTGGPGCSSGLALAIELGPCEVNETSGELYRNAYGW

NDEAYLLFVDQPTGVGYSYGDKANYVHNESEVAEDMYNFLQGFAKRFLSPSITGGNDFYI

IGESYAGHYVPAVSYRILQGNQRGDKPKINLKGIAIGNGFTSLLIQYPFYVTYAYDFCKE

KLGTPCVNASTRDEMLSMMPQCLELIRTCNSFPTDGDPSCLAATQHCSIIENLFSRSGLN

PMDITKRNVGNLGYSMNHTQAFFADAKLRAQLGVADGAQWSTCNDEVTKLFDKDELRNFD

YVIPTILSSGVPVLIYAGDLDYSCNWIGNKAWVTALEWPGKAAFNAAPDVEFSVNGRAAG

QERTYGNFSFVRIYDAGHMVPMDQPEVSLYMVSRFLHNKRLA*

>Lp_000052700.1

ATMPAAPKLFVALAVSTVAGFTLCCTLAVGAAHPGYPSCDLSVVQSSGYIDIPGVGGTQK

HYFYWLFGPRRWPRDGSRPPVIMWMTGGPGCSSGLALAIELGPCEVNETSGELYRNAYGW

NDEAYLLFVDQPTGVGYSYGDKANYVHNESEVAEDMYNFLQGFAKRFLSPSITGGNDFYI

IGESYAGHYVPAVSYRILQGNQRGDKPKINLKGIAIGNGFTSLLIQYPFYVTYAYDFCKE

KLGTPCVNASTRDEMLSMMPQCLELIRTCNSFPTDGDPSCLAATQHCSIIENLFSRSGLN

PMDITKRNVGNLGYSMNHTQAFFADAKLRAQLGVADGAQWSTCNDEVTKLFDKDELRNFD

YVIPTILSSGVPVLIYAGDLDYSCNWIGNKAWVTALEWPGKAAFNAAPDVEFSVNGRAAG

QERTYGNFSFVRIYDAGHMVPMDQPEVSLYMMRRFLYNQKIA

>Lp_000121000.1

ATMPAAPKLFVALAVSTVAVFTLCCTLAVGAAHHGYPSCDLSVVQSSGYIDIPGVGGTQK

HYFYWLFGPRRWPRDGSRPPVIMWMTGGPGCSSGIALAIELGPCKVNETSGELYRNAYGW

NDEAYLLFVDQPTGVGFSYGDKANYVHNESEVAEDMYNFLQGFAKRFLSPSITGGNDFYI

IGESYGGHYVPAVSYRILQGNQRGDKPKINLKGIAIGNGWTDPYTQQPSFAEMAYNGCKE

KLGAPCITEAAYEEMLSLLPECLNKTRECNNAGLDYGASNAECVQAKAVYADYEGYYFAT

GLNNYDIRKPCNGPLCYPMNHTLDFYENPLVRASLGVSDAAKWSTCNAEVGELFEYDFLR

NFNYTFPPMLAAGIRVLIYAGDCDFVCNWIGNKAWVTALEWPGKAAFNAAPDVEFSVNGR

AAGQERTYGNFSFVRIYDAGHMVPMDQPEVSRDTVSRFLHNKRLA*

>Lp_000071200.1

MSTLVRLLTLLAVVAACVVVPVSAAVPSHLEEKLAGRTDIQLANQESAPYPVNKNNGDEE

GSAYTDCTLENGVFTIQGAKTMYADTGRPTGLVQVLRMTVSDGSVVVTGFFPVVTLLNFS

NVKGTVSANRPLIDATAAAFDKKLEIAVIDSSVAWSAAETLPSMQVLLGIPATLDGASSV

FVLGVHLTSASAVVKAAIASGTLNMQVVNSSVVAVDYVNCTTCANGIIDVAPVPIFVLNH

SMIRVSHITLNATPTVSIFMTNLSTVTVDSTSLLVVENITARSSNIFSSSVSNSGSNSVV

LRYLHINSIGTALGSEATYTDVTAESGPSDLASATHVEGKCPAACLNGFTLTNAQLSCNC

TCNSPYHRNYCTAMNDPLASYNPTGCTEGCIWCHNETACSMCSADYTLDTSTAVCKRNSG

SCDANCVKCGASMCMECKDGYGVANNGVCVRCAVDHCKKCNNGYTDVCTECMSGTTLISN

VCVPSCEEGYGLVNGTCVKCADPYCRHCDINPSRCEKCVDRMIPDSVTGKCINSGDCSVA

NCKVCDATSAVLCNTCNDGYYLSLNRSACLQASTTTSTTTTTATPTAPCNVPNCLTCYPN

DGNVCQYCRSGYYTFNGQCVPIGNCYVGNCAQCMLRDGTKCSTCRNGYFLSSTYTCLSQH

VNVNGAAAPHSLWLAAVAVLLASAVTHLA*

>Lp_000081100.1

MTFIHNNSIVYAAVLFVLLLCLSAPSFSSDTCVKRRFGRPNSTEAASDDDDATSRPRRLP

HSLLFARAFPQATATPLLLPPAIPAAPWQTLNESLLPYPIFYAATAVLDDSVVVLGGCLT

ASCATPVGRTEQGVTAAAVAAARHPRASLGENVSARSAATISEKNRWSRRKGHDTSEAAP

SAPAQWTDTGSPYHERSVALEVERQTMTPLPQLTLPDGLGFAGRHAAVTLTDSIYVPRSC

TMTTLSPHDIATLSEKDLTELQAFYAPIVALYPESKKKKRRAPHPDRSAAPSVNLSYFEV

PAERVRVNASCTALATENKILIMGGFLLSSQQVTASVDAFNVVTRKYESEVVFLSMPVLQ

PSVAASTGFAAVAGGWTYETAEVNTPATARASPSPVPSRPVIHYLFDLLFFEGDRWSLRR

SISRQHQLHSANAPLTSSSSICLFSVDPEQLPRDAVQRILLAEQGCHVEVFGGQVVLADH

NLGNIAVLDVRATSAPAVFAGGRELLARALFNPPSSPQPQTSHTTKMSKESGNHGGEKRA

AGVITVSVVASRLSSSESLSASSSSSSPSSDSSPAPSPSPTPPTPPPPKPVYHYRWERPM

LITLPLTRERAAAVKVAAARETSNDTTTTTTTTESPTSTSAPDVDTITDTILVFMALGGE

DVWTQEVPATEAALATTASLNRARAANRVEKLSMFSGYASSSDLHPHHEDDVVVRRAAAQ

RWAQRYVPDPTYDAARGDLLAVKMPTPIWPEGLTLQTTAEGTIHLDFPSVNYTKYCLWES

HQRDETGYGEGDNAEEEVVCAVRLSSRRDCVGNTAGTLDAPYNGAPNASAAFSASGSTMP

VYVCFSYVVQPTLWSTCGAQRSFSVLNPMMPLRILDNTPTLPPPPSPTPSRDPSDKTTSS

PLFIFAVGISVVTLMVAVLLVARLQHVPEEGLLVEDFFSRGDGRGGGAYTRVAGRDVEAN

GVEEDAAAAAVSQADTATCPRGPLTRLDAFQHLVATVAENNEARMLDAAAEVLRLHQHRY

RVLSRVGQSDHTLCFLALRKPTPTATAQPAFTLSDGGHVSAAAVGGGMNYASAGGLNSGS

LFGRRGVPRFPVSSIVPAALHTAGGRSSSNNNNGDNYYNRQPSASTSSSSSLWAARYDQR

AAVVVKYTQCPDDTTRAVITRLCERLRDLQACGGGGGGLHSFASVSNSSRDSRTSSGAAA

LGWMSDSQQQQQQHQRFPHPQTHSSYVPSSNRRTPPPSAVTSRVVSGMSAAPAQGAASTG

APTLTRNPYLNAAEERPRGSVDTHTERRESMLGSQQSEYAFSTTSAAAVAARAHAGAVRA

ASEETEEESPCAWLDAHEVDVAISLFLLLPADLFVSYEVSVLQQQQQQHQRGAEMGRAMS

GGHVRQEATLFRTTAQWNQQCVYIGQSCPQSTPSHHRHSSSSGGEVVEKGKSELRSEAAR

SLEHSIRDRLGMRKRHGSDEATREAVAKNGRLRRRLTPAARAFLTSLTSHAPRVCWDACV

NPLQPTTVSPWSLCLVMPYERAGDLADFVRRAQHVLPADAWQASAYWGCANEGKESSCGS

GVSSPLLALPRARHCWTESLLCSLLFQLGAGLQLLHQQSPPILQGDLKATNVLLREPTTF

SVSRRRVHYASTAVMNSSGSGGGSDIAGIGPTSFPGVERPSQPPPPPPLPRGTSGSYAPE

ASTDDAASGFSARPADMGSPLLPRFTATTTDRSEEDQQQSTSSTKLLPPPPEWYLSTKSY

LAVSLTDGGMSWWLTVQLPQRLRGCFGFTTQSTLRQTPAGRCWRSRYHAHTQRQQQQQQP

RQESDEQIIDEGAPPSAAVAAAPSEESIAGLAHFLFCFVEVPCHIAPELIWGQLCHLSSA

VVVASVAASHDCGAGTFHPRHSRGSPRRRGASSTKMPKAGGCRDSGTHTDQHQQLPTAPL

PPDGDPPSAPQQHAVVVVAVDPGTHVHVSEPVDKGDRSPLVNASAAPCSYSRRRGQRESA

GSGATSRDLVDDLMFDNAFADEEEANDDANSELREQAQAQEDGEEEEEQVETSESEEVES

AEKDDVVDSDEEDDDDEEGVFTTEEVSLLTSTMWKAEGLTRVTRHPPPPPQQQQQQSMHE

RGPSAATAATEGSANDNGLNTSAASLPAASRTRDAPSSLSDTSPINNGATPPPPGCGNVN

VTSPPPLVASALPFGFPPSGPPFAPLHPAHTAYRSTKCLGMSSMYNNTGEGVGGSQSSSS

VYEVLIQRVLAMDTASDVWSLGALLYGMCTDALRSEDSTADSAVVSGTCTLAQRAFAALL

GDLFAMSCGLSPAEEKSETKARRSEEACFSHDDRARWPRSCALEDEVAVAFTSAGYRSTF

SRLLSRMLSPVAARRPSAADIVEQVRLVAFPTAAASPTMTNGSPLMTTLAEPHHAVTRSD

ALNNDSRGEHTAHVSLFRDTAGGGVGGVPAVAVESGSVRTGRTAAAAVLLDERNATMELR

QGDWY*

>Lp_000082200.1

MTASKLLVAVVAAVCVVLAAASVPAYGLHVTSSAAAQFEEFKRAYGRVYATLEEERLRLR

NFEHNLETMRVHQARNPHATFGVTKFFDLSEGEFRKHYLNGASHFKAAKERAAKLPAVSA

DVSGAPATKDWRDDGAVTPVKDQGSCGSCWAFSAVGNIEGQWKLAGNPLVRLSEQQLVSC

DTVDAGCNGGLMLDAYDWLINNANGNVYTEESYPYVSGTGEEPQCNMSSGLVIGASIEGH

LSLESDEDVMAAWLAEHGPLAIAVDASAFMSYQGGIITECEGVQLNHGVLLVGYNTSGSV

PYWVIKNSWGTDWGEDGFVRVRKGTNECLITEYAVSAQVSGKTHAPVTTTTTTTTTTVGP

QPTVVEHTTCNDYSCSRDCSTTRVPVGECRKGRNGGSLSLECGEDQVVERSYKSSDCSGT

PKYYVTPANQCMSVWWGSFKDVCV*

>Lp_000082300.1

MTASKLLVAVVAAVCVVLAAASVPAYGLHVTSSAAAQFEEFKRAYGRVYATLEEERLRLR

NFEHNLETMRVHQARNPHATFGVTKFFDLSEGEFRKHYLNGASHFKAAKERAAKLPAVSA

DVSGAPATKDWRDDGAVTPVKDQGSCGSCWAFSAVGNIEGQWKLAGNPLVRLSEQQLVSC

DTVDAGCNGGLMLDAYDWLINNANGNVYTEESYPYVSGTGEEPQCNMSSGLVIGASIEGH

LSLESDEDVMAAWLAEHGPLAIGVDASAFMSYQGGIITECEGVQLNHGVLLVGYNTSGSV

PYWVIKNSWGTDWGEDGFVRVRKGTNECLITEYAVSAQVSGNSTTTTSSPSTTESPSSLV

VEHTACHDHHCHLNCTTMQVPVGKCMEGYQNTSFSLVCGVDQVVKLSYVSSDCTGLAKYE

VVDANKCVKFRYGSFKDVCVPA*

>Lp_000083100.1

MPPRTRRLIHAAILAVVVTMFAATVQHARATLTAAQQSATLAFLQKFPDEFGALKNSWTG

TDYCSWEGIGCYNDDVSIDLYDCNLTGHLPELDNSVDGSQVMVTSIGMSNSPNLVGGFPD

SWARLTNLRYLDLSSTGLSGAIPDAWNGMSSLETVKISNTYACKTLPNWNITSLRSIDLS

NNAFSGSLPSAWGSMTGLRDVDISGVYPCGCVPGTWTSSVLLHAAATLGSGVSSGDCATV

NTCHNNDDDHCLRYNATAAGMDVRMRHTLAFLRKFPEAFETLRDKWTGTDYCSWEGIRCN

EFKYVNLSSMGLTGRMPELDSDVDGWHVTVTSIDMSNNPNWSDDFEEDWGKLRHLQFLNL

SHTALHDEIPNEWSGMRALQEVYITNTGACKSLPDWTNPSLRTIDFSRNNLQGSLSTTWS

QMPALTSVDISGNNFCGCVPGTWTSSVLQNAAEAAGGSLLSSTCATSNACTIAKLRCTAA

PVSTTAPTPQPTTPTSAPTSNATLAFLQSFPVAIPGLASSWTGTDYCSWEGISCDANGYV

SIDLTGRDLTGHMPSMEDHIDGSQVMVTSIDMSNNAKITDNFRNDWARLSNLRSLDLSHT

ALRGAIPDAWNGMRSLQSIKVSHTNACKGLPNWNISTLQTVDLSNNKMGGTLSSAWAGMG

>Lp_000083200.1

MPPRTRRLIHAAILAVVVTMFAATVQHARATLTAAQQSATLAFLQRFPDELGALKNSWTG

TDYCSWEGISCDANGYVSIDLSGRGLTGDMPSMEDHIDGSQVMVTSIDMSNNAKITDNFR

NDWARLSNLRSLDLSHTALRGAIPDAWNGMRSLQSIKVSHTNACKGLPNWNISTLQTADL

SNNKMGGTLSSAWAGMGSLASVDITGNSFCGCVPSSWSSSTVLASAAAAIGGNLVSSSCS

TSNACGKNSYKCPNAAAGPLQLAVAVVATFVAAAAMMSF*

>Lp_000083400.1

MAHHLCHTIFAAVLALVVLLLTATAASVLAALTAAQNSSTLAFLQLFSSSIPDLSSSWTG

SNWCSWTYLDCTNTSNVTLIIDGAALTGSLPALTSEVTGSSVALHTIALMNMNVTGSFPE

SWGSLTALRVVNLGNTNLFGTLPRSWNAITGLTSVYAARSGACGNLPNWTHSSMLNLDLS

DNYLRGTLPISWATMAKLENLNINGNHLCGCVPESWTARVLEYAAVRSLGLRSHAPNCRS

TAKCNAAQECSRAAPDYGDAVAAPVDYAVVAPPPVVVVVTAVFAL*

>Lp_000083500.1

MVQSFHHVLLAALVTLVLALFTPRVYADFTEAHYTATYLFLSGFPYSFSSLQSDWSGGAF

CSWRGVTCDNSSNVTYISVNLAGQSLSGTLPSILNEVTDRNLPVHSVDLSNNHGITGTFK

ADWAALRSIVYLDLSSTNLHGTIPDSWNSMVHLVTLNLSHTYACKGLPNWNISTLRSVDL

SHSRLKGVLASSWGSMTNLTDVDISGNSFCGCVPSSWSSSTVLAKAAAAIGGNLVTPTCE

SSNKCKSSNYMCPSAAAEPISVVVVAGLLLSLFEPLLSCKTRKKRTLLPSPCPGVVLL*

>Lp_000088100.1

MAVRDRLVLLAVCLVSALLIAAAVAAPDGSGKVEPPCIGVDLGTTYSVAAVWQKGEVHII

TNEMGNRITPSVVAFTETERLVGDGAKNQLPQNPENTIYAIKRLIGRKFADPTVQNDKKL

LSYKIISDKAGKPLVQVTVNGAKKEFTPEEVSAMVLQKMKDISETFLGEKVKNAVVTVPA

YFNDAQRQATKDAGKIAGLNVVRIINEPTAAAIAYGLNKAGEKNILVFDLGGGTFDVSLL

TIDEGFFEVVATNGDTHLGGEDFDNSMMKFFVDGLKRKQNIDISNDQKALARLRKACEAA

KRQLSSHPEARVEVDSLVEGHDFSEKITRAKFEELNMGMFKNTLIPVQKVLEDAKLKKSD

IDEIVLVGGSTRIPKVQQLIKDFFGGKEPNKGINPDEAVAYGAAVQAAVLMGESEVGGKV

VLVDVIPLSLGIETVGGVMTKLIERNTQIPTKKSQVFSTYQDNQPGVLIQVFEGERQMTK

DNRLLGKFELSGIPPAPRGVPQIEVAFDVDENSILQVSASDKSSGKREEITITNDKGRLS

DAEIQAMVEEAAQFAEEDRKVRERVEAKNSLESIAYSLRNQINDKEKLGDKLDADDKKAI

EAAVQVALDFVDENPNADREEFEEAREQLQKVTNPIIQKVYQAAGGAAGEEPDAMDDL*

>Lp_000088700.1

MAGKAHYPDALRAVLAVVVVLLACAFSTVAADNDQVSYTAAQQTHTRRFLDAFAQAVPTL

QSNWTGANFCAWDGVTCTSGGVSVWLDGSVFEATTARLPELTSDVDGSQVMAVDITLIDF

TSLAGTLPASWSRLKHLEAIHMTNSALAGTLPEAWADMAQLRALMLSRSAINGTLPATWS

ALANLQYLQLESNKLAGTIPSSWGAWMAIRQVYLGDNAVKGPIPKAWGTSVPLAITPNAA

LVQQTSESNAAHAKIVAGVNLVAPFSSLPRCSVPNCAVCLAASPFYCKSCDAGYVLSTAS

WCHRDPSVPTTPRPPSSTTTTTTTTTSITSEPSGSDSTSGHNEASITSEPSGSDSTSGHN

EASTSTTTSTTPTPPSSTPLPPLPRCSVRHCVECVASSPFYCKVCAKGYSLTVVSFCRKG

SQ*

>Lp_000094100.1

MRYEKAALLLCATLLVLGAAAVVPASAQATSTPSGCADYHCDTNSSYDQLQSSGSGCLCC

SSNSITNCVAAQKPCMVEHCFQCANNNTSICEVCDTHYQLVNNNCPKCDVTNCDQCMTNS

SGPLRQCALCSNGFYWNAGLCFVNTGSSSAGASGANGVAQCRAAYCKTCQSVSAEKCETC

LDSYMLDGTGKCVSSCTVTHCDMCYSGSTADCKLCSAGYTWQDSQCVTALGCRVGHCAVC

FREKSSSASLVDNSTCATCDTGYTPSNGYCEVQRPCAVTNCAVCDTTLVDKCVRCEEGYG

LVNGTCVKCADPYCRHCDINPSRCAKCVDRMIPDSVTGKCINSGDCSVANCKVCDATSAV

LCNTCNDGYYLSLNRSACLQASTTTSTTTPTAPCNVPNCLTCYPNDGNVCQYCRSGYYTF

NGQCVPIGNCYVGNCAQCMLRDGTKCSTCRNGYFLSSTYTCLSQHVNVNGAAAPHSLWLA

AVAVAVTGVTCVV*

>Lp_000094200.1

MPTFTHRAVVAALLAIVLALVAHAQSIYDFPQIACSKTNLEHCSQCHYTVYNGETFYLCA

TCEDTETTRYSQASLGEHSGTCQVYNGYCNVADCDKCASDDPSVCMTCNFGVPDEKDSQC

YGRTISSATGGAYTSSKTPVKAPTIAESTTTTAPKPTTTTTATLKPTTTTTTTTTTTPKG

ATTTTATLKPTTTTTATTTNTTPKPTTTTTATLKPPPTTPPPPPTTAPKPPTTT

>Lp_000099200.1

MLEFLSTMLVSARALSRRHYTLWFLAALFVAWCARAGTFAAEAPSPSDFEFTCEGPVHLY

TAEQTSATRDFLMAFTETLPALSPL*

>Lp_000122700.1

MVVATLLVSTTIILAAAAAAALPATSSSPITETELIAVDLGHENMKIAAWRVQEEDLYIK

AGGTGTVTTTTTAMSGSISMVLNDQTNRKSPPCVAFRYFKAPAGSAVNEGDLQGANAGFH

SDGKNGTGGTLLYPPGYQLERTFAEQAQALAPRFPTQVICSPAQLLGITLWNSTSPTDAT

AVAAATTPNTIDAVSEVVTPHQLTAAYSFYVKPLNDLLQGRSGGNPAAEDSGSRPPQQPR

NHTRQALGVFVPFFPTSTTDAVDEGAFFSTEELTAMLLGYARQMAEKADAVDNALSDEDE

RQLMDLLNRSGVATTSGRADLLRGHAVPQYAALTIPVHATVAQRQALIDAASLAGLRVVR

LVHSTTGAAVQLAYMKAEQVFLPDKPQYVMVYDMGSQQAEVAVYSFTAVPASVARKVKVQ

GSIELKALVGSRTLGGAAFDECIAAQWDELYFNHSLLAGVGRASTEVARRAAAKQRGSLL

RAAHKAKETLSVNQEAHITLDGVHADPELFRRAAQTRQQGTVSVSPDGLLSLRYTRAEFE

RTCASLFDAAVALRDDAIAATGGLVASVAALDRFEVVGGGTRIPRLLQRLSEDYRGEQNL

VDRTLNSDEAAVLGTTLLSVSTAPQGLQLRGRTAALPQFRVREWLTNAVYVSVTPTFSSD

ADNADDGGDDAQVKREKSAEVGVSTASAAMAKESEEAAAYLRLLFPAHQVVVPATRSVRV

RVSSRKGEAHSIPDHRNDGEAPAAVVQHDNITVTLYSGARADKAYHTRATTTARAAAAAS

PHVNVSADSADLVAAVPAAAAACASCYVRAYVVEDIKAAEETLRGQVLRQHPVGSRVLLD

SAEVVAEVVATVSGIPHCNMAYLRAVYHVTRASATTTTTMIATPTAAAEAEADKAAKSPQ

RAGADELNERAEAPANATEAEEAEENPNGDKTSAETVDVADVAVVDNAAAANLPTTSALR

QVKMMALPLRSTASARPIPTTDKGSGSTVMPTSLQSYNMGRAELRASHERLRRLQAMDAA

RLRRSTLRNDIQSVIVWVKDQPAWGAPENNNDNQDNEKRNKREAAADKQQDRVKKTPDNA

VHELDGAAWRGTVREIGEWLDDYGETASVAALEERLAAIRAVKMALRAAQNQ*

>Lp_000142000.1

MRARPFSRTPVKSVLLACLYVAIASACIAVFTAAAVAAEAQSSSSFRCLRPFGVVDSFAT

VAKTDVSQNCNFHYFTVTDASAELSCITTMSDDALSSISAGAAPVKGGRNKDKKKHDVLD

EGATSCNVTVRWSMRRADDVVQLLRRQVGTDADYFARNASETSTLSAAAAPPGEKERQQR

MPAKSFLTEDDDVWRYALSLRAKGDSTVQYKGFLNGDGYSLCCDGITESECAWMAAEQGD

NDMAADEEVNPSFALQQRRMLRGCPLPFPSAMPSEEESDYDAFGTAPGARVLLTDLDEKD

TEEGSEEGVFHGKVLKPLHRLVEGPWDVTVQMWRRRQRLPRASSTGAAASAVPADESTEA

EVLGRVVVPFRLNLAELQKEGRVQHVESMALTVEEVAEDDREDL*

>Lp_000149800.1

MPRFSFSPSLVAALALLLLLLGSVVSVEASYWSDEVTRVRTYAAQYYMNRIAKSPNVSAL

PSGLLVTVLSRGSGDRAPSADDVCEIHYTIYHRFPGVVDDTRGQPYPVRRSPSQLIPGMA

EAMQLMREGDRWFLYVPSKLAYGTEGWKERRVAGLANVRVDLAVFKCDNPRGKTSEEIDA

YLAPYLKTPMPAKSAPIDYADL*

>Lp_000161700.1

MPRTETRTAHVRSRFSRCLFTGAVAVLLLLSLHGSSIAGTVRQSVTDHTNTAELTRRDMP

PSLSLRSSLGTFAVPKGRTPIVSLQAACRSEVKLYCSGLTSSPLRCLVERFYRERQLGST

SVFSDVCEAWLTARDACLSFVLTHGAELCGAAAATADARECLRQIPAAFLPPKCVSSDYY

EGVRLVGKLRQHQIADARLRRIREQ*

>Lp_000168200.1

MTSTQSIYRTPQRWQQQRQRRTVLVLLLVFGLFLFAAPRLSAKDIRVRVEALWNETSFLQ

EGCEWAGRTFGADAFYDCAEALWTRAPSFAVASDRGAAAGGEEKEEGGARFLTQMQQAVM

IDTVARTLAAEHLTDHSFFTLEMDARVYSPAVEAHYATAVQTWAEVTKETTATAIPDEAV

APSCACGEPFGVLYTRALSPDSQLHAQLVCSGAALAVARAALAALQQSNSTAEADGDGEA

VAQLVSVREVSLPRLDYRYPTPVVAAASDVAAVFVLYGLLGSTATQRLHEAAVEWATQSR

GATSATAAQAASYPTLVYFFRHLPVSRTRLCGSSPPSPHPAATTETEAEERHHHLHDREV

FLREHWDTPLAVVGYGVTADIKSMEYKVVDEKAAAQQQAKATTTTEGEASLHSTAEANSV

SPGTESRVVGGFHVQRLKERYPHLAASLDQLATVVSTAMDSANLKVKFEVWELQNIGLAA

TQYIREMEDSTLRLEVLKDLVTQFPLYAAALSRIAAQPERLETVQKAQTSGRPRLPPGMS

ALFIDGWRVEERDLSLFGVLDALRADEVVSNHVKRALTTRLVPHGDDAAAASRAVSTVNV

EAAVLNKVSDYLKRAARRGAADGSEAEAASVAAFAIPSQHVLWLNNVETEKRFDGMPRKL

TSFFTQVPDITPFPRRNLLNYVFVWNPLRRSHLQMITLIYRFNHQGMMARYGLLLLDPTW

SSEIVSEAMAGDGAAPSSLSGDVLAISAVVYHLIAAGRPEAVLEMLVQLLQAAAAAGSST

SDVIASETVQQVCRHVATVMFEGGSLDELVSSVDFVNYYHDTQAMLRRFPLPEYPATLLN

GVVLDTFSGGFTAGLEYEVMRLREWVASGALQDGMEDLYTQILQLRGAADHLQPALLRPP

RTMLWTETQPVVAFIEDAPYLYSATYTFDVPSLTQILVLPCHSTTATLQQLQTVLAALAA

CAEMVAGEGKKGPRDAVCRSLRVSVATCPTAATASTAAHNMGNESPSLSARSFSLSHHIT

ALLRQTVQRSGVAETKRFSILQRYVEHLLKALSTLPIGSAAHRRCSGWLDGETVAAALAH

NPLPNDLLARDHSAAAADDDDDVWQRRNAAFWAALESATPIGDVNVTLVTNGRVVTMDST

FSTSDVLEAARQVLPLTKKVEEAVLKVNFAEMTPTERQARFTSEELDNVFYSAKVACLTS

VFAREAVAHEAGKSLFPLVREEEIFATAEERKRMRPLLFTVNTTATVGSSGGSNAFPSSA

ASSVDGEGGVAAAAAAREEGRVLHTIFAVVDPSSRDAQLIVSLAHRLLFSPLRIRLTVLL

NPSPDVKFPIRNFYQFVGGGSLAFDTSRRVAAPEARFDDLPPTALLTLGVEEPPTWTVFS

QEAEVDLDNVLLSKLPARTRSVAAVYRMHSILVTGDATDTLTGGPPDGLPLSLTPAQRCY

HNYHTDAIHTRSTDTQVMANQNGYYQLQAAPGLWYLSVKPGPVAAAYCVEAVDDDTVPSC

AEGDRDVNFTQGQRIPVMVDSFSGRYLSLHVRHTPPPLSPTSKGHHSSGRGGSSNVEEEE

ADSRDLHSILQQMAADVKHTWPPPWRSRGAKPQPPAKPTLNIFSVASGHLYERFLRMMFY

SVHRTSSDKYGANTTRIKFWVIENFLSPQFKRYIPLLAAELGFEVGFVTYRWPWWLPRQT

EKQRKIWAYKILFLDVLFPLDVDRIIFVDADQTAQADLHELYNMDIGHSPVAMTPFCQAN

RNEATVKFRFWEHGFWVDHLKGKPYHISAIFLVDLRRFRAMLAGDRYRSTYANLASDPNS

LANLDQDLPNYLQASIPIFSLPEQWLWCETWCSAKSKPKAKTIDLCNNPLTKMPKLDNAK

MVIPGWEELDNTLQNMSDALLELH*

>Lp_000176000.1

MSALQLLTVALAACLSVASADKSEFPLVWVCLGITVAGVIVALFVAYKRPAELHLPGSVV

MTAVESEPAGGKLAEEPI*

>Lp_000233400.1

MLRCGIVVLLLFSTCVALFASALNVTQNEQSTFVEGIKILPRSVQLSNRSFCFFTIPVQG

DSQRGDVSTLCSFDGGASVKPAKLSHSICTSGVVVDQSLYCPNEELPHDEEGEMEFSSVA

HAVQSGELSTEDSEKRFVFKSGKQTGEVKQILFDGNSVRLDDDDAHILVSTIINDAEERR

VAVFKSEDGFTFRAIAVIPDVEMAEAHYLFSEGNRRLSVVSAYENSIYTTTSSGYAGNFW

SAPKAVNVTTPPASAAFSSGVLIQYACSNETSKLAKWYLLEDVAKHPLSPKSFSVPALKD

AGGTPLLFFSVASADNTNELVIVHDDPGSNPAAGIRLSVYKVDDSTEAKEKADRIAKEME

ELLRREAARFKAKMERLEREKAQRREQRRKELERKRKFLADDAPNVNAAKSYMDMDGEMI

IIRRVHKDSIALEKEVFFSDL*

>Lp_000237800.1

MLNALFRRLLTSFVAAVFVCSLLAVSSTAAVRELVEIHRVVTIGDVHGDAENFLEILRIA

DLVESPQGGAFDVLADPPKWKLSQAPNGSVSVHTTLVQLGDLVDRGEQDFESLNIAIALQ

EQTQQSSLADKVILLIGNHELLNIQGYYHYVNKRNYGGFMSKPLRVEAMNADGAFGRYIV

ENFKTAHVEEDTLFVHAGIELDVAVSSVDSLNSQVQEALRSRNFRHAYLRSNGPLWTRKM

ISDSMLGECQEVEAILQKFNVARVVVGHTPQRTGHIEQYCDGRVIAADVGMSRWMYNNVA

ALELVFLKYFDTERQQTTTNFIIRELRDGSKTFSPAQIKRSTLKDPGVTTSEAIVNEGEN

DGDL*

>Lp_000239600.1

MQPRCVAPRRGPPFSVCLLTLLVPLSFLLLVRGSAEASHDGAASGIGSAFELPSASAVLN

LLAESSAARPTLLLFTDDSEQDKATMTDPIQAKVRQLATDIPPSVLRVHAFSVPVAVTER

KLVDLLLNGVATQLPALVLFHSRVSKAQVVPGSLISAVPLSPPLPYPHADQLHRASYMEL

RTWVLSELPARYADPVTFHLIPSLQFIFNPAETQATLRLVQQAAAAAAGEGAAASAAQSG

VAGGRAALPAVVSMAYVRLTTHGSEDVVAALSSLATQAGNAVLTLVTESREVAAAWGLTQ

EHTMATAPWAAVEAAYLTNTNGDDRATTAAVDEVVLIDGVSPAQAIGTVEEVAEATAGSP

DWQHAVESTRAVEASELRAWSRAMEAFNTTSPLRKIDSAAHFVHELVSLQQAIKIIFVLR

ESDEMWFHHHLDIAVQLASRMRQTSVLYNTTTLNSDGGRKGGSAKMPSRVLRSWSPPMRV

EVFWTDAEQLPEVADGLLVAQVPSVLILVPLQSRFQQMSEGEGEAEARASEAGLRSPDPF

LGVHTVNRYDLFTAAYVNDAGIAVDPPTGKEAQPLFPASSDALIRFLASDSFLGALQSTI

RPVRLSQLRTSLSADAARTRGRAASPSSASSLPNRRYAQLDHRYYPLRLAEELMEGPAYV

RQILNGSSPLPVLTEAERRAAEGATRRKSDAPASAKDTKAAAAAAKKKASWEAELTKRRR

EREARIQRKAAEDAATREKEKVVFQKSIEAAFREEEKAAAAGSEAGLRGGGGGATQVEGG

LLVRRPSSEAALEKLAAPEMSADGSVDVTTEREDAEDAAQRVRRRRRYKEYKEWQADRTR

MVDNSVRWSGERALSLRFKWD*

>Lp_000240800.1

MLRFRYLCVLLCFTVLLSSALVATASLLTEVPPEQVFLLHRRHIGTTNENELRAVTRGLP

IPTSPNVSLEHLHGLSSRQLSKMLKDRDRDCYGCVERRDLVQRAYEVQQLPTADERVAWQ

LTVSDRGLMTTAARLDNIASITGSFLNADCHFFNGTIYCQPRGF*

>Lp_000266000.1

MALSRLMRVARLLVCAATVLCVAGAHAAVTVTTDPTSPLLDVAFCATLSPSSVGSRALFV

AAASDCSTPLANATRECGTSGWYAFNVTSEKLKIVLFRPVLAVPDLYWCAKNADGSTTQV

GSLHMSVVRTSPMYFYKGESNTLTFNEATPVGSTVGVFGLSSCEFPLSEELSVGSPGNSV

TATLNTKFPYVYICAEVPSTASSGFVPMTLRNTIISAVRYSITPVVGVRHDTVTVDTQVD

SIVYYAFSSSSTCSDLPVPDVSNGISAGSAKVTVNKLGGEYYFCGADTSRVYVPAENKFT

VREYGVQPHTTYAGMPTAVSLTMAAAEVPAKEAALFTTPGCEGTPVQRWSSSGQLLWTVA

SAGTYYACVRELRPSSSP

>Lp_000274400.1

MRRSYLVGQALTLLAAAFVLVGSVTLAAALTSSDRNGLHQRKCVRGDYVATYGPCLANST

RLRSWKKSASASCVVGDAAGQPASTFVPCAACPPGTQHQGGKRDTISDCVSCSVGKYLDV

SELPSTCTACPFGTTARPVLSFRDGFDGFSGNASALSRSLVTYMASPLSNPSAWQVLNGT

GMDLVGYSRNGSTDLLVGMRSFQLSNDTFAVSSSFKYGFDALSDGLVELTFSLHHIDQAG

DERDEGPGRLEDLMRHRFVLFVDGTEYDVRDAARSLADSRDAVFVVVVPYMHTTTRREHY

IGWQTTDYSQSTSTHYVLVRSLVVSGDLSGGVDACEPCPAGYACAPQTTQASPCPPGTFQ

PATHAQRCLPCTGNTIAPGYAFRACLACDYNRSANADHTMCDETCIYAYDDVLYDFTALS

GVALMTAVDETAVSRAAKLTIEEADEADIDRVYLSLCHAMPIRDVATEEMSPNGLGRTAA

GNGTFDTLCVGDVHGHNSSTSAYACQRVNSTTGRHFGNVVDFASAADGRVNMVTSMGSLL

VPLDMLGNASDYAERTWRAVIQLECSHDDDDGGSNSDDHSHHQHGSRDAHSLRVVGLTTD

TMTLAWRSKQACPLCTPQSYTKVESRCNASSFYTVTYERRGTVFCLEGYTPPVPHVAPCT

PCPAEAYTLEWQACDVATQTQTGLFAVKPAYQGCRDSEAWSPSNQTRSCLANSRTTSTAK

KVSLVIGVLMILLACGLGFMLVNPSDSPQGLWSAVAEDDRELDTYIHNDHHDDNPASLST

HYSVADSVENGTAPSDTAASRFSDLMDRLAHSMSAAFQGVWAAGSTVTASSTGRGSPPLS

GMQQGNGYRTLATTDENDLLFSDDETAQRPSHTSSQRHQLLFTLEDEDDDDLLPPHFSS*

>Lp_000291900.1

MRAVVVLAFLAAVLFTVSSFVSAEPEVTDKVFFDITIGDEPAGRIVMGLFGKEVPKTAEN

FKQLCTGEKGFGYKNSIFHRVIQNFMIQGGDFTNFDGTGGKSIYGSKFEDENFKVKHFVG

ALSMANAGANTNGSQFFITTAATPWLDGRHVVFGKVLDGMDIVHRIERSKTNSRDRPLKT

VKIVDSGVVA*

>Lp_000298500.1

MPAFTAKLALLLVAFALLASYASADCAISGCINCALSNPNMCLVCDDGFHLTSVGSCLEE

GNGAHGPQSMAVAALVLLLGMLMYVL*

>Lp_000317700.1

MFSLLLELFISLVLCGVFAFFAFLNAPVDAGATTTDDDGSAGASAADGACEPALLAPHEL

PLYSVLSTYVTLGEDTPCGIAYRWLLINALHYTPREVYSEAQLREWGSSVGNFGPADDET

GGSTSSSNKKSVAASNSEESSGMEVNGLQRYGRASPSPSTASPSPSAAAAAGGCGSGNPT

PRRSRRRHRRGGPLTATAPRTPTRHESAHWVNVVLRWAAFLCLGGGTVKPEVWTDHLLYH

VEGMLGAVNAGYAEKMRARAAVVRLVASQLSNAGGQPSPSPQQQQQPLHSSLQEIGSSGG

GGNVYACFPPSTLPLPPFYAPRALVRVETLELGAGLLGGPAREVRRPQLSTTADGLLSGG

GKRVDGAGALPGNISPSLASTAPANPPAAVVNPLNSPTLGAQSGPATSPTVAAAAATAAP

LHSNSNGGSGTAASNSPALSNTHPSKTTTTTTITATNTTSPHAPPPHNTAAAIAAASVLL

PNSIAGLTSALANVVSSVTTAANAGGGGGGVGEGGVAGDYGASNMSAGAQLGVVLPRVEG

DVISVEQPYANGETHTVATSAASPPALPLRCFAVPLLYEDQRFHLRLGCCLPLGALLPVS

LCVPPDVLTLDCAVAVRRVIFNGHLYAALHGAQVELSFPVAPQFTAVVEVMPDNASGGSG

STSASSNYHSYPDANHLRYGMGAGVARTLGGHAFTAVNTTRNEVSAAREGGVGVGNAREA

AGPSSLPPRPPRPPSSSLPFTPRSGVYSTYANNFYKGLHSAAAMSSAGSTITTTGTSSIN

ERNEKVQEVVLLAVRRLIQSLTYPNVLAGQLVCEPAVVAEGGDVNGSLGGGAVARTLAMR

WQRTTAKLPLRL*

>Lp_000320700.1

MKQSFLLLAVCVLFLCVVSAEVQVATKSNFDKIVSGDLTLVKFYAPWCGHCKALAPEFEK

ASTTLKGVATLAEVDCTKETELASRFDIKGYPTLLIFRSGEKTEDYEGPRTAAGIVAYMK

AQVGPAVTMVGNAEQLEELKKEDLPLCLVKTASADSALAMTMTKVANSLRTQLNFALVTD

AAISPDDAMESVTVYRHGVEREAYAGASPVTVEAAKQFLSEAQLDFFGELGQDSFQTYME

ANKAKPLGWFFVDKDTSPELKKEVAAVAKKYRHKVLMSWIDGDKYRQVSTQLGMDKDVKF

PAFVIDFERRHHVMPSEAPLTASSVSEFMEKYVQGNTEETLMSESVPEVETVEGLTTIVG

KTMAKYADGSKNVFVLFYAPWCGHCKKLHPDFEKMAKELEAKDVIIGKIDATANDFDRTK

FIVSGFPTMYFIPAGGKPVSYEGGRSAAEMKAYVLSHMVDTPAGASTSATAAAPPAEDKK

EDREENNDDL*

>Lp_000328600.1

MSTLVRLLTLLAVVAACVVVPVSAAVPSHLEEKLAGRTDIQLANQESAPYPVNKNNGDEE

GSAYTDCTLENGVFTIQGAKTMYADTGRPTGLVQVLRMTVSDGSVVVTGFFPVVTLLNFS

NVKGTVSANRPLIDATAAAFDKKLEIAVIDSSVAWSAAETLPSMQVLLGIPATLDGASSV

FVLGVHLTSASAVVKAAIASGTLNMQVVNSSVVAVDYVNCTTCANGIIDVAPVPIFVLNH

SMIRVSHITLNATPTVSIFMTNLSTVTVDSTSLLVVENITARSSNIFSSSVSNSGSNSVV

LRYLHINSIGTALGSEATYTDVTAESGPSDLASATHVEGKCPAACLNGFTLTNAQLSCNC

TCNSPYHRNYCTAMNDPLASYNPTGCTEGCIWCHNETACSMCSADYTLDTSTAVCKRNSG

SCDANCVKCGASMCMECKDGYGVANNGVCVRCAVDHCKKCNNGYTDVCTECMSGTTLISN

VCVPSCEEGYGLVNGTCVKCADPYCRHCDINPSRCEKCVDRMIPDSVTGKCINSGDCSVA

NCKVCDATSAVLCNTCNDGYYLSLNRSACLQASTTTSTTTPTAPCNVPNCLTCYPTDGNV

CQYCRSGYYTFNGQCVPIRQLLRGQLRAAPAAPPAPKCSTCRNGYFLSSTYTCLSQHVNV

NGAAAPHSLWLAAVAVLLASAVTHLA*

>Lp_000358800.1

MPSLARSAAALIAVLAALSGFLTFTEARLRLPYTTPVIQTTHSAVTHPETGHEVAVLPVD

FAVTVADCFSKPIALVNGTRVRGAVIIQTHIFKEASMKLVFDVPNPGLDFVMNLTSDEFP

VIQDQYGSLWKPVSPLKLEMVKQDTKYTYDVGGVGMTSYQSQSESVITPVGTPQYDLECY

RTQACCGLNVTHFRKYALIDSPVPPYACTRSPVTYQPSVYSFTLRGRVYQGTAAAQVLCR

LKASDGAETSFDCTGGAAFHLDSIVTSPSYSIPSTGYALCRQGTTLIDPAYEALPIPVVY

AAFCSTSTCIGYTSKNYMDDRVKCLEHSLGFFTSLDGNNSNQFSFYENKASGLWKVTFTG

LRWIDDIGMSSPTTIQNISFQRDVPASAGWSLTYADFPLPLTDALQPLFPIGRPTRVTAR

LNDDASPTAVVVSVEYVSFSNFTATNWTIRTCVGVLCSSSRYPAAPDMNRYPTLMSVTVP

LSSLGLTSQLLLPTEVNVTVICETTPTANPFRTLLQLSTVTVQAAIRPPATQTVNASLAR

EPLQKVPSPYAASLGANATSRIICQGDYRFSAKTGTCVPLTDRQCAIKYRGRSILFNATM

NKCMCRAPRLTSPLVYRPLPDPLPPPKFTEENIRALVKKIKLAKFMQTMESAIAEHDTKL

ASQAKSGAQESLTLLAAASPPQAPSSVSAAAAANATAAGEAAPSLPAEYPLLYKLCIASF

VVTGGSWVFILFRDVFYKWGGVGRWAKKDKEAEQQPVAAPPLTPTSVPPAAQTPPTQPLT

ASPLPQPAQPTPTPSTQRAGQQPKPSKPVKDKRARGKGVAAKESDQQQQQSAQAAGAPVN

SNKTASDVHPQPFTASSHQPKHHHAGKRSRGESRRHSSHDRQPPHASRQTAPTWSDVPPA

ACEPPPLHFDAHGFAAEQHTPFGYTSRGYAPDADTSLRGEFTSRSGRGGVGEYGSGRDFA

YYDAYEFMPASPTFAESPEDPRWFSFAATPPRSPPTEGAAWGSCGFGEGYFADLPPYARV

PSMHDPFGSSHSAAAHSMPSGRPSFFDSEQHLPREIVRQSSRVEELSSDADEHKTSHHTP

AETRSHHVESID*

>Lp_000382700.1

RLATIFVITFAVTFLIAVPVLEARRSSGATSPSHEVEGVEDVNLENYYDVVGHDRFVLLE

FYADWCGHCRDFASVYSDFAQYVQVRPDLVDKLVVAKINSPENKRLEKRYKIEGYPTVLL

IPPHSHTGIEFQHSRDMNALIEFVERHVIKK

>Lp_000390800.1

MAGLPVATTKQRGGVALMLLAAVVVIVACAAPATAQTATTVPPIIQCPATFEHCKECHTV

EGMQICSKCDDTYAPNTMGECAPYNGECEVPNCKICFSDSKTQCTDCNSAFQVTDKFTCE

AKSTSSTAGPTTAAPTTAGPTTAAPTTAGPTTAGPTTAAPTTAAPTTPAPTTPAPTTPAP

TTPAPTTAAPTTARPTDPPCAVAACALCSIGMPNVCVACTQGYSLMRGGACKPTGSCSVA

RCAQCFVDDDTVCQTCSKGYSLTVSGGCRRGGHSAAVFHGPTVLVTAIVAAVTYALTSL*

>Lp_000403100.1

MLAKCFYLLAVLTTAVSFVFSDVTFYPQGTHVQFGDSNLKIRNLVPLKLDWPSTNISSTA

EALSFSLDNPFSFDFTVQVFTFFFWMPSKMQVTLTGGAAYTKNGNGACDSQSLVREYCDI

KARVVIRGIETNLKIVDVAIDICDVVESMFAPYGQQSFPAVTKGATDISRIIYFQKFAVL

GGLSEVLHLPCSSKFVAKNVMEVSTGLVLPLFFDFDTSNNITSGAIDTVTTITNGIAKLL

NVSIYNNLTDATTTIAYKGGNVTMTSLKNVLDQIMTKKARAALKVYVPAGASLVYDVVVN

DLQCRFFNVICSIPASNGIQVMNSRFTGLGDFNSILGNALGKSIDEMMTNITFGAVKKTS

ILTGGRYYLPLI*

>Lp_000403200.1

MLAKCFYLLAVLTTAVSFVFSDVTFYPQGTHVQFGDSNLKIRNLVPLKLDWPSTNISSTA

EALSFSLDNPFSFDFTVQVFTFFFWMPSKMQVTLTGGAAYTKNGNGACDSQSLVREYCDI

KARVVIRGIETNLKIVDVAIDICDVVESMFAPYGQQSFPAVTKGATDISRIIYFQKFAVL

GGLSEVLHLPCSSKFVAKNVMEVSTGLVLPLFFDFDTSNNITSGAIDTVTTITNGIAKLL

NVSIYNNLTDATTTIAYKGGNVTMTSLKNVLDQIMTKKARAALKVYVPAGASLVYDVVVN

DLQCRFFNVICSIPASNGIQVMNSRFTGLGDFNSILGNALGKSIDEMMTNITFGAVKKTS

ILTGGRYYLPLI*

>Lp_000425600.1

MPAGFATRYAASRLRSGIRSIAVVLVVSLLLTTGCATATSKKKDPSYLRSTVASSGGAYT

LYTVTSCGECTVNLSQRWCPSTMHCYPASNCTCDGPVPCMDLRTCFYGTRPSCRECVDSG

GVYCAGGASAQTLRSGHSAPDARCYPPEGRPSPVATAADVADGSKGSAEVSTRGAGLLAL

LPTCGVSTCQGGRCVRFAGDCPAEVSNSLMRSYEAISAVVVLLLSALAIHSIFRLVL*

>Lp_000430300.1

MRTAAILTHSNHVHRTGWLALFVCALFFLIRGVLAADPAGRTPRTVALTCCERSEEAWTI

LNAWQRSCANAATRDALTTKKFAAMLSLQSRSPIPATVASAVCEDAALSSGAVAAYMRHA

LCTSLPRNHADLARSVYSALMDEVPELEDELTGDMEAACRDLQSRWIAEVRAWTRQLRTE

TVLSSAQTALCPSACRWKNDDFGGPTYDL*

>Lp_000431400.1

MLAQCFYLIAFITTFITVTVSNVTLFPDGVKLMFGSHSVVLLNTKPSSVKVGMSRINSTS

TGFSVLFVDPLEATMVSNAVVNGTLFLSNVTIHASVSGSLEMTKEVYDTCRTDTIAMTYC

DMKMSVEARGPLDGSLTTGNLDVCDRAENIVTELLTPKPYMLYPAAQPGTTSLTKSTYML

KLRLVNLFADTTGMPVEAEFVTSNVLRLSIGAPVGRGLTFDSEHESDVEVFDAIAVLAKL

AKKMLNVTVDPVKGNHTVIPYGESGEIRVPTLRSIIHQAIAEGTVVALQTYVPRASSLVY

DVIVNDFRCKVFNTICSLPADNGIEVVNSRFVGLGDLGTILDNTAGARVDALLKNVTTAA

FKVAGSTFGNRIYLPVL*

>Lp_000455600.1

MSTFAKYFLACVAVASVFLASAAVALSVTERVPDGTYCGNYGGGLVVGNVTTQAGSDKFD

MIMVGLGLDMTCENETFIYDPTTHHAKVPGATDPHDCLGSVLTDGGLTLDVLYKPDVDQL

ILDLGFTKINCKKCSKYGLAVRF*

>Lp_000484800.1

MSRIFLRSGLLSFALVLSSLLLLLSPAAAFPYGKSSAVTELTPASFSSFVNTHKPVVVLF

YAPWCGHCKRFHPEFERFAESVKGTMRVGAINADQYSNIGQQYGVRGFPTIKYWKMGTKS

IASPQDYQGERTAAALQTFMVADITSAHVKAVATTEQLKQVTREAPQKRVGVVFSTKNKV

PPMLSVMALSQRLSKFPLVFAGGCVLDKGIAQSFNVVQLPTIGVLKYTPAEGAGGEDVFE

LVPYPKTVVAYEPVAHFFLDCVENDCQSPQGSKVAASATAPTEARDAAAAPADLDEEDER

HKRDGRDGHHHHNDRIRHEPQDHLRAHRRRQPPVALPVEAVAFTNETIANFCSPDSLKIR

GRSPLCVISLTGHVNLTSLQRQFQNDPLLFFDAAAHREDVLRLFRDRFGVVLATDAAVDA

GAVVLLRQGRPARMRHRLLEGMNSDSDLQRALQKMLNGELRLNKTYLDHEKAKEEE*

>Lp_000521000.1

PRRWQPAALIVAALLVAVFLATTAACVDAAESDETAAADPYAYLDMLAETDLKKMLFEKT

GGRVNLDTYRGKADLVAAVRQLEEHEEREAQFNAQVEAALARKAAAAKKQQQHAANSNDD

SNSAAAHVAASNGNNEHHNNKRAVQLIDDDDDDIVRKEGMPAQHRAPSKAGRVVAKHKLE

VLYCTG*GYAKYFEEMKEKLQLSLPNAQEVRIVGATYPTPPLRAAVAKGCSIAFLGSLAL

ALAGPQMAFLPVAVLNFLVQQRGMVIGTGFMLNMIGNSLSQTGAYEVSLDGTLIFSKLQS

GSVPSIEDVRRVILEKTLLEEYGEAA

>Lp_000524200.1

MPTFTHRAVVAALLAIALALVAHAQSIYDFPQIACSKTNLEHCSQCHYTVYNGETFYLCA

TCEDTETTRYSQASLGEHSGTCQVYNGYCNVADCDKCASDDPSVCMTCNFGVPDEKDSQC

YGRTISSATGGAYTSSKTPVKAPTIANTTTTAPPKPTTTTTATLKPTTTTTTTTTAPKPT

TTTTATLKPTTTTTATTTNTTPKPTTTTTATLKPTAATNNTTPTTNIPCGIDKCTTCATS

GTTCSVCYMGYTVTHTGACMPSGLCEVTNCVQCSATNAARCTTCAAGYTANTNGKCKRND

ANSAVSSPTAALTMTAAMMVALAAVF*

>Lp_000532500.1

MTCYTKSTVCLVALLSVLLATTVSGLYATPKKSPLLQEDFIAEVNKKANGQWTASADNGH

LITGRSLDEIKQLMGVRDIRNHALEPRVFMADELARDIPESFDSAENWPQCKTISEIRDQ

SSCGSCWAIAAAEAMSDRYCTIGKVTDRRISTSNLMSCCFVCGMGCNGGFPSAAWTWWVW

VGLTTETCQPYPFAPCAHHTNSSKYPACPSTIYDTPTCNSTCDSSQNEFVKYKGAKSYSV

SSEEGYQRELMAGGPFEVALDVYADFTAYKSGVYSHVSGERLGGHAARPVGWGVLNGVKY

WKIANSWNSDWGDQGYFLIKRGSNECGIEDSGVSGTPATN*

>Lp_040005700.1

MIAASVRRGVIWLLVAMAVMAGAVVALDGVPYEPVYHIRPPKNWINDPNGPYRDPVTGKI

PLYMQYNPNGPLWGDIAWYHVTSDDYVKWTRPESPVAMYADKWYDRWGVYSGTMMNNNHS

EPVSIYTCTEPENIQRQCMASPPKSDLVGKRTLNSLVKSARNPILTEDDVPGLVGLGNFR

DPTEWWEDPANPGHWLIAFVARINDADGDNAHVVVFSTEDPTFQSGYTFSHSLYVYKYDL

DRMFECPDFFSLAPGGEHYLKVSTMPSHRDYVIYGSYQPNATTGKYDFVEDPDRSFTFID

YGPFYASKTFHDPILNRRVMWGWTNDELSDAQIQSQGWSGVQNMLRSVEYDSTEKKIKTA

PVPETKGLRLAKLLDLKNVAVTSTPTPIITSNTNNTLYHEIIARFTLSDASVFSATATYA

ADGSDAPEIGVVIRANADLSQNTTVSLRMPPYGPSVVSHYQQEEGWPAIKIFDGPDVANC

SAECTKLRLCESYTYWTSTGSCKLYWRLTPMSESADAYSGLAREPLLYLNRNASRSIGST

AALSGRAPFATATPNGFELHIYVDDSIVEVFKDGGLETMTGRLYISNGADTTGVAVYAKN

LNNVTVTADIEVYTMDTIWQAPVANAARNFTNSLYNLLDALIDI*

>Lp_070007700.1

MYELLLYLCILAVAAAVTFALFGEAGEADVAPADALTFFAGASSSVQSDAAPNNTHDASS

SASISAGAAHIQRLLVKCGVPSQQQEMISFLSKHCCVLPVESHTVRVLTHPTVFYEELKK

KVIAAQHSITLSALYIGDGPLSTAFVACLEEKVRWAAEGGYPFSITILLDYNRMQDRKNL

VTLKTLMELAERTGSTSSLPVDATASGSGPHGFPSSSSYSDREDVEEGDEDLEGEAPGNT

GRATATAATGVKVRLFLYQNPSRWNRLFSPFGRAKEVLGVQHTKIFVFDQRHTFLTGANL

SDDYFATRMDRYLIVEDNALVGRWFTRLVRTLNRISHPVICRKEFTQTFPEDTFLVGDDD

RADAARKRASSSSSSPPSPARRMLEKAGRVVRFPASPSTKEGSPTLHRKSNLVILPNAVK

MDPSTDTEAYCAKTKELLHGFADWARQLCARQQVDWSRYDTFLFPTLQVGRAGVYHDSVM

VKQLLRLSTAEDHIFLTSPYLNMYSSFVDEVLQGSSYVDCITASVQTNGWNGHRGMAGRI

PLFYLQLERSFYYLMKVYHCLRRVHIREFSVEGLTFHAKGLWFAGHQPSHPASPALAKAA

AGAAAAEGHPPPQASVDDMAAACCPETISAPYLVAYGSTNYGYRSVHKDVEAEAFLLTTN

DALRATLRNELLFLLKQSVPVTEDSFVGTATGRFQPVISLIAHLGQDFL*

>Lp_180007300.1

MPAIGQEKLEVHKCFSILRTSAEMMSDRKYKVAQNVVPGSLDEFIERYVETVSVPTEDAA

AARDPGANSPAAPITRRDKKVIRRDKMTLACERELGEQNVRKAVVYFCPPNHLTSEVVKK

IAEDALEEQYQRVIFVTPTRPNPIVRKTMDTYNRSAQDLRFELFEEDELSVNITHHELVP

KHTPLSEEELKEVLHAHALELPQLPRILSTDPVARYYGLERGQVVRIERKSMSAGLYATY

RQVV*

>Lp_140013200.1

MRAAPRLLVAAAVAALLLLLSGTPSHATLQTFIEQYQSIMEEAGSGFSMFYKDAVARPTA

YVWTCQNATATGATVAYASATTPYQLTVSTVALTVQPVTMSSSVTYELYGHSCDFYGSCS

AMQKGSAPAFDATPNHVVMVMGRMTSGGVVNFLPMAYYVVLPIAACSSAGSADVANVFRR

NGAVTLWRDM*

>Lp_140013900.1

MACRISCLSARCLRLCRLFALGLLLLTTAVAFIEKTSTGAAAAATAKKEEEEDAIRLLLA

AGDKALGQGRTSYQDALAKYTEALTRSPANERALYSRAELYAMMRDRHAALEDLNTLLSA

DEDHAQALALRMSLNMQLGNLVEAHRDGQHLAHVYTSQGKPAKVKSVMEQVHRLEHYTKR

WTALADLWTQPVETFAAAAGDPALTHRYEDCILLLESVVREFAKDSVELRLRRAACALAI

GNNLIASQELKQVTQRDANNLDAIALNAQALRGLGALDQAKSVVRRCLALDPEYAPCANM

HKLIRLQQRLTSAIEQAIKEKAYEKAGKLIDQAREGEANSPYEEQLATWQCEALVGLRDT

EKGVQVCQSLIDRSKGGNSPAVFDAHIWLAELHLLEDNIAAAEEELQKARELRPGDGKLH

ELQAKIENIKRNGARKDYYKLLGVKKTASTQEIRRAYRAIAKKYHPDQLRSQKLSDAERE

KREKAFRDINEAKEVLLDDEKRAMYDNGQDPNQQPGQGGGANFGTQFPGGFPGGFPGGFP

GGFPGGFPGGFPGAFRQQAGNGGQQQFFFFRNG*

>Lp_160010600.1

MSCLRQKRRSCAGLFSPSVWLLLVIVAAVLCAAPFSARALASEVSPDLATSTDAPSSHNN

ATSESAFVVFGYLPEYRQLRFDYEAFFKAGLTHLIFFSAEVDPTSLQLIHVEDRLPSMDK

WRTIRTLADKYDVKLMLCIGGGGRSAGFPLLVHDVVGRRRFISQVQRVVQQRELDGVDFN

WEYPDSMTEWLSFGQFLAELRPALRHTAAGAAAVRHRRDIVRSALITMALHPHPSVANVL

RSSRVLPSLDYLHWMAYDHIVPGEGHSSVAYATAVLQDDMIGSLDDAVHNARLAQLRKGS

KTRSTAAESAAAANTTVARKPNEVDHRRKLCLGIPFYGRHREDRRLPPESYEHLWQFIKQ

WARKRQPTWQEGGPELRAMSNYAGYDYNGYDDVKRKMRLARSAGVAGIMIWELGQDVPPG

TSPMSLMTAIHEQLAAWQKRKATGKDETWNASVSQAGATHRKEEATQLPRTSTTPRRPVV

QQAGRQAAKEDAALDGDL*

>Lp_160012200.1

MKLLSKIIVGTTCVTIFCGSAYMAAITLAPPPNYTVSNNDRLKSYDVLSVNNLYEKKTKS

QEFYLGLSRWRRKMLRDEGLLHGRVLEVGAGCGGNYAYYPHSYFTDDPEVQRHLKGYFGS

EKQEEGAENGENVTSYVLQHITKENPCDEVILCDRSAGMVQSCVSRIQSRLGYVPFRYPD

YNVAAIRSTIEARLGAQQRADSAGGEAGDKAAVRKRKIVHEDGTVQEVLVTPLSGNEEAE

FLAATAENAPGSASDVIVPVLHDLDAGDKLPLLSKEDEAALAKREQEIRARQKFRLRVEN

QLRREEKESLLRRGAADSSSVGTTAEKTRVVATPSELLDEHHNLQKQPLFAVANYAAEQL

PFPDNSFDTIVDMFGLCSFDDPVRALRELSRVCKPGGKLLLIEHGKGHTVRVNNHLDKWA

PRHAKNWGCWWNRDIRRIIRLSGLSVEKWESKHFGTSHYIIAKPFKSMEEWDKYELQREQ

ELKKAVA*

>Lp_160015500.1

KMGVETSLKCTFAAFLVAFFLVTFEAPRVPRGAITTQPRIVLDTTKLDYGYCHSSPSEID

EGTTSLLAVLNRITAAPFFRYFKVNSNRPCPYWAVSLLCTSAENTCDVCKCDEDSIPKAL

HADEDMSTVDTPDGSVLGAVSRPGNLDDWGIWQKTDNGAEYVDLVANPEGNTGYSGPLAT

QVWRAIYAENCLSLEVDGMCQEVSILRTLLSGLHMSINVHVSTNFYKDPELASPQHNAGI

YNNDNISFYPNCEMYNKRIAPFSDFVGNLYILYQFTLRALAKAKPFFLSDFRIFNTGLHG

EATPADLQLRENIKHLFNARLLCSPTFDESAFLESEKGRELIPEFKSIMLNVTHLMDCVT

CEKCRIWGKLETKGIAVAMKIIMSRENEIITLDRAEMVTLVNLARQLAFSVRNVQRLGVV

CKE

>Lp_260010300.1

MPLSVLTPHRRAALVSLVAVVLACCAISASAAGSLSYYTASQQTNTRLFLASFVDTMPAL

AQHWTGNDFCQWIGVTCSRNGVSIGASRMTMLDGTSVQLPNVPAGVVASEVMLTTFSFSK

ANFISGTLPPSWGTLTRLVTLSVGGNNMSGTLPDELGNLASLWALNLFDNSFTGTIPASW

GQLSACTLIALYNNSLSGSLPVAWNQLTSIKTIYVQDNHLSGSLPTAWENMTSLKTLYMN

DNNFTGSLPAEWGNMASISYVTVANNNLCGCVPAAWTSKGVTILADNALTSDDCATANIC

VQPTTTTTTTTTTTPTTTTTTTTTPEPTTTTTSTTSKAPQPSQCAVPNCAVCNSQNPHYC

KSCKAGYELTPAFMCRERAEDVASAPLYGVGTLTLVLAATLIAA*

>Lp_270007500.1

MTLRTLQAWRPLTRGRFFYPSTLVVAVLTLLLLSILAAPQAVHGHTVSATSSLCPRCPLC

ARPQTSFRDRLKYCHSTSSYAFAFFSHGRSELAAATQRTGASHREQRQRYPLELPFATDD

SVRARCMRDATELLVSRVRGQPQVVRPLLDVLRRKLAYPREPVVVHLAGDNGVGKTYTAR

LVSQALSLRCAVDRDVCDAGDSLLTIAGTSFDGMAVSEARARIVRQIMAHVEHYPHGVVL

IDDLTAMDPALVSALAPLFGRASHFAEQLTDLPMETAGGHRHSADPSHANTTSRKSGADA

RRLLSWPWKVSTFAGKSAAVPPSLSQLLVFITTDFGRQGRTVGKSRADIEAMVQDEFAAL

YGALLPAYTRTFIFFPFTTQMAEEVVRSAVTNLPCALGEHLIASSWISDEAVSFLVEQHR

GTWVGKENGHALRRLIEDELISQLIVYWEQHAVQERLLVRFELDEAEMRVVLRLPKHRVM

DTLDMGATGSSPAADAGRLTEDGDAAHDQSGEDDL*

>Lp_280008900.1

MKRGFSAVLQLLIVVGVFLLTSCGADGAPRPAQQASTAAPAASAHAAPPSPFGPLVSTFA

SLNAYQRHLSFSTNPLVLFLFDHDPQVILNTWQPLITSFSQAMEKFGIDVVDVATNSEVG

VQLQQAIGSSTAVLFFQGVGDTPVDGEKGLKVPIAYEGAADLVSMVKWALSCVSPAVVQR

VRNEADFARFFQLYPQYPTLPRVLFFPKNNYTHVGFLVVSQHFSYDAGFAVVPDAFATDA

TAAIAQRYGVRDESELPALFVLHKAAADEDGGAGESDYAVRMNTTATEWSYAGVKAFLDT

QLTERIEALVAKVEATQDAKVLAVAEARRKYMAAALVERQQDVVDEERLQMAVEPVVVTD

QATWVKHCLLLPKEHKCLVAFVDSTQDPAAASNAVKVLSLVSLRLVELLGMEARSIGLVV

VEQASSEAVRDYFEAGRNGYPDVVLLSFARPARYYNFVGSFSAEGVLQFVTSHDSRVTKA

EVSGGHAFIPRMVPKLENPTAKENGGDDGDL*

>Lp_310011600.1

MSQRTVLVAFVGVLVLCVCLARAEIFFHDEFNSLDGWVQSEHKSDYGKVKLSAGAVHVDA

KKEQGLQLSEDAKFYAVSKKLPTPVNNDGKNLVISFSVKHDQKLECGGAYLKFFSELDQK

DFNGESPYWLMFGPDACGSMNRVHIILSHDGENHLWKGSMRPTKDQATHVYTLEIAANNS

YQLYVDGAYKAGGSLEKDWDIVAPETIPDPDEKKPEDWVDDQMMDDPADTKPADWDDEPA

TIADPNATKPEDWDDEEDGEWNAPMIPNPAYRGEWSPRQIANPAYKGVWAPKQIPNPNYK

PEPNLYRVPAPLQYIGIDVWQVQGGSIFDNIILGDDLQEVLKLVKSTYGAMAEKEKKLLK

VIEEEKREEASKAKAEAEKAAAAEEAASEEEEEKEDEDDL*

>Lp_320014600.1

MAGLDIAWIPFCVMYLLYLVPPTLMEVYSYYSRGGLVRYEDIKYDDTFVPQLVWNFDTDT

YYKRFLDRPTSEAGRAFFSWLDWSAESYDRTIYMLRRNYAKMSGEVSFWKWYVYGPYVFN

WPKKTWLRGDDAAGKLMPAWGVGGEERARKEYAEAKAKGFDKHYHYQIMRKIRREQALKA

TQQQQKPKAIE*

>Lp_330016200.1

MNILMPLTILLVSWSIDSTDPEIQQRIFYIFCAVHVAIVFMGVYIFVRIWQVGDNTLIRV

KDAYTSEESIQRYWEYDLGNLRDLLVTKIGISAALSAFLASRYGIPFPLLLQSLNNPKAV

YQSELFRIYVKGEKAEGELQRPWTEAGMMPEWVKSLWAEGEKDSEKFLATGGGATAAAPV

TSKASRKRK*

>Lp_340007200.1

MFCTISTRLWVCVGLLVLVCGCWVSSTVALTSTNAHPDANTDTAGKKEEDPVQVFMCADS

GEYSGAHELGILQSSMEQAEVLHLPHCQSAQLGYIRHADGHAIRSLTVTTGIGYVSATLC

TTSVLRFRHQRGVAYRSIVYIGTAGFSPMVGGLEPTDTAAYADFAAKEQQALLNGEADRL

TDTTAGPPALFKTLAEVQAMQAREVQQRKRAVESGAKFPLPYPSQLNFEYTKTGAQLEQD

GCAPRLASAVTPLAVGSVCVTSAAFLMESGSCTERTRHTQCSRPHCSGFQNSLSSEAKMY

FAKGDFAKEIEAASEGRAWPEMPALVRAGLQRFWAANEAMEPAGRVAPSSPSFVTCAEST

VNAINVGAERDYLCREYTAYTLNTVRRQTAGPRRDADPVAAPLTVNEVVCVQAMEAVGFL

RSMSADASAAAIPVAVLRTASNYDMYPLKKHYMPPAVWREALASNQVVVSPAVANAIEAA

SPAASADEAAENDIAAYTWQQDVGFMPVAEYEEFVRASFRYSIKTVTFVVSNYFFGGRTF

E*

>Lp_340008500.1

MRKKPRRCGRRLALAAALLLLFALLECFIRYSVSSFIDEDERDPFTPYFDKNLTRFGERY

ISLEDFFDCLDARLNVDHRTKTERAFTNANVTIPYLVLPVTVEQGDMKALMCNLTVQIRH

LMYVQNGGVGSMTAFLDHVQAAFAFTSRLKLPRMPQNYGYSVALNVAMRDALSLPFDEVP

FLMMANNDVRIPGGMLEAGLPQMYVHSLAGRARVAELEAEVATEPNEYTPVRFRDVPLRS

TDGRHVLVT

>Lp_340024900.1

MRATQTIAERRVRGKVALVCLITVVLLILLSSKASAGVVQESVASSEDAEWTHEPGYTAE

PPTDLPATEAPGFVDAAAPQTRAQKTPPEYQQGTTAEEEEQTLAAATAIAMEETRKRIQE

AVRQEEEAERARLAAEAVQHEAREEDMVHRIQEEAEQPVHVRLAEEARLTEEAKKLAEAQ

AAEEEAQAAEEEAKRLAEAQAAEEEAKKLAEAQAAEEEAKRLAEAQAAEEKARAAQEARL

SEEVRRLKAAEKARLAEETRLLNEAARQAEEALTAQGEAERLELEQQAEAARRAAEEKEA

AARQLREQQGELDRLSQQQKDTVQEIQTSSEGYRIALAALEAREADELAAAQQREKLAAS

SFQQSIERELEALSQNLATLDARTAAVRDRLKETATPNKSLDVVLTHLNELFEDVDKASG

ALAAVHVTARSLKKAAKVAESSRGASNASAGPESFKERRSALAASHEAALARLQRQLQAI

DAKMAELRHKLGLPTVDEEAKQNSRSVTGTAEAGTQAEVLTAALSSKPTLSQETTTVAPS

LVTTSAPSSTLHPASGVDAAASTPSPASSTAPHSTPPPPQVNELRPNSSLQLLIRTLVQQ

AEEKGYNVSRYEVHRRENGVFFKAKGRLQIIGAMVLITLVSVVQTMRWCRGDRDDGHAEP

GTATKAAAQKLLPAKAAPATAGAAQSKQDSTPSPAGQLETARDAFPPLKPSSGAAKRESS

YGASSLRQRRFRDGASSSLAANCDPVQQLPDSSSGVPLSLPSPTDSPHTQLPTPMMPPPQ

GSAPTPVQHPSADVQGDSPNRFPAPPPSSLGGAAGRAASHSMMDIPPPPPPRRRRDAEDF

ELQNPFLSRWS*

>Lp_350010100.1

MSSNFTGKSLRGNLAVALICLVLITFAGVATSSVPNIPSIPNVSTSSSSCFAAVPFYTAI

QENNTRAFLDAFADTLSVLRSLWTCTNFCERRFITCSPSGVELHIDDLSVTGSLPDVPQG

VVGSEVVVTAIKISGNGHNLVGTLPPTWGALEHIYILQLAETGLSGTLPEWLGTKVNSNA

LRRPISSGSLTNAVIIPRKLKVVNMSYNNLTGPLPESWSSLSGCTHVLLNNNEIEGTLPS

AWSNLTTLRVLNLSTNKLSGTLPSSWSTMKAMEQLSISNNSIDGTLPTQWSEMNSLSTLE

LQMNKLTGSIPPLWGTSLISLTEIYLQNNYLCGCVPDQWKTPSRISVSVDPPVSSISCNT

ANACEDSSSSSSSSSSSSSVDECTAPITI*

>Lp_350014800.1

MRSLTAFLLIVVVALTLVAAEMTEDDFRKMKVKDLRLFLSARGLECTGCQEKSDFVRMAY

QYRSLNPAGSAEKRAIPAKKFWEAWADIAQAECEKAVKLRSNDPATEPFKSICSTIHSAT

DSYLMQHGRKVANQLKKTPQDLLQTSFKDIYFEAGSHLCQILADYCLASPAAQENCQSLG

AVMSAMDGVSGADFKMWTTNVGIENTNPMYEIIDARDDL*

>Lp_350037900.1

MKWMPKVVLLLLVATLALVASTCCAHVIGVDFGSEYVKVTGPHSDRGLDIVLNELSRRKT

ENFIGFRNGDIYIGDTAKSLAARFPLCTASAINQLIGIRKDSDLHSAFQELQYEYHVKFN

KHGSATVDICNVKEPFTAEELFSLMLSYCKAAAEHDDVILPTGVVITIPFHTSPVERRSI

LDAARFSGLKVLGLMHSTTAAAFYYGVRHRGFGNNTVTLLVFDLGATHTEVGVYQITPAP

RRPPLQNAFGTLRTLGVVEDRSLGGRVFDLCVARIMEKEAREKLNIGPVLGGTSAAQLKS

HFSLLRAANKVRETLSVNSVTPYTVEGIVPDRDFHSSMSRATFESECADVFDRVKNLALN

VTKTLNISLSELTAFEMMGGISRTPKIIADLSQLLGREVDRTMNMDEAAALGAGYYAARL

SPLYHAKSLKLDEAIPYSVEFEVHPQLDRKKALTRRPLFGVDGLLLGEPVSLMFNRTDDF

HLDLFSGAGAKAPIATIEVSDVKSVLTSLGALSFAVKHPNNSHMIRMQLVMNETGLIQVE

HTEAVVRYAAEVTQRTRENVTDPTTGEVQEVDKTSTVIRMRHKTAAVPATLVWKNPAELT

LQESRASEKKLADIWEAEHVKHVRATAKNNLQTYIFWIKYEGVGDNTEITSAANPAVLQS

VMDEVAIVQDWLEDGDGSSDHCPTEEYEARHAKLKELIRQLLSEPEPESPAGTTKNSTDS

SPVANNDVNDDNGDL*

>Lp_350038800.1

MFFWMVVALLVAVAIVVTPAGSLALAKAQVYAHGAKFMFQAKDLPKVFRGTNYLRMDAVG

DYSGGKRTIRIVFVRHGQSVWNSLFNSFNFGLPSRMVKAAIREFTDFFTNPFASCIIDSP

LSSKGKKEAEDLASFMRTAKTKVSFDPASSVVVCSNLRRAMETALVGVSPRLTVTKERIV

VDSTLQEGSQNIDAQSFSTERGKIAPCTMGSITDPAKLGTVFNPYLNAGNRIAGVDVYQR

MDEFIAHLFGGHGERSLVPAAGGSNAALQEVIVVGHSGYFRNFFRRFLPPSSTHVSKKKK

LQNCAVVMLEITRDQNSGEVAVDESTIKVLYKGF

>Lp_360020800.1

MRSFFAHRHAVFVVAVLVLIHLLNVAAVAHAFGRARSTKEPASVGPSHPRVNINWTEQKR

NVNFTKEERKRILQNKLVLTKLIVLNRHGHRAPNAAYWQMCPNDAFNKRRYDVAPEDLTG

LGMQEEYHFGQYIRNSYHTFLGDKFNRSLHYFRAVGEPRILQSAMAVAQGIFPNGFGPGG

YLPSRPQFVPIFSDMDTHEYLLDDVPCFRRAENDSNRWLEGRYPAFIKEPYNRKVIAAMK

ELCGPYNGTAKGFAYIKTVADGLTFNTDFGLNVLNGKLSRQMLFDIRNVSLQLLLQRLYN

TDEQQTYTVVDLPRRVLQAFNHTHVGDSPAQLNDFSDTRQESTFYFMHREALYAFAQFFG

FQYNVAGLPPGEMPVAASLIMEKLMPRGDQFSHDARKIYIRITLWTPYNGMYSVPISKCR

IPELCELHELRDIHDERVKRTGTWEKLCQYVPQEIDHTTDIR*

>Lp_360034100.1

MPVFSCVYAAAPSRVLLLLLLPFLLVAAPLGGASFTGIAEGGTSGGATGGALVTYCEGSG

GGAAAAPSAELGGCQRKLVIDLTLDDSTVAGSVLETVVVVSTALHKAVFSKAEINAGATT

TTTALSTTLQVTMPPIRIAFRRGGVQMRYRLSYIREFAAALRERVRALRMAMSCDDGVTR

CPSYTPISPTSSSSSSSSSDSGSVAVVTAPLGVCCLCITVECTLNGALCNESMRTYFCFR

SAAAGTICVNEEGVRYAGWSISAGTPYYAMSTEVSGAGITAATYPLTTDSTDLERGTSRM

QLLQTSGVNVAAAGQLLNVSQRVLFIPLSGERAAAGVAEWMLVPTSLVTASGKACDKVGV

SAEHFYSLSSASQCNAQRGTCLANQLEDLRAADLAAVAQGGGGSYLAAYLGNFTQQSSGT

QRYLLDEVERSGGATLRWSVNADALAFTPVPISGSLIAATYAVGAARVSVAVSNPNVFAG

FYYVAVGNCSDSTRVIHCGGGDDDDDDDDATQECVASLLV

>Lp_360042800.1

MGVVRSKQMPCLTAFATLAFLLLLSCGGFPQIAKAWSGFDAYTIVVLNDFDTDVALQEQF

YNAPTGTPFSLSRTGYRVLDAGAADRPTVGHRGGNESLHSAYPLDPALSTGAAFDDLKAS

DAPLPFRGRFSGEWVVWTKENVAHPFGTAVAAEPLLLHCRLPLSRNRTEVQLQQRRYIAQ

HAAALSEALEVWWHAAPAAYASRCLYGDGEKTLTGTERYYELCPGGPVTRVQHHRLTALL

LGSGGDLVRDTKKRGRRKGGTALSVLAFLHQMHQSTSNRNAVLRNPRYRGVFEEIGPSPP

RHSKPRWNSEHLVWESWYPSTQRCTTHYTAVSSSRKAQPFDAAGNASQRRQHSKIHPEDP

YWKTVVRFRCPSQPNADATQQVTTWTVTEARQLCEYHVELMSVLVCGWEQELDSFNVNPV

PCVVLD*

>Lp_360044000.1

PQLHEMSQRITTAFAIVFCLCMCGAVLPCVTARKELPVKTAFPPVVQDAIRCDVCSFIVA

NALNYVEEKRDELRAKRLPLREDDVLEETENMCIPFKDEGQWIRQVSLDVAPAGPHDASQ

GHRVEVNVVNFYGHCGRTCYTVAALCEELMDRDVMDDFPGELLRVSKDANMTDKAHRSAV

FGKFCYPSRHCRQHKRYVVALEKVLRRDAELLRELKADKPRRIKNEEREVEMMMYRLMRE

QHQSADVFSRDDIRKMQHAFIHGTKEDVAAIDPKAFDLSDEEFHVLREHM

>Lp_000047100.1

MAAFPTRRCAIALLCAALVLVAACARANSIVVNGDVSQSDNAFDDNSITFDLGSVSADVV

TVQLINSKVSGSGLSIVGYEDSVPSSVTTRVSMSVTSTTVTQSTIAFTGVMPPNSDIRLT

ATTATLATAQSLFDFSGLTLSGNVTVTVEDSSVAWPSGSTNTGSIVTYTAGATNIGISNK

GALFILNATAVNGASVLHIATSSVFSITDSGVLAVDYGGCDGCSSALVTIDVPLKVDGTS

MFRIMHGVVGNGAKGLLASTGYVTVSGQSLYLISDSTIDSGSFFDYHVSGNANDSTAFPF

TVTSSTVSFLNLVGPSLGIPDGAYVPSTADSSSTVNGGGCTIGGTALTDTSGYLSKGLKV

TQVVNSNGAAGGTCANANCVPGYSSSGAAEADGVACSCICTAKIYNPPSCTTVSDPTQNY

HSATCSLANCATCSLLYPSSRCAQCNSGYVLSASYQCELDNPDNATTTTTTTTTTAQPTI

TSTALCSVAYCEKCSPTDGSTCTSCRNGYKLTSGACIANLNGAAAAQSALLAAVACAAAA

AFYVL*

>Lp_000404800.1

RFCLSLLTALAFSAALQVVEVCGWGCVGHMLVAEIARRRLDEANVEKMEAMSHNFHVSGP

FSSSPDMVQAACWADDAKQWHQYAMRAWHFVDIPYNPENVPVEEPLDPENAVSASASMIT

ALQHPKAPLYMLNFAWVNLVHFIGDLHQPLHAATLYSHRFPHGDRGGNAITVKVKGHQVK

LHALWDDICEVPGPNYKRPLSDTDAFALAATADRLVDSYKFSASLRRVRELATMAEESHE

FAVNSSYDGVEPDEQLDDAYLLRCKEVAEGRIVLAGYRLGYLLNDLLHNITVSDEAVSAY

R

>Lp_030007000.1

MRRFTAAAVTLPLISSCARATQQSRHFTFATLPNLSAHTASSANKPTAASPPEVCGYGAA

SATSPLKYMTIKRRDPRPNDVSIKIEYCGVCHSDLHTVRNDWGMTTFPMIPGHEIVGHVT

AVGSDVTKYKVGDRVGVGCMVDSCLQCEECREGLEQYCIKGNTGTYNSMTEDGVTMGGYT

NHIVVREEFVLRIPDNLDLCAAAPLLCAGITTYSPLRHWGVKAGSRVGVVGLGGLGHMAV

KLARAMGAEVTVFTLSADKVEAAKKLGAHNVVVSKDAEQMAAYKRRLNLIIDTVGSAHDL

APYVDTLKVSGTHVLVGAPDHPHPALQPFQLILERKGIAGTCIGGIAETQEMLDFCSKHN

IVCDVEKIDIAYINEAYARMLKSDVHYRFVIDMATLKKPK*

>Lp_050006100.1

MRACPLLRLAFLLATPFDHQPVVLPSIPTAASVVQGIVPAKSELSRVDELQLLLRHLPEA

CNRAKELQHDAQYASRYYRARFHRGSQLQQKQLSHVVALNPQKAQQWARQRLHPVSDPFQ

CS*

>Lp_050006200.1

MLRSSCTLLAAAVTAGLERVEAGNAAKQKAALQSIQHQHNEGVARNTNRTITVVNTYDTK

TALFFDRYNLAVLPSVNGGAARQPLRTHLLASGAVAGVQAQIVSPWLAEGERRVVVFGCR

SVHYLRRQLETDTTHTYLVVDSSLKDLTNAAAALTPEFPQRVFFLRSESMFTCLEVLQPD

TVDVALVPMPVPFWSRQGSHRRLVHFDFYGAVHRALRERASAADPRGIVLFTDCEPYAAF

MMEHLEEAKLIVPYTRKNPHDLFRRWLPQVHTVTIEGRGTRQQREFVQQREIEVVAIAAA

KSGPTTAEAVQLLSSYNYARKYYRDFATTNTPTAS*

>Lp_050016700.1

MQPFRFVTSSLTLLVCMAVVGRTLAAVVNDDLPPGVMRGPLLHEHSFQEPLVSDWWEEGV

PHYMIGGSAVANERFLRLTNNDLDDHGFAFNTAPLDHDNWELRLRFSIRPPPPAADMNGN

STHYQGGDGLALWYLDQPIGDDHQHVIKYSKSLSPELRDEILDADNPWRVADFLLNTGED

DDDDDDVDEDDLTEKEKKERRLEKERREEKRKKREELFKRIFKRGTSVDESDGEPRIMGV

KYSDFNKGFGIILDSVGDEAAHIDEHKGKGDAVDHHHHSASIYLLLNLPDHAAGSGAKVT

NNFNPADDNFRKSPEVLRCEYDFRQQGTKPFDPKARLMDAVTRAATAAEEPLELVVRYYK

KKLTIIIRREDVAKRQVVAVTNEKGETTNEVEVTRTYKETMCGEVFPIQLPLKYHFGLSA

STGHRTHRERRSRRIQSVFHAADKNTFTHVDVHDVYKLELRELGLDAKAMGYSKTIPLEH

FDFEADKREREHFSRQIPVEPDSDTAH*

>Lp_000039500.1

MALVVGLFVLAAPAMAAEKEASRVVELNRDNFEEYVHGPMRHTFVLYCVKWSRQCQTARL

AWDRLSISQSTKELRDVFTAAYVDGDRYPDIIGKMNVQGFPTATFYTPIYPEGVEYGGTR

EPFLLDSFVFQFS*

>Lp_000058200.1

MFRHLQRISLLLGLCTARWAHVINTVPTNAAEKDVSSSEVVNVAQIVSETVRTTDENNSL

KTPTVDSDDVSVMSKRDNEYIELHKLQYMYSQDGPFIGSRTKDKKVDQVSLRKRRHIVRV

PQPEVEYRPNETSFRRLPKHYILPNEFELRALYPMSTSLKLIEGQADCVDHNAFDVDEDP

APIAYSMVAPPDWKPDVEYPYMVVLPDHRGIPRDFEDVCANFFERPAHREHMLEQRWVII

SPVVNLRHNMQIPVEGVVARFCDWVTDNFCVEHGKVHLFGKGNGGYVALRTVLENKDLVL

SVTAILGRNGSPFRPLDRAQDKVKNFNGVHSLVYVPGLLRKQDWYYKFKFMLDMARVRPP

IRNVHFADVRDHQVYYAINPQEFWNYMKYFRQYNTKMITESGYAV*

>Lp_000063400.1

MFFCPFCSTLLLVETLPDGNALQCATCRYVHTVASTRGIVATNVSGEPVLTIHHSFADQN

KKLMDAEDEAGNSAPPTVAVAAASSSATATNTDASITGTTAEELAGDGSAEGGQIMTIPC

QNEDTPCDSTKAYYIQLQMRSADEPATVFFKCVKCGYQWRQD*

>Lp_000081700.1

MRALSRITFTVAAVLVCLSSAYANGPISKLQCSTINPNPIVTNCISYLPPIVDLYGNLPY

DSIPAYQNCVKERCVCTGAAVNTDITTDGIYCNESGWMETGYTTCSRFNHCFLNFWRCIN

TAVMYRYDHNRKELTEGEMDMAADIVAHGRKPGEEFELTDTYRSCRLKMCAAADSRENCG

LITCLPNYTQCNEYMLPPPLPYHHQLCTQGCRAVLLMMALTISIVSFSLCCVCCCPAQVI

ITEPVIKENSSKGKKGSDDDDDSKTGSQAATSGRNSPTHTNHNSHISSKNENKPASHGSN

AAPKETSTPHTEAQQRGDGNQPF*

>Lp_000097500.1

MSFARHVVLLSAGLAAGVSTAPYFCTAASDPTRFVDTTATLTDPLKRNRRTGELATFNLC

SSPMPTSGLEPGGGGAWSSVVEGACRVITLVAGGGAAGVRAAVISGAHRTALAASDAATT

TTTADNGAADTAAGITTSKPNTAAAGTESLRAAPHKLKERFLRYAQRDPESGELVLTLDG

FVRCMLLLPVTAAEEVVPALYPGTRRDVEHHSPQRPASPSLWLQRLPEAVRRRFEHFFHC

VDLDGTNTIDYAEFVVLFTFLSTRQQMLKRAFHVFDLDDNGRLSEQEFCRLLNTIMVDPA

VQIRCTPSGSKNNGEANVAGALGTSPSITNRAGESRSSKRQRQRHKDELSFEVPSDLMRP

LLFGPLPLHVGAPPGSTHYNGLADATASVSSSSSSPATDLRGLQKDAGALAGTRLSIPKT

AEPRNTAEGFDSSEAPDVKPQKGSASLNAPAALTSSFSSLWGGVCQGVQSAWSQLWRHNV

FHPPAFVSNATAAAAPSRMAELQEMAAQDTLLETVSYPTFLYRMDYLRWELRAIEFGLCD

PENKGSISVEDCRKLLRRDRLGITDGATAKTPTAGGSPTRTTTAAAVATNTRPARELLQT

APVTWQVYQKLFDVIKESNTILPALELMLESLPPVPKDVLSGGAIPDAELKPATLAVQAV

VQQTVLRYNNSRATEADNDAQASIEAAEQESSAKHISETEMEKQAHHSAVSAPSTEATAQ

EHYNEERFGLSSDPAQSPEEKQKQARQLQATLVRPTALTWTQFGHAVAAVSTIPPLSKTE

EALFRALLDEDGSNTLSPPEFARLCALKESFFADDLPRFDEPKRNTVQQFFFCMQQLE*

>Lp_000102700.1

MKHILALFGGFGAHALSDALMVNWPLYLVPASRTAHGRAVQVAYLGIATYDLPEPQKQTG

ELTRHGCCVRPTRASADIIVVSCGNTFCAIQRWERLGLDVLLRRVAAHCDHRGRVVLAGG

SAGAICRFAAGHSRSANPATFRSAMLQAAAAMTTASSAHAASSAKTKGTWSNIHVHGLGI

LPGLLCPHYNIAQGDRAVRRGEDFTELLLRHPTERGVALDQWAMLVLPKDGRYEVFLVPG

HVHQAVCVAGALDPCTDAPDKQAPGLS*PSITDDGRVQRRRVAVAGNMCDLLRAPTG

>Lp_000113900.1

MAWLNPPLRRMAVAAALLVLCVVDASLVGARTASDYTAAQQASTLQFLQGFVTANPSLSA

VWTGTDFCSWSYVSCSLYSPSLLDFSPNSSPYSGTLVLPELGDDVNGSAVIFTVIKVRSM

GGRVTGTLPASGGRLTLLTL*

>Lp_000119600.1

MSLFFRRFFLVALLVAVAAAVVANAECDPHCDTCQLGVCVTCKKGYYRGADGCAPCNSEH

CAHCEALGRCLACEPGYKLEYVTGTGAASIFTRMASV*

>Lp_000152200.1

MIRLASEPQGGKGPKVPECSRMLFGLALLLTILSEGMMNTQYTALSTVITANTDPTALQQ

LATFAAMEDVAEEEHLDWGSSGRLVRKPTNTIVKPRNAGTPDPAKVAAVSPSHSRANSLE

AHSEGLSKSITDADSSSSHTVALESSAASAASSRRRPHRRHHRRSGALSDALSESAMTVS

ANVEAELTVSPSQPVHSYPMWTQFHVMEVAYSTGTNNFIHLTERFLQYQHMFPVVYRCDL

AKVVCPAEAELLATPRSGLAASSHWACANCGKKHEVVRESCLRCRAPGPYAKLFIGQAVK

ELDCTESLVRFLHATHPDVKLHRVECHHDTHAGGKLGRGKGCASVYVAREDVATLRDKLH

HNAYFDVDDATGDIIVYYVYTEQQQWLNSFVQLRNEQVAQRPLFLPLAPLVVEESVGAAT

AGPARKPGR

>Lp_000154600.1

N*GRHLVVLVACLIFATPSPSSFSSSRFTASPQRLLLLRRVCRCRRHCRLQSRLSSPSAQ

ALLALLTLTSLPTASVVPPRTQHPRSP*RPPACPTTPA*WPSR*SISSPGPSPSWPRPRR

SSRTPTPCPSTAATPCGATATSAATCRTIARVTPPSAAAAAPAPCTAPPPSPSWRPSSAS

SASSSASCSTRRSSCPSSSPSRWPPPASPARSSPGPASPACTTWPCATAPSTRTSTPTPP

ASR*WSRPGAWISLQWGCSASRRGPVLPRRRRRRGRRTTPPST

>Lp_000158900.1

MAWLNHPLRRMAVAAALLVLCVVDASLVGARTASDYTAAQQASTLQFLQGFVTANPSLSA

VWTGTDFCSWSYVTCSYSSDKLDFSSSKSPYSGTLVLPELGDDVNGSAVIFTDIKVRSMG

KGAMVAGTLPSSWGRLTKLQTLYLDGNGLTGTLPSAWGGMASLRTLNVSGSRLSGSVPAS

WGGLWQLRSVDLTGTGLCGCVPAEWAGKAVAADAALTGSDCAVANACSKVVSGSQSESSS

RKTKSG

>Lp_000172300.1

MSERVIRVGFLGAASIAWQAWAAIQNSGMVVTRVGCRDPERGKQFVKDVCHTLKISEAQA

PSVCSYDELVAADDVDVVYIATPVTTRDHWVKACVAHKKHVLGEKPPAVDAEQLRSWIEA

LDAQNLLYMDGTMLSHSQRVKDVCAAVKKLGGPVKHIYATYSWNASPEFLAKDIRLDPSL

EPHGALGDIGWYCIRYVLHIMDFEMPTEVTGRILKQNEKGAILAFAGDLKFEVNGVPAIA

SIFCSFNTAYEQVLYVATTEGTLQIEDFAHPITTRPDSLFYEVHNTSHEDVCISHTATNI

VTHTVPGETANGVRDALWRDAGSLLHEEGEGEQRRLKAEPHASRYWATIAWKTQAVMDKM

LESARQSSNAAPVEK*

>Lp_000202300.1

MSLFGAFLKAAALGAVANAFLPTRCVQCAPLLDASQTPLAEATTATQDFRASICPQDGDA

HWSVEADDFQNGLRAATAQLRILRLGDEYVDAEGNSAPVMRLVGLFPGVSTDVLHRHLTD

LRLRLAWDTNYTFFERFPGEFCGTLQGSLLQRPLAAVAKKRRHCEGDVCTLVPDISSRTF

DHGWFCHGVGSSSLRRFGLVDRLFQYERLSHAFHFTADTDVTPATAATTLTMYDILFSGS

KTAREAAGAASPPLAAWLKARRSRGDCEGVDVNFQHILLLPIADANAQILERPEQLQRLC

TMGSVLDVQTAKLVYSVFKDSQERTNNGQASLPASLLVMTSANNVSVPVFLPLWMQKKIS

GSVSRKAYGNLMAACLKEQGQ*

>Lp_000210900.1

MQYLPLLAVGVSIGRQTAAFAVESLKAIRDPTDADAVSAVGELSALNALECMKRCMMADR

RGRGILRHKPVIGDDVLEFSRTLPDTTFGHRYAAYMDHNHFKPSGRTAVKHVADPMLAYV

MTRYRQCHDFLHTISGCGRSVEEELAVKILEWKHTGLPLGLLAVAGGWPHLNPTQRANLR

VYAQWANENAPCSVHGQRQVLMYLNVPWEDMLAKDFDEVVAFSGITPLPVYLEKLQAPK*

>Lp_000212600.1

MKAIIAAAVGFLGAGMYKSSSQAEAWSWSNKPAFSHKEFSSYKLISVKDESHDTKVFRFA

LPNEKTRLDLPIACCITLRCTDEKGHEVVRPYTPINLANDEGHFDLVVKCYPDSKMTNHL

FSMKVGDTIEAMGPWHTFDVKPSQYSQIGMIAGGTGITPMYQVINNLLHAEGNMTKISLL

YANKTEGDVLLGEQLDAMAKEFAGKFATYYCLTTPPKRWTGYSGLIDKKMIEETMPGPEH

EGDSVILCGPPPFMKTISGSKDYKSHPPKQGPLSGYLKEMGYTEKGVYKF*

>Lp_000214300.1

MAWLNHPLRRMAVAAALLVLCVVDASLVGARTASDYTAAQQASTLQFLQGFVTANPSLSA

VWTGTDFCSWSYVTCVWYSNSLDFSTSKSPYSGTLVLPELGDDVDGSAVVFMEIKVRSIV

VRVTGTLPASWGRLTALETLYLDGNGLTGTLPSAWGGMTKLETVNLSNNSVTGTLPSSWR

AMAYMSSLNLSRNAISGFLPPQWASMSSLRYLYLSTNALTGSIPASWGQTWTFLFTAAFE

GNKLCGCLPSQWENNNIFVTITVDPAVRAAECAVKNA

>Lp_000230400.1

MALRFSCAFLVGGALHSFLASASASADSPNKPYIAPYSTNPANSRVYFDVAEQGGRSFFG

MTSQDPIGRIEFELFDDTVPITARNFRELCRGTSDKSPDGKPLTYKNCVFHRIIPDFMIQ

GGDITKGNGTGGCSIYGTRFKDESFDGKAGKHKGPGILSMANAGRNTNGSQFFICTVACP

WLDGKHVVFGQVVSGYEHVKKLESYGSPHGKPSKTVLISDCGVLQEMAK*

>Lp_000253300.1

MLRVFALSLCLVGVATQLCRKSAPSIALKKKKAVGRSHSKSRSRLRRKRSKSSVASRCTA

SPSLSTSGPPVRASVTRDRKSKSLLSLSGRAAVSATAAAVESSNTPASHRAASSTAPSPS

ALPLSKEQLLLATPRTLYTRLRSLLVQLVLLSPEPLKPSQIFAIYREVVDAETAALFACA

AHAVQLCTSEPFHRTTGRPLSHHSSKDNRRTSESSEHHHQGEGAVAGGDAAASSTGSREE

LIKIGQAFLAAYRGSIHPLVAALHHNAHEQHVRASADGVAWGKQEKRGSSSERPVVPAAK

DVVQWMLLHILFCDPEFRVSAYGGTVCYPALPSLLLATAHRMQDSERLKQCYALQWDASS

FSAAADSGPPSSTSDVRWLNGAAGSPPWTMAHALYAIYDIHHARKSSNSTTAAATETTET

ATAEGREKAPTMVAAVAPMSNSSGHRLVDYFICQAASTLCPAGAAAAMELGSDKAAAKAF

PSMGWNEGTQGSCMLQQYVAGFADVDGPLSFHMEQNATRRQPSGEFYHSNVRLMPDLFRS

LVPVPISVARLSTLLRWNLSIHLAGVFRSFFYYLLVMGANPTVHRLTRGVQQRIAATAYC

ADLFGGQGHGNRGRHRRAGEKDRAAVEQQVRQALLLEHDAAVTAAAATTTTLKGDAAQSP

PTSFPSASFQAALRDAFAREGEVVHHPPLTDDRVVELLCLRVLPHARPGCAPMQQQKQHP

RASAVASTVVPSSFRGYPLTYIEVLPTWSCTPEETLKRLRAKYLRERAGEEARTNAQIGV

TTSSGAGWLEKSEEPYILVYAMNSPAACGRLRVVAERFASVVTSGAMTLRRLSVITLWAH

EYGVEMAAEMLFVVLLLNPQSARLHPPAPAHAESASVAARRSYGDWTVEFI*

>Lp_000267400.1

MAWLNHPLRRMAVAAALLVLCVVDASLVGGRTASDYTAAQRASTLQFLQGFVTANPSLSA

VWTGTDFCSWSYVRCGSYSAALPAFPPCRSPYSGTLVLPELGDDVNGSAVIFTDIKVRSM

GVRVTGTLPASWGRLTALETLQLDGNGLTGTLPSAWSGLKSLTTLYLDNNQLSGGLPPQW

RFLSALDTVYLQNNTLSGTLPAEWGRLSSVHYMYLNDNALTGTLPPVWGAAAMLRYLYLN

NNQLVGSIPRLWSDMRSISAIQLMDNRLCGCIPQGSRGSLLSYYVDAAVSASDCATANVC

EVQSATGAEDSDAGAVLTANEKQTLAFLRLVGAAVGGALQSSWSGVWYCSWPNVECDASG

LATVDLSGVALSAAVQLPELTEDIDGGLVDVTTFKLYGKGAMVAGTLPSSWGRLTKLETL

YLDGNGLTGTLPSEWSGMTKLYTLHLGNNELSGSLPPQWRLLSELYSVYLQNNTLSGTLP

AEWGRMSSVNNLYLNDNVFIGSLPSSWASLSSLSNLYLSDNMLTGTLPSTWSGMKSLSSL

YLNNNRLSGSLPCEWSLLSALFSAHLNNNTLCGTLPSEWGRMSFVVYLYLNDNALNGTLP

ASWGTMTYLTTLNLSNNALSGTLPEEWKSMGSLSILHLAKNMLFGSIPAAWGQTWWYFFT

AALEVNRLCGCLPSEWERSGAVLSITVDPAVRAADCAVKNACDALPSTSTSTTTAERSPD

EGGADEHAEVPAGAADGGREPERAVDGHGLLRLGVRELHGGRVRGAGPVAAAADEDCAAE

PDGADGEGGDVERGCVAAGADGRHRRQPGACDDLQAAGQGCGGGGHAAVELGPPDKAGDA

VPGRQQPHWDAAL*

>Lp_000270700.1

IDRHDVVFILLRVLALFKTYPNFVDVTVPDGEDITVCGDTHGQYYDTLNIFKLNGNPSPS

NRYLFNGDFVDRGSYSVENVLTLFAYKLLYPDHVFLSRGNHEGLPMNRVYGFEGEVRAKY

SAEVFDLFSEVFNALPTGHIINKQIFVVHGGLYSRDDVTIADLQKPNRFREIPENGLICE

SLWADPQPMPGRSPSKRGVECPAFGPDVTAAFLKNNSLRLVVRSHEVKEEGYEVDHNGKC

ITVFSAPNYCDQMGNKGA

>Lp_000299600.1

LLPFLALVLLLLLCAAPAQAAEEAPEGRPVRPLHGPGYEHLDCSACLTVSRALFGRLNAT

LNENPSTYLASHRLNRANQLRRRPYRNSELLVTEVMDNFCRSYENDERTLRLHPKSKVRL

YHQQVWKDATLGVRPSLREDEVYPAGAHDPQWDNYVELRRYEIAKVYSPKDEEALKGMSM

LKATPAMCATLVEEFEDEIEELVKLAHNLSDIEYGLCGLPLPSTTDAAEVRATILPITNI

CARTEVLRAAAHRDQLRWSQYMRREAHRLEKLAAKRKASQESTEEDAAEEKAESDHAEIS

AKTTVGQDHHNGDASAE

>Lp_000339800.1

MSCLSRTHSAAVQLLVAWLITLAASHWQAAQAGADVVDRFPTIMNAFANTASGAAVRAVY

EASQLTTSTSTGTVPAVPTPLQSAHYMLCQSDIALSSTDANIAALAKTWCSRYSLGGRLI

AQTVNWSTRAAPGLQLLDQSASSYRARAWDCLTYACDTENTIVSASAATAPAALGGLGVS

APCCGTTNSTFVLYNTRGLVVQANMLPVRTLNREALQLAQAYATAQASSSASSSASAAEL

NLTVAMNYCLLRQLSPSFYGNSLEGTAQRGSGVGDGGIVVPVDVAICGEDLSDRIDTRHA

SLAITGYTAVLTTADGVDELAFPKELIEGVASWIARGAGTATGSYDAANSLSRRRGLCAW

YYNASYASAYRTGDYYDCSYAHTSLLAHLPPLLLTVRNESIVDTEDTNSSCSFAIDLTAC

AVRSEQVLRFFSTGSMMQELRLRQGAMISYRDPPIVVGMQHLRGATVAVTRAAYIAKWRN

SGYNLATGLPLLYIATPRGSAVVTHPATTSSSSWYACVQPQSCKKRQIFYPSLNRCASKE

CVGVFLYRFDPDTFTCSVRVSVVSLFAAMGFTLLVAEATVLYLRRLTEQAKEEHMRLVHA

VQRRDAGDAVE*

>Lp_000348800.1

MRRLLALQAAGVALAAARLITTESFVSSVDQVQQQLQTAHDRRTELLRYGRQLYEVEHLD

DLQTAYTPAIASKVCHLASQLRIDSTKGHPFAAVVEASISAEGEKTMDVASLARIVHSCL

VLRSPYLYEVLFTFVPFVRTKAATMDAVSTAVLINAYGRSGVHHPGLYKAMCDNGAVVLK

DTRVSLAHIANVAYAVSRVRYLHAPLMLTVRDHALRKVDEASPIIALTLLDAFTELHQFD

EDLFSAYEQRLLEHLGELQAPLMASLISCVARAGRGRPEVMETLGARVTAIADTFDAASI

AKVTNAYYQAGVLSEDVFGALAERACKMASDFRPDEIAAVLNALSAFDLFDAELFPLLAT

RLSSLYKQVGYVDVADAAVILSSFAAVQECHDELIYVCSQIFAAHPDAAMDSATRINALW

AFATLNVHNEAQTKMLEETRAKPTLLRLTTGAHLSAKDKAVLEERREFVGKVYSIDLSTA

TSAAA*

>Lp_000348900.1

MYRESILGLALAATLAEINPRLTPQQEEEVWRVFDASMEVSAAEAPLMSHVAVHVPPPSA

VAGGGMTAVSTPIPQFHISAEQGCSEASAPSTVTGQREPPPTPATANTEASADTPASTAA

SPPLPRTTEAPFDDSRIAFPVYRICNGVWTVLLKDPTVVVRDEFGKSEKMQLDYLKVRLK

DISDKPKDTRRKRAKKS*

>Lp_000361900.1

MSAPHLTALFFFLSLPSFPCLFSTAHRTSCACPSSSTRPPSPSALQSFADVAADALLLLL

VCACAVPCVVTSVLSLSVHWDLLVHLTRLLCVSEPQNPIDIAATELALQRMNLFRGIQHY

FSASSAELKEVLDTRVYIWNIPAGGTTLTADRALRVGRTTTRSIASTCAHLDQEEADHYI

IFNFSPLMMELVDGCHRGQVLDFSKQPVENFGLIMEVCFTIRKWISAGDFEGSMYSPASA

SLSTSVNSTAIITNGGRPHTHCAVLAFLEETPAVAHPNYAAMMAACYLIFSGFPTYGGSG

TLEFVEKELGVARSKYHALSQVSYINYFQLLFEIPEVPNKKRLTLTRVSLHNMSSLFQQK

LGLQLENGEGQQPKLFSDPDAWKVGDTETLELYLDVNESVFGDFVLNVFQYDVLPLPADF

DESSLQGSFSPLGTTQGSALASSIDAAFMTAATAPASSSSPYAQSASSPSFDQAGSPTIN

RFVIGNAKTKKIVQKKRLFRLAFSTIFIAQCVHRVRVQDMDYARANALLEDFYVQLHFAE

CEPVDSDASYIEQLSQRVEQSPQRQMINSRRDPRDLLGLGGGGSGRAGGAYSASRHRDSN

PQRYGGGVYYRSDNTRHGAVESMDPRWQEPSMPLTGATILYLPTSLKMTSEDERHFDDED

DVAETEPTLVRVLPPPGMTTTRPRQPTPERMYGGGVPIYPEAVEDHAAEKAEAPAQQQQL

QQRLPPPPTGLPPPPPPPVPPSEATPLKHVDPMLEDSPPPRPPPPTAGGLPPPPPPPGKL

PPPPPPAPPSGLPPPPPPPGKLPPPAPPSGAPPPPPPPLGKAPPPPPAPPSSGTPPPPPP

PPPPPAAAGAAPAPPAPAAAVKSKPKYTGPRLKTFFWKKVNNSSGIWAVSDGDEVRRAVV

DEAFMRQLFEVKAVTQASQAEAAAKKAEQERRSELRSNVFTGQRLQNIGIALKRVQVPVE

DLCAALIACDAAVLPIERRETLSAVLPTPEDVAALAIEKKAGRVVWTDVETYMYTMATTV

KDVRERLHLWTAAEELPDSIRTVSGLLASVDAAVQAITQRSGRFARMMRVILAFGNYLNR

GTPHADAAGFRLENLNQLNFVKSTDGKTTALMALVVSLLDTAEAPHASLNGDDAADPQKS

TGGAAGDGGKEREDSRSGPSPPSGGASAPDGGISDILRFTEDVSCIRAVSASPLQDMGQQ

VTQLNFTLQRMRRVVEESKDVKAWYAKRLPSTSPADAADALPGLLAKAVESDLATVGQLA

LKYQQLREDVSAMLASFGEDANGDETTVWGYVLQFSKDVQHCVEKAKASKLTKQRLMGSA

AEEGPVKMGKEQVVQGVVASPAASSAAAAKAGSVVASLGSADGLRRARLPKLVDDEDD*

>Lp_000369600.1

MRVVIVGAGLSGLSLAAFLRRLNVDCVVVEQAPFLQANYCVPYTLFANALSCFKAFGMEY

IFTNSGMRPEECFGIQNDRGSWLLRIPNKDIHLQALGGEDVIPLSTAAPANSESIVSQRL

MEAQKQEMGYVPLRCTMPASYLRDALRRHIPEIKFASRVVDLVPHDGVKGGVHAVLEDGS

TEWGDVVVGADGMHSTIRRLLYPNEHVGTSSRSLGTMQIDGFTDCPHCPPHLVQPVECWG

THRTLRVVPLHRFGENTLAFSATLHRLPQEVVDVRADMDAMEVLQVFQRLLGREYADFGT

DIATLLSRAKLAVPTEVLEVPLMPRWYNRRAVLMGEAAHGAMPSFLDQDASLCVEDAALL

AVALLDVPLPRDSGFEYAFRQFETVRRDRVERYIRQSRRARRLTAMSSTAVRNTLLRLTP

PIVAKLSQRWLSNWSYSSQQLAVDPKIKMETAFR*

>Lp_000372800.1

MQFRSIPRLAAAAMVALVCLLALSANAASGPISDSDTHYFVLLWAKKYPLNYVWSGDNLC

KWEGISCDTVKQQVTMLLPKVGLTGTIPMWGSKSGFTPANVKVISINLAKNKLTGAFPAH

YGQLTKLQELYLLDTSLSGTIPQAWNNLASLTIVDVSNTKACGGVPAWDQTSMPSLQYMH

FTNNSLMHGTVPASVMTFSKVSFNTSGCHLCGCMPAGTSPYIRVQLARDQPQLATSTCAT

SNTCSASDLTCMTPKTKPSKGSSHSEASAPVLALSSLVAVVASVVAALAV*

>Lp_000382800.1

MKRFATLLVLTAAFLAALVSADEDPGANMPGVVQLTKSNFNELVGKKQAALVEFYAPWCG

HCKSMAPEYATLGAAFQRSKNAKDLIIIGKVDTTEERDLGDKFGVTGFPTILYFPAGSMK

PVTYDSERTADAFAKFLTGKVPGLQLNVLKEMNYATELTATNFDQVAKDPTKSVLVMFYA

PWCGHCKALKPKYNQVAKIYENDADVVIARIDADNAKNKAIASQYNVHGFPTIFFFPKGE

KTEPEEYKSGRDVEDFLKFVNERAGTHRLANGDLSWDYGVVEALSKAAASVARAEGEEAK

ATAVEAVKAAADKLPESTSTTYYVKVAERIAKKGSDYVSTELARLQRTLDGSMTGPRRDN

MLIRVNILTAISKEL*

>Lp_000395200.1

MRRFASCFVVYAAVPASRAVHSVAVKQNAVAATTADKLGSLVSKAAIAAANNVPASILEK

ACNPRLVSFTEMDLARTRFAPLIASVTADIDFNHVEHQVANKFNADPSKLKPVPGALKKL

RETQWKDMASVITFYEETMYPIRMKHNEYKDYELTSFHIKDVLKRGLTAFKQDFLDQQKE

EMTKVKAHMKVCDDFVNAAVKDAFDTAICNDIANILRIAGETHAHAHRMALKILDDMNMM

KIPYNAATVEILQATAFNDGPFDNSPLLFEVIEYPERGEITVGSRSLESISDEMLKLISN

RHQTPLDDGVLLRSQETHPNLQRSPE*

>Lp_000403800.1

MLTFDRKYLIGAYLVLLCLLYCPDVRAENSTHITDEAQSNTLKFLQALAGSTTELQSTWT

GTSYCGWTGVTCSGATAAVDFADVTLTSPMKLPELSTDVDATQVAVTSVKIYNKDSMVTG

TLPSSWGGLTQLQVVDLHGNALSGSLPAAWSGMTAVTTVYLDSNQLSGPLPATWSELAAV

ETLMAAYNTLTGTLPSEWSALPKLSTIDLSNNALSGTLPAAWSGMSAVSYTHL

>Lp_000417300.1

MKRFSALVPSAVLSCAAFPCVQVPRRSAAAVADANERVHSEQQEDEKAAGKEGKEGKGDT

KTTALGAAADENAPKRRNIVVLSNTTKAVYVHPINHIASWYWFITDFHKWFMGFVFLFVG

TQVIARYRVAKLQTATQAQLGENLLDQRTRDLLSDIEVLRKKDPIRLEHEANMYHEQFWK

RRAVAVAESRSQVRNLEIQRGKMQGEARGTDMTEWLGAKAKDDEEREVARRTQDYIQGFH

QHLKSKRLI*

>Lp_000442900.1

MRRFLVVACALLTLAVVGEQEVKHISLKRAMNAAVAHSLGMPLVPTRRFKGPQGEVVIHD

YLGVQFYGAISLGTPAQEFQVIYDTGSSNLWIPAHNCSLSCLLKKRYQPSSSSTNKPDGR

EFKIMYGSGPVSGHMIADTVAIGNFSGPQGFAGITDASGLGLAYALSKWDGICGMAWPSI

SVNGVEPPMFSIAKANPGFANKFAFYLPQKDSDEGDLVLGGYDSRHVDGELVRVDLTTKT

YWTVDMTGASIGGQNISAAVTVIVDSGTSMMTVPKALHSTIMTMLNAKAVINGQYSVECS

AVPTLPAIVLTIGGKEWVLRGKDYIIGDDSTCIVGIMGLDLPGPIGPAWILGDVFMKRVY

TVFDADDASLSFAYAK*

>Lp_000444300.1

MLTLLLFILAALCVFCRACKRGYAKNTVGFLHASAGAGGGGERVLWVALEGLQRADEAKN

VQRQYVVFTQEYKSTDPTLTETSDAYLLRLVESQFSVHLSRPVTFVYLRPSFTKWLNGDL

YPHLTLLLQTFYGGAALFLEAAVVNSITPTVIETVGVPFVYPLLRLLAGCTVISYTHYPI

ISSAMAQRVRSGESGYTNANFIARSPVLRVGKLVYYKVLFLLYRCMGWCPHVVFTNSSWT

QNHVQAAYWPRQCIRLFPPCDTKSFAAASKPASERRHRIVSVGQFRPEKNHMLQLTAFHQ

ALPRLPADAKLIMVGGARNAEDEQRVAALRVRSQQLGIEGRVELRVNADVHDVRAELGSC

VVGLHTMLDEHFGIVLLEYMAAGCIPLANRSGGVQLDIVNSAELGFLASTQDEYADAMVE

IFDMHQQHPDEFERFQRLESDHVKSFDDATFREQFVKLLSAYLYAA*

>Lp_000449900.1

MLRRSVLLFTTTAAAGVAAVCAPQRAESVPPMPVHLKPSSSPHRKPSFYFPRMREEGHPD

YVFQPARLGAALPLFNITSSSCEYYALRLPLLKAGFRRVPAGRFDVASNLIWGRSMPFRE

LLHSASSSSLDFPPVHTPEEAAYLNKLTMVNQHQRFNHYPLSHANLGCKRGMATNIRNAQ

RQAEAAATTAGEREAARMRYGFVPRTWFYPQEKNSLVAAMKAAPTSKHFIWKPARGSCGR

GILISGGGARNAGSWEAAMREIDAKAACKESGRLFRSYVVQEYIDNPFLVEGRKMDLRLY

VAVTSYNPLTVYWHEEGLVRLGAGSYADAAVSPSSHIDIGPAETEEGDSAVTMCNTVASA

AAAAAAAAAAGTTSASTSHAGLLGMHIHDRFRHLTNYSVGRKYVAIQEAIAAAAAAAAAA

TPLSKSDSSRTDVPSAEVAAAPELKWSLQRLWDYIDAQRATSVTQPMRRPSVQVRETIAQ

LITRTLMAARPVIDSAVSRVPMPGHYFELYGFDVMLDAQLNPHLIEVNTLPSLESSSSFD

YATKTNVVADLLNVAMIEPFERAVKPGSSLWNATTLREELQQPLGPTELALVNPSAAESQ

TKGESYGKDVALTREDVQLRLKDELVYARGFQRIFPAKPLPGFSYAVMGESMFSVSPFVP

LSHAERPLDSLGSRPTYLDDVRFYDKTHVLTPRDIWALESS*

>Lp_000460700.1

MSSVKKALVLLVLAAFALSCVKADLVELNPENFKAIVNDPKKNVFVMFYAPWCGHCNHMK

PTWQELADKYSVDKDDTVIARVDASAHRGIAKDYDVNGFPTLKLFTKSNKVGKQYSGPRE

LDAFEAFLKANVE*

>Lp_000473300.1 hypothetical protein, conserved

MRRVVCLSSAAAVSALSLAATTTTTVHARSIYTTWGSIPCESWACKESWLKRITSQCEYR

SFEFWHVPEVAVPEGLKLNAVERYLLSSLENDTDRLLHISWCDDFDSYWHDRVSSLEMLY

KTIYTDDYPLYRYVFGNCSHKTAEKEFMLRKLNYLRNILFWAGRTERCYTSIVKARYYMQ

RCVWNALERERYLCACVEAVDSFGKKVPEELHEKAMGELQVALVNMRHWVWDCPNAKRTF

TRRLA*

>Lp_000486100.1

MAWLNHPLRRMAVAAALLVLCVVDASLVGARTASDYTAAQQASTLQFLQGFVTANPSLSA

VWTGTDFCSWSYVSCSSYSPSLDFSPRTSPYSGTLVLPELGDDVNGSAVIFTDIKVRSMG

VRVTGTLPASWGRLTALETLYLDGNNLTGTLPSAWGGMEKMSYLHLNDNTLSGTLPVSWA

SMTSLSYLYLNNNTLSGTIPGEWAGMTGLSSVYLKDNHFHGCVPVPGRCWSPSLSVYAAL

KPEPGPLSNRCTSGSGSGDSLEGMAMAEEGTSTLKFLRCFAALNPSLASIWTANYYCDWA

YVSCSSYSPSLDFADYRPAYSGTLVLPELGDDVNGSAVIFTDIKVRSMGGRVTGTLPASW

GRLTALETLYLDGNSFIGMLPSEWGGMTKLDTLYLEKNALTGTLPSAWSGMEKLSYLDLN

NNTLSGTLPVSWAVLASLTYLHLNNNQLSGAVPEEWTGMPGLSSVYLKDNHFCGCLPAKW

RSSYSPSVSADEALKSDRCTVFNRCNSGSESGDSLEGMTSEEVSTLKFLRGFIALNPSLA

SIWTGNYYCDWAYVSCSSYSPSLDFADYRNSAYSGTLVLPELGDDVNGSAVIFTDIKVRS

MGGRVTGTLPASWGRLTALETLYLDGNSLTGTLPSEWSSMRYLRYLDLSANGLTGTITLS

*

>Lp_000488000.1

MKGKRGAQVASTATLLNLLLLCDVFSTTQYHLVLTSASAAAAASEKTAFSPAAFTNLVQM

TEAVLAPAWGTGGFAYVLDGFPVDQKSSLRPVCFHTRTSPSLVPSPFTFAAFDVQFSPLY

LVAADNATAQAIGQLDLQHGCYVALHTNPAACTAAWASQARQCAFGSDDALRYNWTAASL

AAPTASLAADLAKATSLPASGLRAPELFAAFVNDYLPQRAWLRTAVPRTFDALYTAAQTA

EVLRSSMAALADNNEMVDMKTAVVSEWNRYLWSVYGPQLRRKWDLLYRYAVATLTRGGGV

APPIITSSDPAHVAESSITGVVGPVNATMCPLVDAWLQLRYSPDDVNVSRLTQLMEAAFD

ALADTFAGTRRNASSSDNTSPAETDALLLRLVRSTYCVTTTNLFAITALLDRPFLQAVSS

AWIEKADHADSWAWQTMSLASVTEVVSDASAFQSFTSMRVSAMSAVLATAAEQSFLSCLA

SFLDPTTPLTSATLVSPSDCVRQFLEMRDASASVTANFTALFELAKLSSVSPDVGVPGMT

RYESTVALPSVAGSVANESLQWLMVQESARSLTAQTAVFTPATVSSSITPVDPCTKATQR

IAWTDAWDWVYECDAGTYVLERNATCATCPSPCAAATASFAFPPSTPQSAPSLSKAAVPV

YCPGDGLLHGCPSQPAGAVYAGSTSGGHSWPFAICRYACASTYMAPLNFSCLSVSGFFYN

TSATGNADVNNGGGLSSCAGPASLFPSSAFPVRTQTRLYAFVGSGATNAPLTCPFTLLHR

ATSSAVSGTALATVGAIPPPPFFRDSMMPAGALASRSGTTWEVEVQLNATEMYALHAALR

ASSKERQDSNGLSGTIHANADTDTVAAHELIGVHKTATLAVSGSASSAGDAERVMTWYLV

STLHHGEDRSSSENNATADPPTASTTVHLQFLLNLTVASLSSSSSSTSAETANMAATAAA

SLTPAPSSLVVLSSSWLWSPWSNNAAAASLRSTSTYRAVLDRQTNTVSFLVDGQTPLSAA

PVTVDWPLWAPAETSVDTDGVTAVAPTPAGASDLSFFLSVGGWVAGYARAQQWVLVPSLP

GSPAASDNVVLPLYYDYLPGMVTRVSTVASVVNHLHNTLQAAETMWRGEMAAASQSQSVA

QLTSLTDLAAYVAATPRAAANVSAALNALALRTTKGLDGVCRPGYGILATTPSHALCEVC

MTGSYTVMSAAAGSNVCACITQRVRGDTVAGAAANQQRDACLARSTGPAVPRDVFAAART

LYVDGDVVEAANATAARIPLGWYAAAASGDTVATTTSLVSSVTCTQDLTYPIAAAGSSTD

TLLNTSVALLYTTGLGGETCETRAAARAGTVPSSSTSTDGTTTAIMPAGTSRPTYTASHS

YGVRLQTPALTPTSISLKNNTALFTDQTVLTVGVRDYALLSFYAVHSLLQMPQLTEYHLS

FLAYVLRSVTVAATLECRADSTNSQKRSTTHSLSWAGASALDTAVLWLNLTEAVTHDCIL

TIRVTAAAAAARDSAVPFSSWPYSYLRVPSRTPPSSFFESSAPLTFVVRAPSSAFAVASP

KHNVLGEVSSFVMAICIMLVSLALVPFRAIGGYAPTLDLLPWRRYERELAKTLSAQDEG*

>Lp_000507300.1

MAWLNHPLRRMAAAAALLVLCVVDASLVGARTASDYTAAPQARPLRFLQGFVTANPSLSA

VWTRTHFCSWSYVTCGSYSPDKLDFSPSSSPYSGTLVLPELGDDVDGSAVIFTDIKVRSM

GGRVTGTLPASWGRLTPLETLYLDGNGLTGTLPSAWGGMAKLSSLYLNDNTLSGTLPVSW

ASMTSLSSLYLNNNTLSGTIPGEWAGMTGLSSVYLKDNHFCGCLPEKWRSSYSPSVSADE

ALKSDRCAVLNRCTSGSESGDSLEGMTSEEVSTLKFLRGFIALNPSLASIWTGNYYCDWA

YVSCSSYSPSLDFADYRYPAYSGTLVLPELGDDVNGSAVIFTDIRVRSMGGRVTGTLPAS

WGRLTRLETLYLDGNNLTGTLPSSWGSLAKLDTLYLEKNALTATLPSEWGFMAKLNSLYL

NNNALGGTLPVSWASMTSLSYLY

>Lp_000521100.1

MHRGSRRVVVVSRAILMVACLLLLSSAHELSQDGNSHDLVARFSFGSGGVVQSALVAEEP

AGPREAYASSFVPAEASARKVHSVETVEDSTAASEVENAGAAATTMTVEPEASSSSDTTA

GHFDLSGLSDEELKTLLSDKKKGQVNFKRYKDRNDLVEAIREVERKEVAQQAFRSKVAAA

ARWYAETQEELKWQRATARRAKASRIKRAGDIPASLRSDGSADLPPTHELRVLYSEANGY

GRHFTALVKDLESLEAVRLPNRDAFRFLGVPYPITKKSALLGQTLQVCFFGVLALAIVPD

LVPFIPEAGRNMVRTRRSLIISTAFMLNMLGRTVLQGNAFEVYLDDELIYSTLKLGGRLP

TSEMVSNMLLERTLLKDYASAMSGQAAAAAAAAGALL*

>Lp_000525700.1

MAWLNHPLRRMAVAAALLVLCVVDASLVGARTASDYTAAQQASTLQFLQGFVTANPSLSA

VWTGTDFCSWSYVTCSLYSPDKLDFSPYSSPYSGTLVLPELGDDVNGSAVIFPAIEGRSM

AGRVTGTLPASWGRLTALETLHLDGNGLTGTLPSAWGGMTKLYTLHLEKNALTGSLPSEW

GGMEKMSYLHLNDNTLSGTLPVSWASMTSLSSLYLNNNTLSGTVPEEWTGMTGLSSVYLK

DNHICGCLPEKWRSSYSPWVSADDALKSDRCAVLNRCTSGSESGDSPEGMADEEVSTLKF

LRSFVTLNPSLASIWTANYYCDWAYVSCSSYSPSLDFADYRYPAYSGTLVLPELGDDVNG

SAVIFTDIKARLMSGRVTGTLPASWGRLTALETLYLDGNNLTGTLPSAWGGMTKLGTLYL

EKNALTGSLPSEWGSVAKLSSLYLNNNQLSGTLPVSWASMTSLSYLY

>Lp_000527900.1

SLNRALVIVVLLLSLPLFLDVTRLPDSADAQEYVFTALAPCNGLRMCNLCGYVLEMVDLV

QRALANPFTDELTLSAAEWDPHSLQSYQRCLREPIQRGCDTLIHKVKTQRIEFAKLILEQ

ILEDVAFQHKWGLLGEDYIVSSAVRLTNYIFEAQERSLGGRRDPYKEDSAARSIVYQHST

LSLSGDPAVESLVTRVWFLRRDVQQLHKDFCYAVCEAKLGFLGRLRLALVRFYIHHSIKP

RLLLLREYHRGTLIVVELMMIGLAVCVERMMGPAPPWNGAAGGAAGAADGRRLPAGAEGN

GNIVGGRGGDGASHGPSAAAGGGKSHYRQGRRR

>Lp_330007000.1

MLRGLVKNEQLTAEQEQHIDQLQSEIDEIWNRRGNYQIDPDAWEDMPFFMEHISEEDIAK

NASCAALASIVYDEVPPDEIAENRKEHGNRALNLALNPSQERRENLARAACSSYTEALQA

KGKDKKLTSTIYANRSLAQYIIGNYGHALEDAQRSIVLNPDYRKAYYRAAKSALALKKYD

IGLQLLDKGRCVTDPPMDAATTAEFAELEAQCRSCKKKKSAEDAKLSRTARVQAAKISNV

ARAITSAGIKISAKAEVTSEQMGVYGNPQPYFDTDGLLHVPLLFMYDEYQQTDIMHDVAC

DVCAAELLDELMPFPWDDRGRYQKFDDIVVFYKIDDGVKDPEYYELHQGWPLMEVFRCET

YAMPQLMPVLHVVCKSSELVERLKICRAE*

>Lp_350006100.1

MLVFDLVAAGIIAILILFLAVYVVCRYSAEEEDGDAWLPRALVVLSLSVACYMVLLLPLE

VAQKHEKPTLSFVEAWEGMLACAYFFLFVGGPYAFVFYESWSPHQSSVWAQVRPALGTVA

AVNGVFFLSFGALWLWGNRIDTKNGTLTHVLPFEYFVATASSFGWVVFFIFAGVGLAAVP

LHGALTFLRRPRPITKSEYELAKVKLSMEVQHLLRRGRQLDAEVGGGRPNQRQRQRMLLF

KREVRELERQSVANETAFHLSGLTILRYYVAAALSVVNGVVTLLWVLHILLWNVLQVFPL

LDSLLVCLDHLLPMLAILVYAYCALYMMWCTLVGCSTVSGHLLVSVMYPLRVRGTMLNAL

LFNALLVLCASFAVLQLCAVSFSTYAASTTMHHIFTVTIARMYGVQYVAQYLQYALLSVF

ALAIPWLLFCPQCMSGEASGEDEVDSLV*

>Lp_350006200.1

MLGLDNAGKTTCVKKFCGKDTSSISPTLGFQITAFTFRDCTLNVWDVGGQQSLRSYWRNY

FESTDGLIWVVDSNDVERLQMCKEELHSLLQEERLAGASLLIFLNKIDIPTAQSPQEIAR

LLDVDTIRQGKRHVHLCACSARTGEGLLEGMSWMVDDVSERIYFSM*

>Lp_350016700.1

MRPCRLLATSWKTKVDPKNPHRIVAIANKTPCFTELQAGKKYYWCSCGLSKKQPFCDGSH

RAYNEEHKTELQPKEFTVDKTKKYLICRCKHTDSAPFCDLSHVSVIFRTAVGIEKLPQEQ

PAKPMDKHEN*

>Lp_090010100.1

DAADVLLVLVLCMAVALCEQRKQQQHQLASLSLRGAATATSSSASRQNRSGAGVVSSSQD

TSAHSQGSSVFCEKGGSQTPPTRHHTSSGSVYTVIRDYYNVKSEAAASVVTAPREQNQQE

GLRARLQRILPVLEARLPMTELWREPSLLAALDVMARGVWHAQARLHGSTEPHKVGSSGR

CPLCYAAALNNDLAAEAITQLSHYYVIYHGEGDDEQRVAGEEGRSDVRNALESQADTARQ

QKEKHDDTSCLQMQARGWVAACRIAIWNGSGTTLACLMETRTAPWLIEAPLIMPNATHIL

FSWRREAADVLPRLWTCWWRAERALAETEHEKQSTASGDGNASPASVGVVLVSRGAGQRL

HSCLLFLETLVPSSTTSIISSPQCRSTDSGHNTNCGSRQRSVDSTQNAGADVAAAAAAVR

AGGLEVLSWLLRLPLQPQNSLRTAAPSHLDLSLDAQQRQGPAALDMASDVAAVTGTLFHW

LCAHNDLVLLRLFILQLSNSCTSWQSPSNLLFEQRRFHCCSAGKSEGGNCVCAARPARGA

ACAAAAAAAAAAAPGRPQQSRHNANAPSPTSAGTLARIPVPDPTAATRFSCHNVRLTALL

SLRDAHGRTALDVACQYGHVQCVRLLLQNGLSPNSLALRTWRTDAAAALPLPFLRLLYDP

AVLDGSVSVGGHQPTSNAPRVSLADQLRSTRCPAAAARTLAQLAHRLWVEKAQQQRCPTM

QAAEDASRSEAASTEATAITLEKPRPGDGCSYEAWCTAYLLCQAAQDRVLRPALAALAYD

AHEPLARLVVVAVLTRKLMEWLTTTPPAMRSGENSSFPAATGVMSLRDLHLHHTVALRLY

TQLTCPFGRRLLHDAVAIRSAICAAVELREERAAATLLSSTAPPSISTPESTAAPGEEGG

AIGIKKKEDKAKHGIAERSDDGIKADKHGRPGPDGLTLEHQQQQQQLPLSEAAKARRELQ

ALYANVIDGVQHLDALLVAEQRRVTARAAAQEEQQRQCDWPQQRRRQSQDGEAALTGAAT

VLRGRRVQRSGGGSGDDDNSESDVPSYVLVVIPSILALQRARWTNNNNDNDVSSRSNSIN

AASTSNLSSCGCAVVEPTLWSPVWTPRQVWLQLPRGLHLDELLRSHGGCGYGVVSHDMPL

VAPVQRGDLITFHHIKGSGSAAPQFVADAIFSEPRMLPYLTVTVKHPQQEMPYASPPTSL

VLPFTHGPQSEGASTTLFSASPSPRSAAATATQRSQQPRCATSVTLHACCLFSLYLEGTA

GTSQPRDDEVPQPHPLDWLQARPSVSSGSESKGGKGVAADADVLAVLRTQPKFLSLWLRL

VWTGVQPCWLASVREPTNAGGAQAERSSA

>Lp_100005500.1

MTVLSIASRVAVVAAVVVLAVTVASAQSAGQTCTASTKNPLASLIYTDPAVEGCVRARCI

ATLGSVPSLRTGGYCNGDPTSESAVSCTTLFNAYNQYYSCVIRALEGSTESALAEVVQEV

NMFLRTPGYPYHLSPLGCYACNHFRESVFPNLGSSCTNWTCDSVSSQSADSDWLGQPGKG

YNHQLCGSGCIAAMMLTIPFALVSASMFIACGCCWPSPSLKTTYAEAQKEEEAEKHRVRS

DDEDDEEDTNHEPADGDNRDVAHEPVVADNNINHPEEEARPNEYVNLG*

>Lp_100007700.1

MLSRRSIHSAAVVFAGCSAVATAARSFSTGSSQVWIVLSGDAAHPCANGIPWTLRVPLDV

SFHTSAVSTPSKTAFAHTGHGVLRRSRTLLAPLHSPSPPPAGSSTATASKAEETKELRSP

YDVTIDEALLGSSNLHSSSTTTSVRDLVRLLVDPDAGTAVEAVSLRDFAAMAPKRQLRAV

NNDHDCLDVLLNSERYQSEKDARTLRLHGITTVGQWREQPDRVPLSAYGRGVLEATYHRH

ARGQQQSQKDLVVAWLTRTIQRAVAEELMAPRQLPAGSGAPVYAQPSLVSATATVAAAAD

VAQKAAGDIAESTEEAAVAYTTTFGMPEEQAQTAKPSKATEAAMAEEEEEVVEEEKEAAP

RRRGPRQRPAVPQVSKAQAKAAAAAEEQVDEEAVEEVLEEVSAEQGEEVDAKMAEAEAIL

KKDGPLPPVSAAAAATTTATAVSASKAGAAAVKGSEGATAPSAEEEEEEEVEEVVEEVED

EEAEAEAAATPKATPTASATPPAAVNAAAPLPASAAVSTSSGQQKEKLARLAELMLKQFN

TADGYLHPTNKAARAEQEQRGDILVLDEDTAQVNPLTMFSAYHAATVTEADPIAVRALKT

IWTSYNKHQMALEEASTEAAAEFYKHESVNALLYGSVLMRALAREEPVMVLEGTSLLPPY

GFPLVSSDTRPFSAAATSSSSPSVTDMTPGKVLTAMQAYKVFFSDKEKRYSPFRPAIDRN

ANCSVVRLSGTTLHLYYTSTNDHVVEEEVRMPFAEVMLGMLHLLRQKVLPRVNVVHYHLI

ATRVADTAADADFLLRSTATLDLTLPFTKEEHTRLKALAAPLGMSDEVEEFGCIDDVVME

IEDNSGASVVHSLLHNREDLEATLTATLAQLLPEDVERGAESADGEEDNEDMVEVEEEEE

ELAAPRGRAHAAKKPPHHPAPPVAPTTRRSAAAAAKDDVDDEEAKLEEAIYTQWQKQKEQ

QQQRGSAATRVSTTARTIAAPQAVADEDTEEVEEEIVEEEEFEEEEEEQAGEAASSPSPA

VGATTSGTTARMTRSAAYSHHTASASTAAKSAAAASPARRTAAPPPPSSPADEEEEEEME

VVTAPVLPRRPAGHRAVPVEVDDEEVEEEVVAPRSPPSPPPPQQQQQQHLQHHRAARHAP

SATHTAASPAPVRARAASSPPVTASAAPAEDAEVEEEEEEEAPPLPQVRTRRSAARRHAA

RSRVPTSLEEEEAHGAAFHGDDSAAGNDVDETVANEDQWFKKRPKRRHF*

>Lp_100008700.1

MMCGFFCWCCCSCCSLSKLFASSVDQECALVPHCLMACFLPCITSICVRTNLRNRLGVKG

NMMGDCICVWCCGCCSQCQELRSVPTVEWNLLEPAWKTPAASAPECIFIK*

>Lp_110011800.1

MIGEILLALLAGLAVYGWYFCKSFNTVRSTDPPVVPCHTPFIGHIVAFGKDPLNFMLNAK

KKYGGVFTMNICGNRITIVGDVRQHGKFFAPRNEILSPREVYSFMVPVFGEGVAYAAPYP

RMREQINFLAEELTVAKFQNFAPSIQHEVRKFMEANWNKDEGEINILEDCSAMIINTACQ

CLFGEDLRRRLNARQFAQLLAKMEGCLIPAAVFLPWILKLPLPQSYRCRDARAELQAILS

EIIIAREKEEAQKDSNTSDLLAGLLSAVYRDGTRMSQHEVCGMIVAAMFAGQHTSTITAS

WSLLHLMDPKNKKYLEKLHAEIDEFPAQLSYDNVMDEMPFAEQCARESIRRDPPLMMLMR

KVLKPVQVGKYVVPEGDIIACSPLLSHQDEEAFPNPRDWNPERNMKRIDGAFCGFGGGVH

KCIGEKFGLLQVKTVLATVLRDYDFELLGPLPEPNYHTMVVGPTLEQCRVKYIRKKKTAA

*

>Lp_110012000.1

MRLKSTNLCCAFVWSPAVLGHPPLLATASYSGAMDENFSNDAFLEIRLVDVKVTDETDLP

VVGRVRLPDRAYRVDWSPYTGDQGIIGVACGNGCVYIFSAKEVLAEGPNGGHQEASERPR

GLLWMVKEHAGHAVRGFHFNPAKPHFFATGADDGVWQVWTLQDGATGGVCAPTKISVIAN

VPNSGAIVHLQWHPKYAHIFATATASGVVNVWNLKMATRVTALNVSKASHGATITAIAWN

PTAATQLVVGLDDAHPVLQMWDLRTGVVPLREMAGHTGGITGLAWSEQEASMVASCGGDG

RTMWWDPNTGEKLGELQKVEQYLVDVQWCRALPAVLATSSFDPMLCVSTAQDMSNAGGDK

LSAVPKWLIKPCCASVNMSLTVASLVPGTNHDLMLNNLHNLPMSPKTTEDQKVMQQMLKF

PAGTPERTQWLRDNHHDLLAAFASAQSSRQPILDFLNAESDQPNAAARAGHGRNDDDDDD

PFAAISHENQKSYEDRSSELVASGKIDEAVDLCMDEHCFDDAFAIAFLSGGELIRKVHQR

YVAYIAATDPKKRHVLFAGAIASGDFRALIQADVPWKEVLSAIVAYVTGDGFAESCNMLG

DVLREQQNYEGAYHCYVCARNVDAVVDLWRLENRPSREVVQDTILLEETTQRAASGAYLA

HCMCDYGMKLLTDGHPEAALQYLQRASRIGDHTAAVLVDRMKYLYDVSQEKPTVPYEPAP

VSDEQSASCQAFLAAAEERRLRDLHEQQQQQQEQQRMSSPHPSAMPQPQPQQPQPQPQPR

PQPQPQPQPQPQPQSSPMMVGGGAGGYMARPAAPPSNVMGGFAPGAPMPPQNYLRPSSER

PAPYTASAAPGLPPPPPPPQPQQPLSSNNPSARPPLMPNTASSLLPGRSSSSMPPPPPTS

NGSVAYNVNHHNPGTVTGMPPPPPSAGSGAGVAAGGVSALTPNRMSSSNASSALGQPKPR

LMPHPVSSFSSMRGPNSAGAVPAGPPPPPPPSSASSLGSRAASQPASLYDVTAQPPPPQP

LQPQQPQQTPPPRPGMPSAAGAAPQAPPPQPTASLASVNPAQLPAPLHGQVVQRLQQLVP

YIADPRRRAAVEQAAMELVRQMQLNLLPEDLLKLLLYFCNSAGTPAAAQVWAQITDRYGQ

GVQPYANLRYL*

>Lp_130008300.1

MFGGCLELCLTFIGVQPARCGCSSADLSEFICKDVLPYQGWAVNTTLRDTIVRLLLTSKA

EDFSVFASVRDDKTVQEAAVAVTGASVGGGHSARGKKAGIATTTTTVASCTGKSGVPRGS

STCTPLTAAEAVQQLDAGRVAWARPMIMVSSSSSTSAALLKGEDAPSDAGQDSGTVVLHF

MPSRELRERVLGFPIRNEKQTMVADFIVENAMQGWEKLPNTADFKARYKSYLVRSGLKWL

RRTHKITQLYLYRKATHEVLYRFFPHFALPLTECEKAEQPLCMTEAGATSTHAFYTSAGD

DATKPEGLEANEDGEAQDDSVHHPTSLAQRRLCRRYHPLDHYFARQRHGDVAFPPVVFDL

QLTRRLIQQSPERRLLMQDAVHAIFAAHHLYDTRWEDFNRFERMHLRRLIDCAGLRIVVA

TLTLRGRRRELRLVVPAEDVLSEAVTGSSRLPSSSQKRPRSGSESSDDDGDNDNNNDHSD

GEENTNDEDGDGDGDDDDDDDGSNSEREAPPSRRVKREEEELPQGRATTRHKQLEAAAVL

RVDPSQTVELQAAHEAERRPLALTAFFPRMRFEMTEKQRNKLLGAFMEKYIKLRGTLERQ

YLLMRYTKVFTLVYAPRLPTHGSVAQGGVKQERLGDEADEASGKWPMLQQSVHAAQDSVT

SGAPSPATATFHFSANDARLPKGVSAYAVNTVLDLLLRATPHHAASLPRLSQVIDITTLQ

RRVLPLLRQLKYVYTTGFTQRGKKRVGMVVLCTPQEGPFSTSTFVKKEEGEEKGEDGGPR

LSDEAKRAVLDAEAVEAARLQRTARPPPPPSGKAIPSALVLVAVPAMAEGVAHSTAAAGL

MRAHPKVTRVVSQVMAVRNGYARSTLQRVSRLHMELWAQHCRYADAVAGEGVRVCEMYDR

MTLSTYCIVVGLPQGDIAGALELASNTHNASSSGNGSGAGGGVSSSFPLPSASPSSYLWS

TPIRALPPSVYGWCVQQGLQMLCMCLTELQDRQLVRSTDNFQHLLVPELREDVRYALCPS

ASVDGYTHTFAVSPPCASSSLYACMRYWLPYWNTVRCVQHSVEAQLLSAEPNPSVPQVVA

LSRVLRRDAGAVAAQLYQNCGVPRELHRSHHLRDMQRAVGRASQGRLTLATPHPALARQH

LKSRSQHSRPAIIADGPSSAALFGAARSKANGAVARSRGPRLHRAARTEINGSTVGDALA

YAFHNFSPLKRVEEVVQVVLRGRTSYLTQHPLFCVPPLLPAVLRRGADVAATSAASSSPA

TAFGGMENVLRTEFNTSHSGQGGGSYNRMLGVSMLHMGALAETLRALRQGHGARRVAHAE

GRPAFASSTEALKAEVESVYAPDAGLSASDVNMSDLTSFFSSAAPAPAPALSASVVAQAV

SAEGAAQQVRLSAEFEVVTDVLRMILLSDQAHYRASIARALLSALNAEALVHRARAFLQK

MPVFSRSRRHNFRVPLLAFVQSPYVLAPAALRGGPQPCANIAVLHHWTHAMEDMGSALSS

GRRRLHLHGQDAAIHALLSSSAPSNESYMNVFPAPVHMVHQPTMELAQLAMAPSEVRDYM

TRLPLPHFTWPSAAVECERQRWSARERHSAADTRLRRHHRKRARSARSDASTIPAALIAE

EEDASDSDVEDEWAVRRVEAEATRQLYPARSLRHLGDPTYLQDLPVDGPPTPSEQQAVER

SRLAMAEGRCAALSSDAAAAAGAPLGYPIAYPSIFHHVDGSFHQYMWQLVVHVVFRFLLR

VPGIGHADLESRLMASGVLSQRAVRAVLQFLIDRDLLRCQQEEVSDDRCVVGGGVMDAPP

RVTGPFQRRTAAETEQRGAGAAREGCVTAGMEGRFNRCCYTAIAPLEAVQFAVSAATW*

>Lp_130017000.1

MLYFIAAVLCAISTLLLLNRALARVRTSPARTQYDYDVIVVGGSIAGPVVSKALSDQGRK

VLLIERTLFTKPDRIVGELLQPGGLNALREVGMQECAESVGMPCHGYVVVDQKGEQVDLP

YRKGSYGVSFHFGEFVQKLRAFVYDNCRENVTMVEGTVNAVLTEGLSFSERAYGVEYTML

EKYKVPANPFREDPPKADPNAPTVRKIATAPLVVMCDGGMSKWKSRYQHYTPASDYHSNF

VGLILKSVRLPKEQRGTVFFGKTGPILSYRLDDNELRMLVDYNKPTLPSLEKQSEWLVKD

VAPCLPAHMQEEFIRVARDTKELRSMPVARYPPGFPCIRSYVGIGDHANQRHPLTGGGMT

CCFRDAIRLANNLKAIENLRCADAEQMAVIEDKVQSAILSYCRFRYAHSCCINLLSWALY

SVFSTPALRDACFDYFVCGGDCVTGPMDLLAGLDPNVGSLVFHYYMVMLYGVCHVMAKTG

AYSQGGKQLSGSEKLTNVISFFADWHRIKLAVYLLFKSTMIALPLAKNEFYSMWRFVDPT

SPIAIISKRIKMMIYLSHLNGRQRKPVGL*

>Lp_140007500.1

MLFKFLLGGVVGSVCSAYHRSSTNVSLALDPPTLDIIMDPTKDVIPIRAVIIGGGYAGSK

MAYQLDSMFDVTHIDEKNFYELTNDVIPIITNPWKEDVNPEACRRMMVLHRYYLKRSNVV

TGTVSGVDDKQVYLRDGRTVPYDLLFVATGERKPFPFQTRERTISGRVQELKRFNEFMQT

CKKVAVVGGGPVGTSLAHDLASTRPDLEVHLYHQRAELLPQLPGVCRRHAQAKLTSNPNL

HLHLSTRVTAVDGFVTDEATGTRRIAPPMSSTTAESALVLDHAEQPWWQSWLGAVSAPSG

VVPDQFSVRYDSLQSELRPQQSILQQVYYGKHDEVQACGEVTGHGAEDNFDYVFAVTGDT

PRPVQWDEPKPVHPNILREHEMKDGHYRVSTLMQLFNRPNIFALGRCANFPVIRGYGASD

IEARTLFRELNSIVNNPTTVFLHSRDGIQLSHMEIPRMHVRLGVDDAVGCTPWSGGMTGV

SSVHEFMQDRNYLMKEFQKPIFYKQQDQTKIKQRVSQWMAEEITDVVDFSHC*

>Lp_160011700.1

MFLFFFFLLEELCRLVWSSAPPSSSRLFPSPVCPPCVASLYTLASLRCSFFRARTQFSLP

RESHYSHCAKLGCISSKSTQIGERECKTAAERKAAWEGIRERLPRRKTPEDKERRIELFK

KFDQNNTNRLTLEEVYEGCVSILHLDEFTTRLRDIVKRAFNKAKEMGTKDRGAGSDKFVE

FLEFRLMLCYIYDYFELTVMFDEIDTSGNMLIDAKEFKKAVPRIQEWGVKIEDPDAVFKE

IDSNGSGQMTFDEFAAWAAAHKLDADGDPDNMAA*

>Lp_240013500.1

MRTTSLTCAIAAFGCTAVCRRGASSSGSSSGSGGGYRFFEDMYPKHDPAPRYAYYEDQMQ

SASVKSLEPKKQMSESEKLHHLHDWKKDKEGFTPPPEFGWPEEWGPEPGEEGHSEWYKKN

RMYMSYEEKSRYDLRAGVPVEKNSMRHQEMPYQKRMKAAYVNLEEKKPQYFEPRQRKYWA

EYSYEDQKDTMAKARDDYLTEWLEKPGVTKENVAKKIGDYNGTAKNQNIKSVQRRPEWEL

APRPEYDDD*

>Lp_240016900.1

VLSLAALGLAHVLYIVLSNPDSRCTAQFCPAPPLCHQQTRTLHQRVVCAVGGSPLHPYSN

EFHRYVLTRLSHFVEESIKGQLMAPAVLEFVRYKLHHPYEPMVLHFAGDNGVGKTRLAEL

VSLAYGQRCGDELCSTGDSTLVLSGTGYDGLTTTEFRTLIVRLVTQHAQLHPRNGVIVLN

ELSSLEPEKVRVLLPLLGRARAFPERPDVPISTQLVIITTDFGREGRTRGKSLSEMRSFI

DSEFKDLYSAQSASHVRTLPFLPISLDTAAAIVRVVVREIGCTATPPLRLGIADAAVLWL

VERTKATLSVENGRAVAQETKLHVGALVERLLSEKPSDRGGADDREAPDDLARCAEPECT

VYMDDEGQVDLTC

>Lp_250009900.1

RFPRAVLALLVFAWVAGLCFELMPLQSVCAAAAAAKDADAKAMEAILELPEDDFYGVLGL

GEAGEDATESQIKSAFRKLSRRYHPDVAAGGAEDSFRQVYQRIQRAYEVLGDRRKRRVYD

ILGVDGLKKLEQPQQQQQHPLFALFGGGQAASDRGGDKELLVVVPLEDIYKGATHTFRFA

KQKICRACHGSGARSKEDLVKCPHCKGSGVLLQRVQIAPGFVQQLQQTCPHCKGKGTHVA

HLCPVCRGKAVVSGETALSVDIEEGLPEGYVLKYELEADQTPGQVPGDVLLTVITAPHAL

FKREGNDLYMNVTITLKEALLGFEKSIEHLDGHKVALRRTGVTQYAQQVRIVGEGMPRHH

VPSEHGDLYVTYNVELPKSLTPEQRELFEKNFV*

>Lp_250012100.1

MGFKTFLQGLFLASCAGVGGGVLSIRSKISDIEPLTNDEACDLRNASSLTMLAPHDQPTH

RRVYAYRLDGLMPSKHAAEQVHSKHLFRQRTSDASATSNQSDTKDHPHPLSPSFGSLYAL

HAIEAAFPFRWGWMLQTFRDTYRVSDTYNVWWRTRPAWMSGMYAMLAETPDTQQREALIH

DAAEEVERMSFVRYEALLESGDTHGAVIRGGQRPVTVTQPSAKEGGGPAETITTEMTNVA

PIYGVRFDLPSVPDKAEQEHPHPRSAPLYVLYVSAPYNSDNPTSTWEQVRMWKDGWYSRW

FAAHVVKALSEM*

>Lp_250015000.1

MKCRSLLCMKATPLLLLSAGKLSQYEQEAFEAHRRFVESQTYPGPIRAATPGDTRFYLGS

VETILQDNDRHYWRAVVDDPQVQYLVPLRIRFKTFVWVTTGWEQRMQVVQVMAHRDATIA

ELMQQVRIENQSPYLCTSAFKLSIDGRELDELKTLADYNIDEFSRIDAVEENDHLLHTEA

ERPKDWNVDEITEETLKLSPYKEMSMQPQPNLAPRYEAKPKGFYGKNNYSGMKQTS*

>Lp_260006100.1

MRRVILAALLLLGIAWPSTHAWLLGVQREHVHHYDAAKTSNSFQCLDGSKTISFSAVNDD

ICDCPDGSDEPGTPACATLRNGVVARLPDGWLFQCTNKGFSEQKISHSYVNDGICDCCDG

SDELNSGVSCPNRCAEVEGELARKMAEEKKRMQTASANKAAMRAEVAKHREELAKTLPLR

KAERDRIVAEMPALEEKNNTLQKALEPVREELRVKYAEWEATKEEREAAEEKSSCIKWRQ

TGKCKADGPREPNQDKDCGIVIPGKDSGFCECAAGNADHASQDDGTNDVADSDSNEESTI

HYEFACGHPHLTCMHVCEHNGEAAVGAVYEKPKDPNSYTTPEATAAEEALTARRSKLAEL

EKTIEESSKILNSTTLSTEELLRTLEGKTFTMEFQDYTYAVTMFKDVYQRGKGQSSGGSL

LGEWKSFAENTYAMWGKDAYDLSQMLYDRGARCWNGVTRKVDIQLVCGPENKLTHVEEPS

MCIYRMVFETPVMCDD*

>Lp_260006500.1

AVVALALLLSAMCAAHSALAGSVLEADVDRAIMFDRHFGNQEDVPYGSRAVLLRGEMVRY

TAGTSKAYQGYTVYVPCREHLTKSALQQLAGPTRATAAKGLVLELCDKDNANEEALVLFF

TTAASNLPIYFLPSSEASQQLNKVLTYAAHNTATDRVVLSIGKTVKATAPAANFTLPSAL

IESSFTHRPREARKQGAALPHVLVTAHFDSLGVAPTALTNGGASGAVAAMELWRRVTATP

TLGDDGAAAGPTPYSVTVLLGSTSRFNYAGTTSWAAQHSDDDLDRFKVVLSLDELLPVPD

AAKDATDLYLHVQDSLLKRPHGQQVVEQAEAAAKAHGLTLKIVPAKTNYQHYDLKFEHEV

FASRQMTAVTLSSHRTHHVDQLFRDVRRPPLTEADAATLAKRVDFVEAFVRALVLAAASP

KEATLTWPGSSSYIMGMLQYASESHRSPVAHHGADLRYYATAVGHQMRTQAAVAQRTASS

VATTTVDFQRLRMPGITLFSPYEEKMSVFLAKSYLVECAVAAAAFSALIAFLYVELGV

>Lp_270006300.1

MRFALSCLSGAVGAWLWRPPQADGVQCYQGVDNALTFRNKQRPGQVFSQRYERYQLGEII

PITSDVALFRFLLHDPEDEFNLKPCSTLQACYKYGVQPTEQCQRFYTPVTANHTKGYFDL

IVKRKTDGLMTNHIFGMHVGDSLLFRSVAFKIQYRPNRWKHVGMIGGGTGFTPFLQIIRH

ALTEKWDSGEVDKTKLSFLFCNRTERHILLGGVFDDLAQRFPNRFRVFYTVDLAIDKDAW

LKKNNHFLGYVTPEMVRQSMPAPDEKDKIIMLCGPDPLLALVAGTPMSTMASMSGSLNIQ

PIATDINNLVSLGGILGELGYDNNDVYRF*

>Lp_270007200.1

RHTPVSLLAAMSAAAATPSSISSASPPALPAALDFPRVKLLAVGDVGVGKSCVIKRYCEG

RFVTKYIPTIGIDFGVKKVDLSKTAVLQSSQRTVSSGCSTADSHSIPAAVRVNFWDSSGD

DDYREIRNEFYDAAQGILLFYDVRSATSFDHLQTWWEELVTYCAGLAVSVATSSADGGAA

EGKRASVAASTVTGSTAAGKAVGHNEGKAPIVVLCANKADDTVTSGAGSRTARMVSEEQG

RAWSKEHGCAGYYETSASTAMNVKEAIEDLVTQVVARFM*

>Lp_290008400.1

MSSSAVLRTVLVALVLLGTISASSAGDGRGAPITFQAEVSKMLDILVNSLYTNRAIFLRE

LISNGSDALDKIRVLYLTSPKEPLNAEGVSPTMDIRISFNEEKNELIIRDGGVGMTKDDL

ASHLGSLGTSGTKRFLEKLQEGGAAGDQNNLIGQFGVGFYSVFLVGDRVRVASKSDDSDA

QYVWESSGNGEYFLYPDPRGNTLGRGTEITIELKSDAEQFLSAETVKKAIHQYSEFINFP

IYVEEEVTVEAKKDDAAAKGEEEVLDEDAVEDEVEAAPVTEKKWTLVNENRPIWTRPIGN

VTEEEYNKFYKSFTGDYRDPLYFNHFRVEGEVEFDSILFVPAFVDPSTFSDDNALPNTNI

KLYVRRVFITDEFRDLLPRYLNFVKGIVDSNDLPLNVSREVLQESRILRVIKKKLVRKAL

TMFSDIAEQDEAIAQGKQPESPPPSGHTHLTKPTYAKFWELFGKHLRLGVMLDSNNRNRL

TKLFRYKSSKSEGAYIGLQTYVDRMKKGQKGIYYISGDSVERIKKSPVLEDAVNHDVEVI

FMTDAIDEYVVAQMTDFAGKKLINLAKEGVQFEETDARQRVIDKKRKEKYEAFFTRLHAL

FGYSEVRKIILTKRLTNEAFILSSGENQITARLANIMRGQSMSLVDQQTTAERVLEVNYR

HPLVDEMFKRFAVNEDDEVAIDVAWVLYDTANLQAEFPVADVAAYSRRINRLLRSSVDLP

ADDTLLPPDDAEYAVSNTEAAEEEGDGHDGAAEENADADAQEDDEDL*

>Lp_300018200.1

MKAGGLAALLAQAVTAGDVNACCGVLRRIFSETMLDDGVPLATRGAWVTARGTPHRSSPL

SSSSSFSSTSSSTSPAASGLAARAGRLSKVLRAPAQASDSIQTTSLHAAAASAAAPHLSS

SVHNDVVNFARYAQRQRAQANSFANAAAQAVLRTTFPQPENKGLSPGSKAAQTSPFTVSS

PTTPTSSPLSHPLPSSVNASDAKSLIQRVWALATASLKDNPSLVKTSSVQVLCHTAQRTG

FWREALHFSEHLPRPPAPLFLSSLLRPQNVQPVLAHCARHGWALDVTNAIRVLAEEHGSW

SAALAVAQDAEHRFPYGEVYSLGVLIPYLAASGDAAQARQLFESGVAQGALVDATLVQHL

IMQTAALKQWETCLYMLQCLYQTQETAQLLPTDVDFFRQLMELSPSWVTSLRLLHIARAS

EVKPDERTVAILLTQCDKAGAWREAAAVYDMAVREDFIDSLAAGSNYHTLVRSFSAMQQW

QKALEALSWMSKAGDASLTAGMSELVTLCEQAGQWEAALTVGASLMESGTHLFSAQTSMA

LLFACAKGAQWSFATQLFMAQLHDVRVDTHPLALCAVMQACVAARQWRAALHAYHEAQAS

QPRIVVPPLAHRMAVKACVLGQQWSQAIAVLDTMRQDGLPLDNNSQRLGLWAAALQGQWE

LSLAFLQGIPRRSRTPQDRMVARNAARAVSPAVDALTLRLLQSR*

>Lp_320018200.1

MLRRAFTPAACATSLLGSVVGVNAEAPLPCFAVTHTTTGRTGLVMHTPKADIQLGRFNFM

EQYNEFRNRSGRVPDYYVEKYRDTYWRVDEYIRRSENYIQSQGFFYLPTLEFPWYKGCMP

LLSSYQIRVHYGRHHRAYVEKLNQLIEGTDMYGMQLDSIIVKSANDDRLRGVYNNAAQHY

NHCFYWKCIQPYGSNIPPDLSAAVSAQYGSVENFEKQFITAAQNLFGSGWVYWIYDRKAG

KFDIMSYSNAGCPLTNYDHVPLLCVDVWEHAYYIDYENKRPEYLAKYFDVVDWHWAERHW

KRATGQEYHEMKFW*

>Lp_320021100.1

MGVFRLLLLLLCAAVVLINVVPDGNTVEPLRDLLLDFDQEALKTRFGSTRSLDHAGTRSV

YIKVLSEAENRIFNSLEKPDQRAFTCSLMRSEARRYARSRDGSYGGHLTDAVLQLRDSYV

HGLRYLPLAVKKDIQDSFALQRPTLYHVALVVKETFSCLAPALSSGRCPSYAFLREVRGK

TDDEILGSCTKANSAYN

>Lp_320023900.1

MSRLSPHACVLTRVTVCVSTLLLLCVAVESHAFFGGGNFKGNAPPPEVQHRPEVDYYQIL

NLADKREAATDKEIRREFRRLSRLYHPDVANTEDDKIKYSEVNRAYEVLSDKRKRKVYDM

RGERGLEQLEQLDRAKDQPGGGMNPFTQFMGMHVDDGLRGPNMEVETKVELSQVFTGAQV

PLQIHKHKVCHRCKGTGADNTAPMVPCSQCGGKGVLHQQIQLFPGMIHNVNQPCPSCGGS

GKRPERLCPVCNGKKVVYGSSTVVLELEPGMEEDHVLKFEMEAEETPDRLPGDLLVHVRT

HPHPVFARRRNQLDLDASLVLTLKEALLGFDRHITHLDGEEQVEVRRETPSPYGTVMRVA

GKGMPKLHVPSERGDLYVHLHYDLPERLTEEQKALVEKLL*

>Lp_320024500.1

MRSAAQPLSPLLLAASHAQSWADVLRAYSKCRMYLHNCYQPSTAELQFGLSRMDSAWSLS

LFYYDFIKSPRTRAPPDASLVVMMIQRCKELTFVKGLKRVIEEDVSSSTIEGAKGKITLA

SYTGMWEVALATLMSQPKLQQSLPLRRTVLATLSAHNQWRRALEVLRMKPALELRPTVVR

PLVRCFGRLELHEMALRLTAASLAAGYAFDAALLSALLTTLQDTNQWSAALEVAQRMQLL

SATRAEGRKNVVLFNQLVSCLYEANLYGDYTLEEVVLDVLSRSNPRDTATIWREARQKQF

RLRLHSEVFQHFQGVLLPLSQLYSKIMRIPRWYSRSISHIVTAAVKDNTVVLVMDTNFLL

QLVHKNLSPEHFYAYIKQQYPDIHAYSYATIVVPFTVLQEAHMLIWSTRARMSMSTRTLL

WSRVVAIVEQPHVYALSLAGEYPSTSLGIIPKLAYAKMPQNVAGSFEHDPDLRILNVCVS

LQHYLRHAKITENLGGLPPVEGIALFAFLKYHVRRYSNTVKGCSVDRLLLCTMDRRMAHA

AEELGIRTFPHISNAP*

>Lp_350018800.1

MADLLLGLALGQVAAMALPVLANGLLSVKRPPRVNISQERRNVHTCRPVIPYRAPVPYTG

GRVRALFIGINYTGTGNALRGCVNDVRSMLGTLQRIQFPISECCILVDDPKFPNFTAMPT

RANIIKYMLWLTSDLRPGDVLFLHYSGHGGQTRAKHDSEEAYDQTMIPLDHQESGSILDD

DLFLMLVAPLPQGVRMTCVFDCCHSATMLDLPFSYVAPRQSGTREQMRQVRRDNFSNGDV

IMFSGCTDEGTSADVHNNSGTANGAATLAFTWSLLHTQGFNYLNILLKTREELRKQGREQ

VPQLTSSKPIDLYKPFSLFGMLTVNQAMMTHCVPQQYQRPPSNIPPQAMPPQLGGYPAYM

PPPPQGFPQRGDPNDGNRGAGYNNGNSNGARGIPQPQYTPQPQYGGGRPQYPPPPNVQYH

GGPPPPSWGGNPPAQYSYNPNPPL*

>Lp_350038000.1

MRRLFASVSVAAAATLRCYATKFYTDSHEWVEQADGDAIIGISTYAQENLGDVVYVSLPQ

VGDKVSAKDVVGEVESVKATSNVYSPVDGTVTAVNEKLKDEPGLINQSPEDKGWLVKMKC

TEIPKGLMDAEAYKKFLE*

>Lp_360006200.1

MVELFRPVALVVIAVTCAIAAVGENAEGLYSGIWDVSTLVPGRGSVPPIHELHVEMAHWA

LRERNFSRVSRLLSSAAERLGVIYTAVPAAETTLCHAAPSLHWEWEATRKDDGLLVDCLP

GQWLTPAALRTFEHDGFLHLRFDRPQPLSVMSNGACVATSWSLHDVALTAVTEDSVVVSG

TLSDSTTGSPATCKAYFELTRRPMATAKEMSPSSGVYTPIVMLLVVIAVRFLPRYVLTHI

GQVDSSSYRGRNPGKLSSARRAELLQKQRRIIQQMKAQDLANGPKEKTS*

>Lp_360008900.1

MKLTLQKILFIALPAVISTMVSSLLILGCTELWETQTSNSVFAQMCGQAGPCVRITSGPI

DRLVVDHLDFEHDTVGDVIKRIPAWFGSQARPLGLTVHNLLLKEDVPLRHQGLSALSHIS

LVSMKEMRLEKEPLTVRVGVASK*

>Lp_360023400.1

MYRFRAVPKLATLAALRFLTITPIPMPALSPTMEKGKITEWCKHPGDPIATGDTWCKVET

DKAVVSYDNATEEGFFARVIVPAGEETVVGQTVCLMVDEKEGVNSDEVKNWKPEGEEAAA

PAAEEAAPAATPAAAAPAAAAPAAPAAASGDRVKASPYARKLAAEKNVSLSSIKGTGGGV

GRITSKDVEAAVKSGAAAAPTKAPAAAAAAAAAPAAKAAAPSAPAKGTPPANPNYVDIPT

TTMRAVIAKRLHQSKNLDIPHYYLFDTCRADNMMALIKQLNAKGNGEYKITVNDYIIKAV

ARANVLVPEVNSSWQEGFIRQYSTVDVSVAVATPTGLITPIIRNAQAKGLVEISKEMKAL

AKKAREGTLQPNEFQGGTCSVSNLGATGIPNFTAIINPPQAMILAIGSAKPQAEIIKNEE

TGEYELTGKVENVVNFTASFDHRVIDGALGAKWFQGFHDAIENPLSLLL*

>Lp_360036400.1

MSLLALNVCLISSGFDTVALPLSSTLCPVDPDDDLDEGTFGYTLAPLTEKKARRNYGYAH

CTVVQLCIRALDLKTVREIVRDTWVKFRERVTEAKESLILDGIADGPVFAKTDDGQTVRL

PNIKVERSTELMWLHETLVRELEPYHVKVASTDVAKRSFNTKFPADSNTTTAEWLLSFLT

KSSHENYVPHISLGACTMENVAALSYLQRTEVPWRECRLVVSHMGNYCSCFDLME*

>Lp_360040300.1

MQKTWSRLPSWAAVILSVVCFAQLQRFAAASVDLSDTLQRNHVLYFTEHGDHTAPVDGGD

AASANTRYVQLSNGSRFVCETTDIKHHHPPSFADSLLNMRMQSIIGSIKAAGPPCTHVVD

EQRSIVLCWNREIRVDKVAARTSRILGRRNETSPPDYWTGHDAFGRYVATRYDEGEECWY

TRRPSETEIRFYCRYTEMENPVPFLTLHESAQCHFMLRVMSDRFCFVPQLDHAVETETVH

CYMLE*
